# Genetic glyco-profiling and rewiring of insulated flagellin glycosylation pathways

**DOI:** 10.1101/2022.03.25.485807

**Authors:** Nicolas Kint, Thomas Dubois, Patrick H. Viollier

**Affiliations:** Department of Microbiology & Molecular Medicine and Geneva Center for Inflammation Research (GCIR), Faculty of Medicine, University of Geneva, Rue Michel Servet 1, 1211 Genève 4, Switzerland; University of Lille, CNRS, INRAE, Centrale Lille, UMR 8207-UMET-Unité Matériaux et Transformations, F-59000 Lille, France

## Abstract

Glycosylation of surface structures diversifies cells chemically and physically. Sialic acids commonly serve as glycosyl donors, particularly pseudaminic (Pse) or legionaminic acid (Leg) that prominently decorate eubacterial and archaeal surface layers or appendages. We investigated a new class of FlmG protein glycosyltransferases that modify flagellin, the structural subunit of the flagellar filament. Functional insulation of orthologous Pse and Leg biosynthesis pathways accounted for the flagellin glycosylation specificity and motility conferred by the cognate FlmG in the α-proteobacteria *Caulobacter crescentus* and *Brevundimonas subvibrioides*, respectively. Exploiting these functions, we conducted genetic glyco-profiling to classify Pse or Leg biosynthesis pathways and we used heterologous reconstitution experiments to unearth a signature determinant of Leg biosynthesis in eubacteria and archaea. These findings and our chimeric FlmG analyses reveal two modular determinants that govern flagellin glycosyltransferase specificity: a glycosyltransferase domain that accepts either Leg or Pse and that uses specialized flagellin-binding domain to identify the substrate.

## INTRODUCTION

Sialic acids, also known are nonulosonic acids (NulO), are nine-carbon (α-keto) acidic sugars featuring acetamido linkages that are found in all domains of life [1]. The most prevalent vertebrate sialic acid, (5-)N-acetylneuraminic acid (Neu), occurs on surface glyco-conjugates like glycolipids or glycoproteins [2, 3]. While meningitis-causing eubacteria also camouflage their surface with Neu, most eubacteria and the archaea typically decorate their cell surface structures with (5-, 7-)di-acetamido derivatives, either pseudaminic acid (Pse) and/or its stereoisomer legionaminic acid (Leg, Figure 1). Pse or Leg are constituents of capsular polysaccharides (CPS or K-antigen)[4] or the O-antigen of lipopolysaccharide (LPS)[5], but they often also occur conjugated to proteinaceous surface appendages, for example on the subunits of S-layer arrays [6], pilus adhesins [7] or flagellar filaments (the H-antigen)[8, 9]. Pse and Leg derivatives synthesized *in vitro* can be added exogenously in metabolic labeling experiments to be incorporated into bacterial surface structures [10, 11]. Moreover, Pse and Leg are attractive vaccine targets as shown by the recent report that mice immunized with Pse chemically conjugated to a carrier protein were protected against the Pse-containing pathogenic *Acinetobacter baumannii* strain Ab-00.191 [12].

**Figure 1.**
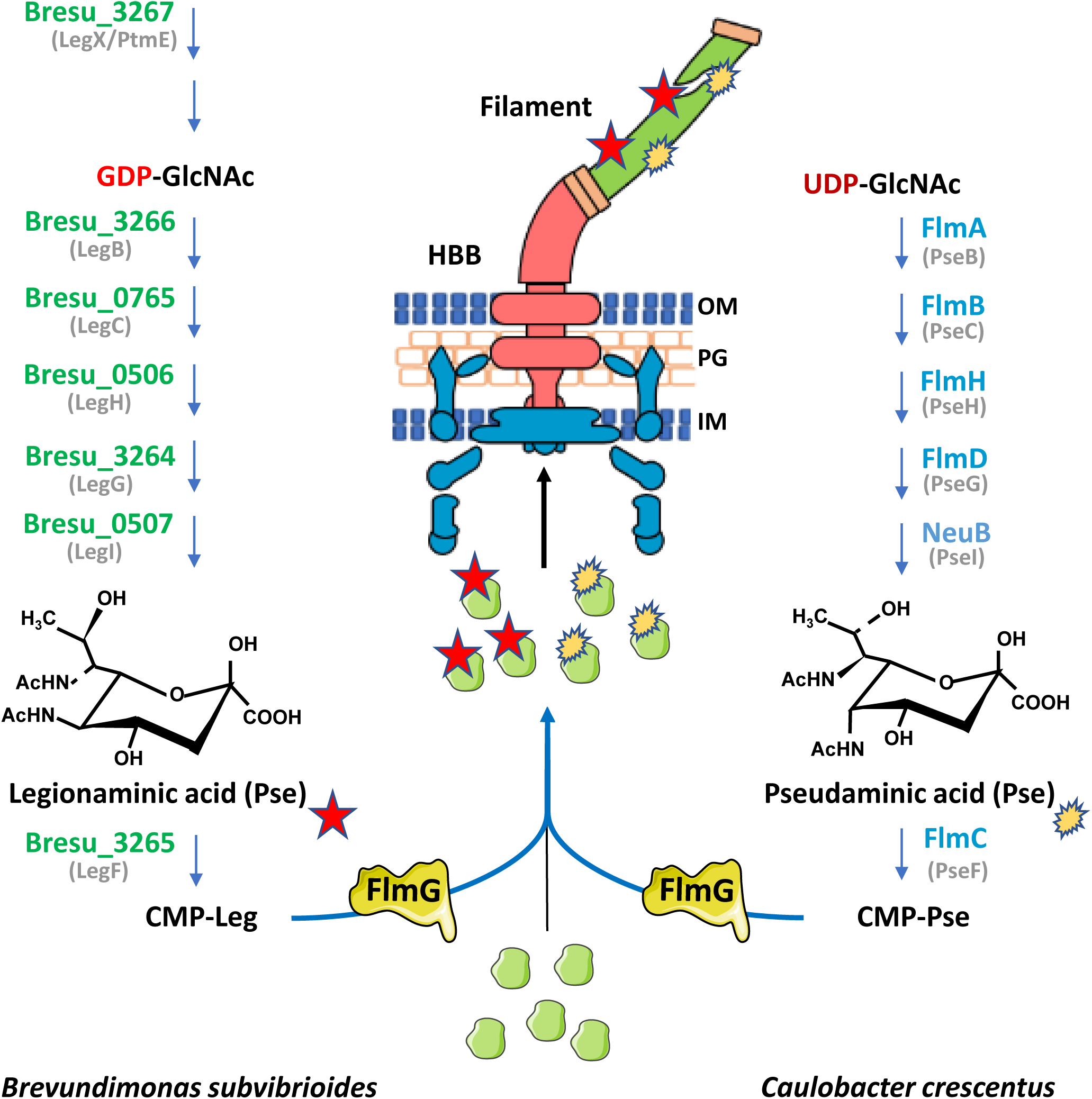
Model of flagellin glycosylation pathway in *B. subvibrioides* (*B*.*s*.) and *C. crescentus* (*C*.*c*). Schematic of the *B*.*s*. and *C*.*c*. flagellum with the MS- and C-ring structures (in blue) inserted in the inner membrane (IM), the hook basal-body (HBB, in red) components spanning the periplasm with peptidoglycan layer (PG) and outer membrane (OM) and the flagellar filament (in green). The legionaminic acid (Leg, 5,7-diacetamido-3,5,7,9-tetradeoxy-d-glycero-d-galacto-non-2-ulosonic acid, structure shown on the left, red star) and pseudaminic acid (Pse, 5,7-diacetamido-3,5,7,9-tetradeoxy-l-glycero-l-manno-non-2-ulosonic acid, structure shown on the right, yellow shape) biosynthesis pathways that are present in *B*.*s*. and *C*.*c*., respectively, are shown with the (predicted) sequential enzymatic steps. Shown in parenthesis are the respective enzymes that perform the equivalent reaction in *Campylobacter jejuni*. Activated legionaminic acid (CMP-Leg) and pseudaminic acid (CMP-Pse) are transferred to the flagellins subunits (green shape) by FlmG prior to flagellin secretion through the flagellar secretion apparatus and their subsequent assembly into a flagellar filament on the cell surface.

Pse- or Leg-decorated flagella may also be immunogenic. The flagellum consists of three major parts: an envelope-embedded basal body that houses the rotary engine and the secretion apparatus, a universal joint known as the hook that transmits torque from the motor and that protrudes to the cell surface, and finally a tubular flagellar propeller composed of flagellins that is mounted on the hook (Figure 1)[13, 14]. Once flagellin subunits are translated, they are exported through the flagellar protein secretory apparatus along the hollow flagellar filament for polymerization at its growing tip. Glycosylation typically occurs post-translationally on serine or threonine residues of flagellin by highly specific and flagellin glycosyltransferases (fGTs)[15, 16]. Unlike the pilus-specific glycosyltransferases that execute the glycosylation only after the acceptor protein has been translocated across the membrane [17, 18], the fGTs are soluble enzymes that act on flagellin in the cytoplasm before their secretion through the flagellar apparatus. Two types of fGTs have been described to date, the Maf- and FlmG-type [15]. It is thought that these fGTs accept CMP-activated forms of Pse or Leg as glycosyl donors and then join the Pse or Leg moiety to the flagellin acceptor molecule that they bind directly. Inactivation of the fGT or the corresponding Pse-/Leg-biosynthesis pathway results in failure to modify flagellin and (often) a motility defect [19]. It remains mysterious why these flagellin glycosylation mutants are non-motile and flagellin is often poorly secreted. Such mutants harbor a hook-basal-body (HBB), yet they lack a flagellar filament [19].

The synthesis of CMP-Pse or CMP-Leg proceeds enzymatically by series of steps [20-22], ultimately ending with the condensation of an activated 6-carbon monosaccharide (typically N-acetyl-glucosamine, GlcNAc) with 3-carbon pyruvate (such as phosphoenolpyruvate, PEP) by Pse or Leg synthase paralogs, PseI or LegI, respectively (Figure 1)[23, 24], whereas the sialic acid Neu is synthesized by the NeuB paralog [20-22, 25]. A major difference between the Pse and Leg pathways is that the former uses UDP-GlcNAc as starting material whereas the latter usually builds on GDP-GlcNAc [21, 22]. Pse or Leg must first be activated with CMP by the PseF or LegF enzyme, respectively (Figure 1) for used as glycosyl donors by terminal glycosyltransferases, including fGTs.

The structure-function relationship and specificities of Maf and FlmG fGTs is poorly understood. Some Maf enzymes have been linked to flagella glycosylated with Pse, while other Maf affect modification of flagella with Leg [10, 26-28]. The determinants conferring donor or acceptor specificities in these enzymes have not been elucidated. Recently, FlmG and Pse biosynthesis enzymes from the Gram-negative α-proteobacterium *Caulobacter crescentus* were shown to be necessary and sufficient for modification of flagellin [19]. *C. crescentus* encodes six flagellin paralogs [29-31] that are no longer modified in the absence of Pse or FlmG [19]. Conversely, expression of FlmG and the FljK flagellin in heterologous hosts producing Pse resulted in FljK modification [19]. Sequence analysis predicts a simple 2-domain organization for FlmG: an N-terminal tetratrico-peptide-repeat (TPR) domain and a C-terminal GT-B type glycosyltransferase domain [15]. Bacterial-two-hybrid (BACTH) assays revealed that the TPR domain can directly bind flagellin, whereas the GT-B domain cannot [19]. The donor specificity of the GT-B domain remains unexplored in the absence of a FlmG system that links Leg to flagellin.

Here we establish a glyco-profiling platform for functional analysis of Pse and Leg biosynthesis pathways using motility as a proxy and we exploit this set-up to uncover a novel FlmG glycosylation system in *Brevundimonas subvibroides* that modifies flagellin with Leg. Using the *B. subvibroides* and *C. crescentus* Leg and Pse biosynthesis mutants, we show that the two pathways are genetically insulated, defining a first level of specificity. We then reconstitute flagellin glycosylation using the *B. subvibroides* components in *C. crescentus* and we reprogram a Pse-dependent FlmG into a Leg-dependent enzyme through domain substitutions in chimeras. Thus, two modular determinants govern specificity in fGTs, with the GT selecting either Leg or Pse as donor and linking it to the correct acceptor identified through a flagellin-binding domain.

## RESULTS

### Genetic glyco-profiling in *C. crescentus* Δ*pseI* cells using motility as proxy

Phylogenomic and functional analyses show that the genes encoding PEP-dependent synthases of sialic acids are wide-spread, present in all domains of life. The PseI and LegI synthases predominate in the eubacterial and archaeal lineages, sometimes co-encoded in the same genomes. As a rare example, *Campylobacter jejuni* 11168 has three (NeuB, PseI and LegI) synthases [22], while the *Pseudomonas sp*. Irchel 3E13 genome (NZ_FYDX01000009.1)[32] encodes two predicted synthases, a PseI and LegI homolog, and *C. crescentus* only encodes only PseI (previously called NeuB)[19, 33]. Our previous heterologous complementation experiments of the motility defect associated with *C. crescentus* Δ*pseI* cells showed that of the three *C. jejuni* 11168 synthases, only PseI could support motility in *C. crescentus* [19]. These experiments provided strong support for the notion that the Pse synthesis pathway can only function properly with PseI, but not when it is substituted with LegI or NeuB. However, it is known that Pse and Leg often occur in derivatized (modified) forms [1, 3]. Such modifications could occur before the PseI synthase acts or afterwards. In the latter case, most (if not all) synthases would be predicted to produce the same Pse molecule, which is then derivatized once it has been synthesized. If so, then the protein executing a particular enzymatic reaction should be replaceable by an orthologous enzyme executing the same reaction.

To investigate this idea on a comprehensive scale, we individually cloned 21 synthetic (codon-optimized) PseI or LegI coding sequences (CDSs) onto an expression plasmid for genetic glyco-profiling experiments using motility as proxy to report the ability of the candidates to substitute for the endogenous PseI of *C. crescentus* (Figure 2A and S1). In support of the notion that derivatization occurs after the PEP-dependent condensation reaction to form Pse or Leg, our glyco-profiling analysis revealed that putative PseI proteins (identified by sequence comparisons to *C. jejuni* 11168, Table S1) conferred motility to *C. crescentus* Δ*pseI* cells, whereas putative LegI synthases did not. This stringency for PseI synthase function using the *C. crescentus* motility readout was not only observed across species (e.g. *Shewanella oneidensis* vs. *Shewanella japonica*) or class (*Shewanella japonica* vs. *Magnetospirillum magneticum*), but also across the Gram-negative / Gram-positive divide (e.g. *Pseudomonas sp*. Irchel 3A5 vs. *Kurthia sibirica*) and, remarkably, across kingdoms (e.g. *Leptospira interrogans* vs *Methanobrevibacter smithii*, see Figure 2 and S1). Strikingly, in the case of certain *A. baumannii* strains, only one synthase least was able to confer motility to *C. crescentus* Δ*pseI* cells, suggesting that it is a PseI ortholog, while other two genomes might encode LegI-type synthases (see below).

**Figure 2.**
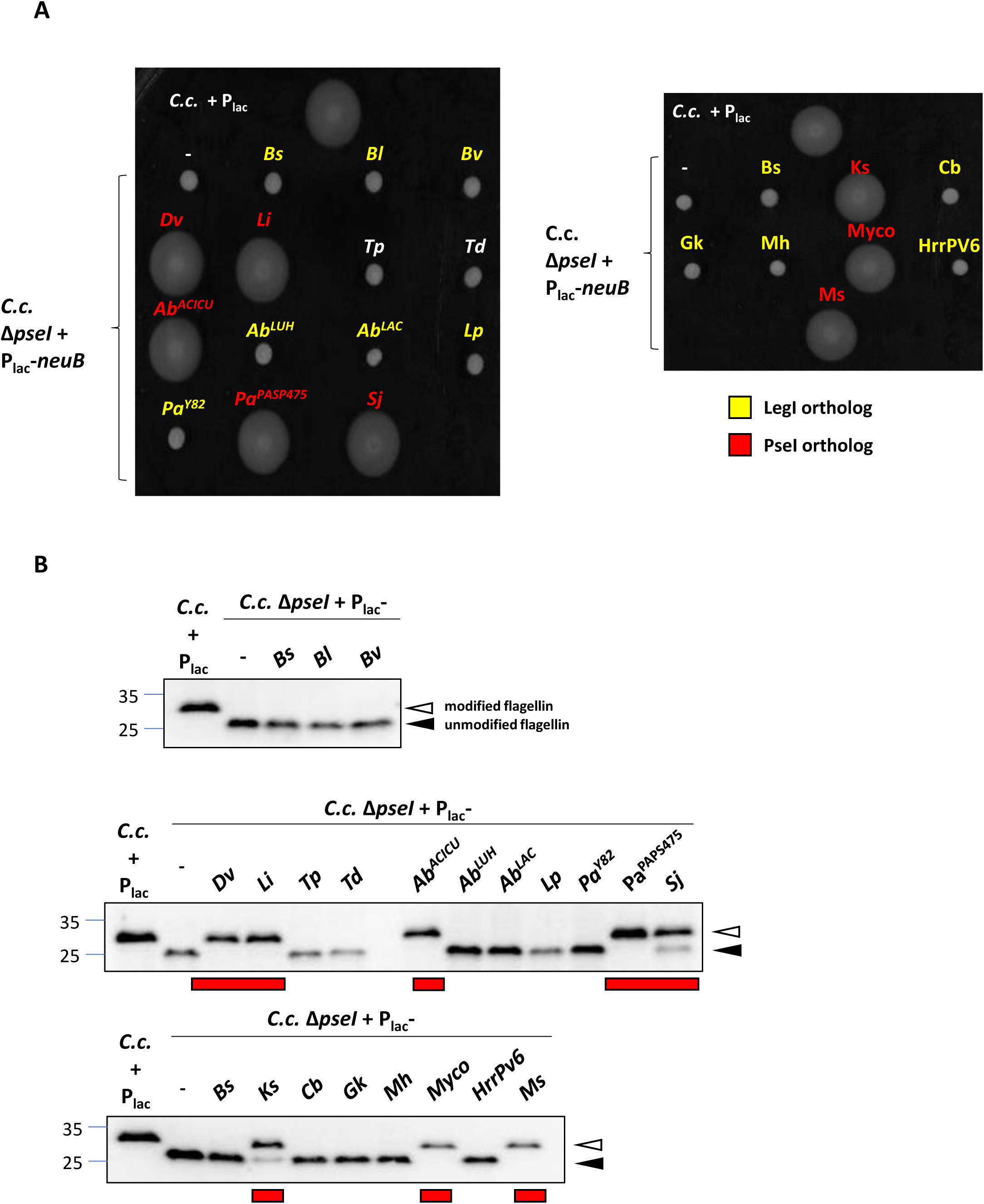
Heterologous complementation of *C. crescentus* Δ*pseI* cells with candidate pseudaminic acid synthase (PseI) enzymes and legionaminic acid synthase (LegI) enzymes. (A) Motility assay of *C*.*c*. Δ*pseI* cells expressing predicted Leg or Pse synthases (LegI or PseI, in yellow or red, respectively) from P_*lac*_ on pSRK-Gm. Overnight cultures were spotted on PYE soft (0.3%) agar plates with gentamycin and IPTG (0.5 mM) and incubated for 3 days at 30°C. Only predicted PseI-like synthase coding sequences (CDSs) restore motility of the *C*.*c*. Δ*pseI* cells. Note that the predicted synthases labeled in white do not share similarities with LegI or PseI of *C. jejuni*. (B) Immunoblot of cell extracts from cells in (A) probed with antibodies to *C. crescentus* FljK (FljK^Cc^). The migration of flagellins in Δ*pseI* cells is shifted towards lower molecular mass suggesting that post-translational modification of flagellin is defective in Δ*pseI* mutant. Molecular sizes are indicated by the blues lines (in kDa). Bl: *Brevundimonas lutea*, Bv: *Brevundimonas viscosa*, Dv: *Dermabacter vaginalis*, Li: *Leptospira interrogans*, Tp: *Treponema pallidum*, Td: *Treponema denticola*, Ab^ACICU^: *Acinetobacter baumannii* ACICU, Ab^LUH^: *Acinetobacter baumannii* LUH, Ab^LAC^: *Acinetobacter baumannii* LAC-4, Lp: *Legionella pneumophila*, Pa^Y82^: *Pseudomonas aeruginosa* Y82, Pa^PAPS475^: *Pseudomonas aeruginosa* PAPS475, Sj: *Shewanella japonicum*, Basu: *Bacillus subtilis*, Ks: *Kurthia sibirica*, Cb: *Clostridium botulinum*, Gk: *Geobacillus kaustophilus*, Mh: *Moorella humiferrea*, Myco: *Mycobacterium sp. KS0706*, HrrPV6: *Halorubrum sp. PV6*, Ms: *Methanobrevibacter smithii*. All the genes come from synthetic fragments codon optimized for *E. coli* (except the synthase CDS from Gk that is codon optimized for *C*.*c*.). Empty carets indicate the position of modified (glycosylated) flagellin, whereas filled carets mark unmodified flagellin. Red boxes indicate PseI functional orthologs.

Immunoblotting with antibodies to *C. crescentus* FljK [19] (FljK^*Cc*^, Figure 2B) revealed that all the PseI-type synthases that restored motility, also restored FljK modification. By contrast, the non-orthologous synthases neither supported motility, nor flagellin glycosylation. We conclude from our survey that (heterologous) PseI synthase activity generally confers motility to *C. crescentus* Δ*pseI* cells, whereas LegI-type (or NeuB-type) synthases are unable to do so.

### Flagellin glycosylation in *Brevundimonas subvibrioides* is FlmG- and LegI-dependent

An unexpected glyco-profiling result was that the synthase orthologs encoded in the genomes of different *Brevundimonas* species, that are members of the same family (*Caulobacteraceae*) as *C. crescentus*, were unable to replace PseI (Figure 2). A closer look by sequence comparisons revealed that three *Brevundimonas* orthologs tested are in fact more similar to LegI from *C. jejuni* 11168 than to PseI. For example, the *B. subvibrioides* ortholog is 42% identical and 64% similar to *C. jejuni* 11168 LegI and only 32% identical and 52% similar to PseI (Table S1). On this basis, we speculated that these *Brevundimonas* species likely synthesize Leg rather than Pse. In support of this idea, our bioinformatic searches using *C. jejuni* 11168 as reference genome identified all six putative enzymes in the *B. subvibrioides* ATCC15264 genome (CP002102.1) predicted to execute the synthesis of Leg from GDP-GlcNAc. Importantly, *B. subvibrioides* also encodes a FlmG ortholog (43 % identity and 59% similarity to *C. crescentus* FlmG), raising the possibility that it uses FlmG to glycosylate its flagellins as *C. crescentus*. Yet, no obvious sequence homologs of the six Pse biosynthesis enzymes were found by BlastP searches, whereas orthologs of Leg biosynthesis enzymes are readily discernible. Thus, we reasoned that *B. subvibrioides* FlmG could be a Leg-specific flagellin glycosyltransferase, rather than a Pse-dependent enzyme as for *C. crescentus* [19].

To test this idea, we first confirmed that sugar modifications are indeed present on *B. subvibrioides* and *C. crescentus* flagella. For *C. crescentus*, flagellin glycosylation by Pse was inferred, but not yet chemically proven. We purified flagella from supernatants of *B. subvibrioides* and *C. crescentus* cultures by ultracentrifugation, dissociated covalently linked sugars by acid-hydrolysis, derivatized them and then analyzed the liberated material by HPLC (Figure S2A). A Pse-like molecule was extracted from *C. crescentus* flagella, having a retention time (9.8 minutes) that is nearly identical to that (9. 7 minutes) of a Pse standard (harboring a triple acetamido modification, Pse4Ac5Ac7Ac) isolated from an *A. baumannii* capsule [34]. Co-injection of this Pse-standard along with the material extracted from *C. crescentus* flagella, revealed a co-eluting peak at 9. 7 minutes of double intensity compared to that of the standard (Figure S2A). When the same procedure was used to liberate a derivatized nonulosonic acid from *B. subvibrioides* flagella, a major peak was detected by HPLC analysis having a retention time of 9.8 minutes, along with a minor one eluting at 15.3 minutes (Figure S2B). A known Leg standard with a double acetamide modification (Leg5Ac7Ac) isolated from a different *A. baumannii* capsule [35] eluted at 12.3 minutes, suggesting that *B. subvibrioides* flagella are modified with a Leg-derivative that is distinct from Leg5Ac7Ac. Indeed, Leg derivatives of different mass or just simply epimers are known with substitutions of the N-acetyl/acetamido groups at the C-5 and C-7 positions, such as N-acetimidoyl or acetamidino, N-formyl and N-hydroxybutyryl groups [1, 3], that are synthesized from a Leg-type biosynthesis pathway requiring LegI.

To determine if the gene predicted to encode the LegI-like synthase of *B. subvibrioides* (Bresu_0507, henceforth LegI^*Bs*^) or the FlmG ortholog (Bresu_2406, FlmG^*Bs*^) are also required for motility in *B. subvibrioides* ATCC15264, we engineered in-frame deletions in each gene. We then probed the resulting Δ*legI*^*Bs*^ and Δ*flmG*^*Bs*^ single mutants for motility defects in soft agar and analyzed flagellin glycosylation by immunoblotting using antibodies to FljK^*Cc*^ (Figure 3A-3D). Both mutants showed strongly reduced motility on soft agar and increased migration of flagellin through SDS-PAGE compared to *WT*. While no difference in the abundance of flagellin was observed in extracts from mutant versus *WT* cells, flagellin was barely detectable in the supernatants of mutant cultures, suggesting flagellar filament formation is defective in these mutants. Moreover, transmission electron microscopy (TEM, Figure 3E) revealed substantially shorter flagella on both mutants (average length 1 or 1.2 μm, Figure 3F) compared to those on *WT* cells (4 μm), suggesting that LegI^*Bs*^ and FlmG^*Bs*^ govern flagellin glycosylation and export (or stability after export). However, we cannot rule out that LegI^*Bs*^ and FlmG^*Bs*^ also promote filament assembly in addition to flagellin secretion. Flagellins are exported before their assembly into the filament [13, 14, 36], but when the assembly step is blocked they typically accumulate in the supernatant. In this situation, the resulting cells feature only a hook on the surface lacking the filament or possibly a very short stubby filament, similar to the ones revealed in our TEM images.

**Figure 3.**
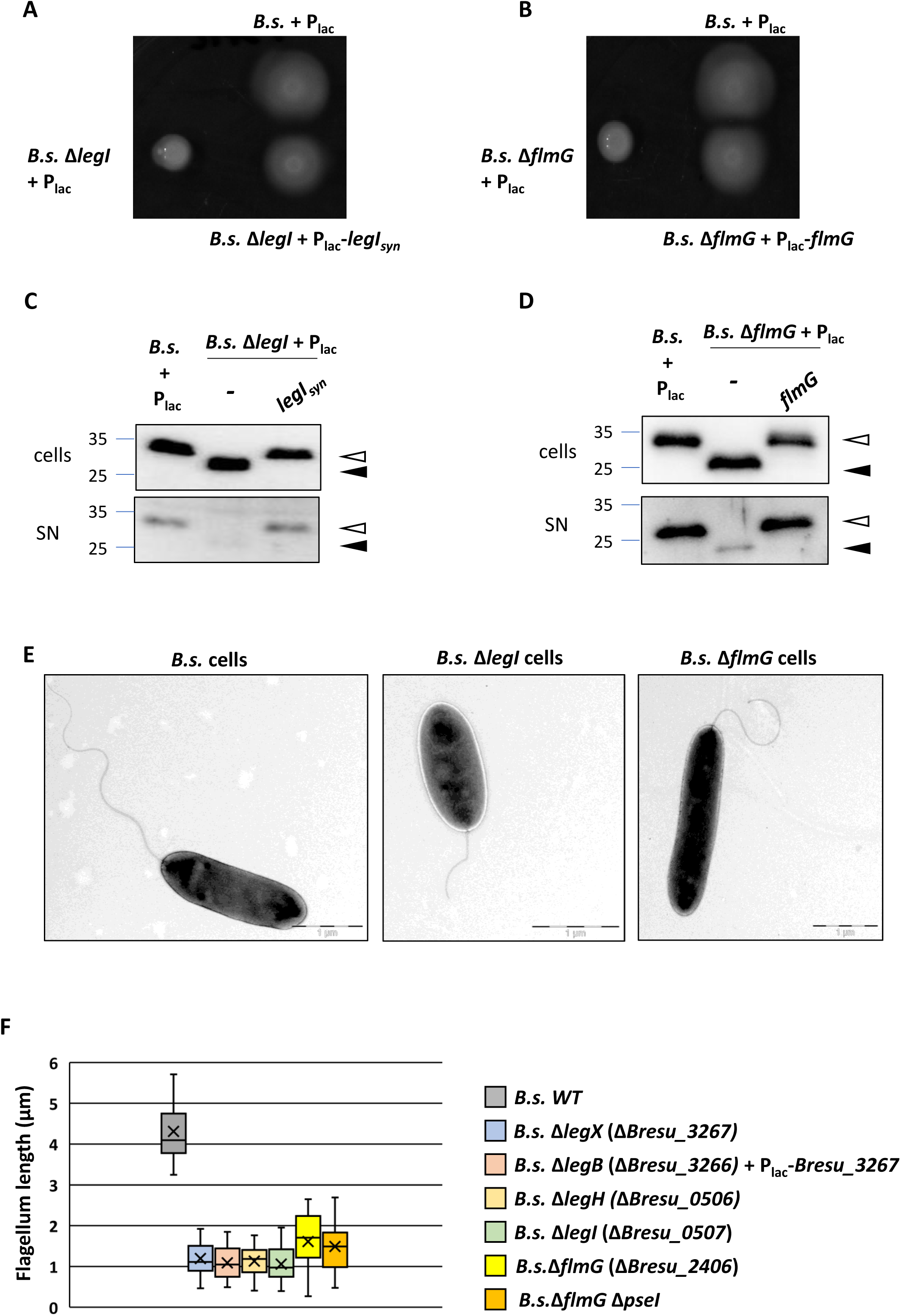
The LegI synthase ortholog and the FlmG glycosyltransferase ortholog are important for motility, flagellin glycosylation and secretion in *B. subvibrioides*. (A) and (B) Motility assays of *WT*, Δ*legI* (A) and Δ*flmG* (B) *B*.*s*. cells. Overnight cultures of WT and mutants harboring the empty pSRK-Gm vector (+P_*lac*_) or the corresponding complementing plasmid were spotted on PYE soft agar plates with gentamycin and IPTG (0.5 mM) and incubated for 7 days at 30°C. (C) and (D) Immunoblots of cell extracts (cells) and supernatants (SN) of cultures from *WT* and mutant cells probed with antibodies to FljK^Cc^. The estimated molecular mass (in kDa) are indicated by the blue lines on the left. Empty carets indicate the position of modified (glycosylated) flagellin, whereas filled carets mark unmodified flagellin. (E) Transmission electron microscopy (TEM) analyses of negatively stained *WT*, Δ*legI* and Δ*flmG B*.*s* cells. (F) Flagellum length measurements determined by TEM of *WT* and mutants *B*.*s*. cell: *WT*, Δ*legX* (Δ*Bresu_3267*), Δ*legB* (Δ*Bresu_3266*) expressing *legX* in *trans* from P_lac_ on a plasmid, Δ*legH* (Δ*Bresu_0506*), Δ*legI* (Δ*Bresu_0507*), Δ*flmG* (Δ*Bresu_2406*) and Δ*legI* Δ*flmG*. Box plots represent the distribution of the flagellum lengths and the cross indicates the average length. Twenty-five flagella were measured in each case from TEM images.

The impaired flagellar filament assembly observed in our mutants are clearly due the absence of LegI^*Bs*^ or FlmG^*Bs*^ as shown by the fact that introduction of a plasmid harboring either *legI*^*Bs*^ or *flmG*^*Bs*^ under control of the IPTG-inducible P_*lac*_ promoter into the corresponding mutants, restored motility as well as flagellin modification and export, whereas the empty vector (pSRK-Gm [37]) was unable to do so (Figure 3A-3D). Having confirmed the importance of LegI^*Bs*^ and FlmG^*Bs*^ in flagellin modification/secretion and motility in complementation experiments, we asked if *C. crescentus* FlmG (FlmG^*Cc*^) can substitute for FlmG^*Bs*^ and vice versa. These heterologous complementation experiments revealed that the FlmG variants are not interchangeable between *C. crescentus* and *B. subvibrioides* (Figure S3A), whereas the PseI^*Cc*^ substitution experiments described above showed that PseI orthologs are functionally interchangeable (Figure 2). To test if such heterologous complementation is also possible for Leg synthases using the motility defect of *B. subvibrioides* Δ*legI*^*Bs*^ cells as proxy, we conducted the orthologous glyco-profiling for LegI orthologs expressed from pSRK-Gm plasmids as described above (Figure 4). Strikingly, we obtained a near mirror-image of the complementation results from the *C. crescentus* Δ*pseI*^*Cc*^ glyco-profiling: the orthologs that were unable to restore motility and flagellin glycosylation to *C. crescentus* Δ*pseI*^*Cc*^ cells, predominantly restored motility (Figure 4A) and flagellin glycosylation (Figure 4B) to *B. subvibrioides* Δ*legI*^*Bs*^ cells. Since these complementing synthases exhibit greater overall sequence similarity to LegI than Pse of *C. jejuni* 11168 (Table S1), we concluded that *B. subvibrioides* indeed encodes a Leg-dependent flagellin glycosylation pathway. Thus, while the *C. crescentus* and *B. subvibrioides* flagellin glycosylation systems are clearly evolutionarily related, they diverged to exhibit dissimilar donor and acceptor specificities.

**Figure 4.**
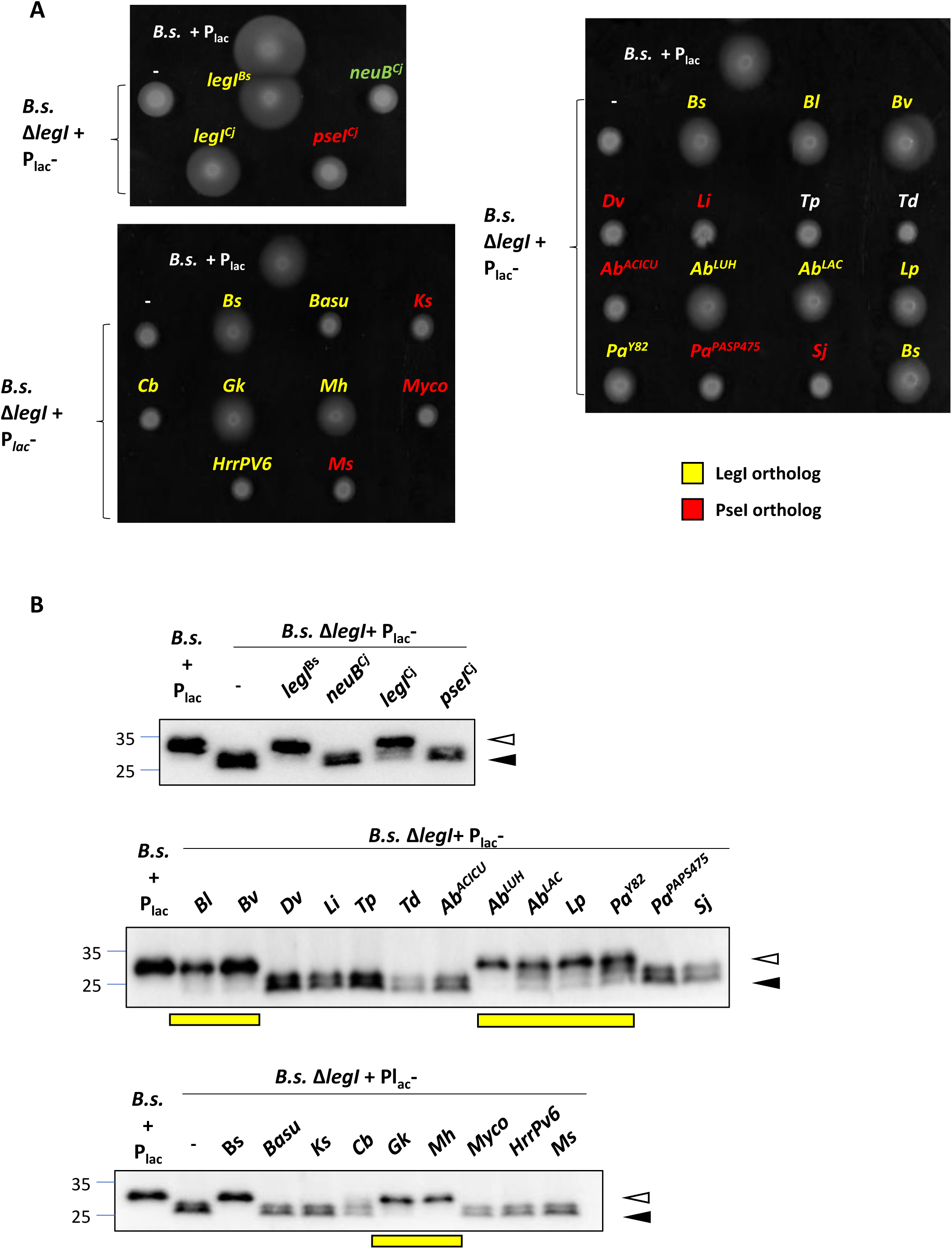
Heterologous complementation of the *B*.*subvibrioides* Δ*legI* mutant with (putative) PseI or LegI orthologs. (A) Motility assays of *B*.*s*. Δ*legI* cells complemented with LegI-type (yellow) or PseI-type (red) synthases expressed from P_*lac*_ on pSRK-Gm. (B) Immunoblots probed with antibodies to FljK^Cc^, revealing the intracellular levels of flagellin in *B*.*s*. Δ*legI* derivatives shown in (A). The blue lines represent the migration of the molecular size standards (in kDa). Bl: *Brevundimonas lutea*, Bv: *Brevundimonas viscosa*, Dv: *Dermabacter vaginalis*, Li: *Leptospira interrogans*, Tp: *Treponema pallidum*, Td: *Treponema denticola*, Ab^ACICU^: *Acinetobacter baumannii* ACICU, Ab^LUH^: *Acinetobacter baumannii* LUH, Ab^LAC^: *Acinetobacter baumannii* LAC-4, Lp: *Legionella pneumophila*, Pa^Y82^: *Pseudomonas aeruginosa* Y82, Pa^PAPS475^: *Pseudomonas aeruginosa* PAPS475, Sj: *Shewanella japonicum*, Basu: *Bacillus subtilis*, Ks: *Kurthia sibirica*, Cb: *Clostridium botulinum*, Gk: *Geobacillus kaustophilus*, Mh: *Moorella humiferrea*, Myco: *Mycobacterium sp. KS0706*, HrrPV6: *Halorubrum sp. PV6*, Ms: *Methanobrevibacter smithii*. All the genes come from synthetic fragments codon optimized for *E. coli* (except the synthase CDS from Gk that is codon optimized for *C*.*c*.). Empty carets indicate the position of modified (glycosylated) flagellin, whereas filled carets mark unmodified flagellin. Yellow boxes indicate LegI functional orthologs.

### LegX, a new molecular marker for Leg biosynthesis pathways

Irrefutable molecular evidence for the complete dissection of glycosylation pathway typically requires the demonstration of sufficiency by reconstitution of glycosylation in a heterologous host expressing a minimal set of the required constituents. Since FlmG^*Cc*^ and FlmG^*Bs*^ are not interchangeable and the glycosyl donor and acceptor specificities must have diverged, we tried to reconstitute the *B. subvibrioides* flagellin glycosylation system in *C. crescentus* using heterologously expressed determinants. To this end, we expressed a synthetic operon of the six *B. subvibrioides* Leg biosynthesis enzymes (i.e. those predicted to be responsible for the production of CMP-Leg from GDP-GlcNAc, Figure 5A) from the *C. crescentus xylX* locus in cells lacking flagellins, PseI^*Cc*^ and FlmG^*Cc*^. This synthetic Leg biosynthesis operon included the following CDSs of the predicted *B. subvibrioides* orthologs of the Leg pathway from *C. jejuni* 11168[22]: Bresu_3266 (LegB), Bresu_0765 (LegC), Bresu_0506 (LegH), Bresu_3264 (LegG), Bresu_0507 (LegI) and Bresu_3265 (LegF)(Figure 1 and 5A). Next, we introduced a plasmid co-expressing FljK^*Bs*^ and FlmG^*Bs*^ and then probed for modification of FljK^*Bs*^ by immunoblotting, asking whether a change in migration of FljK^*Bs*^ was discernible. As shown in Figure 5B, under these conditions the migration of FljK^*Bs*^ was not altered, indicating that i) additional determinants are likely required to execute the glycosylation or that ii) the synthetic CDSs do not express well enough from our plasmids.

**Figure 5.**
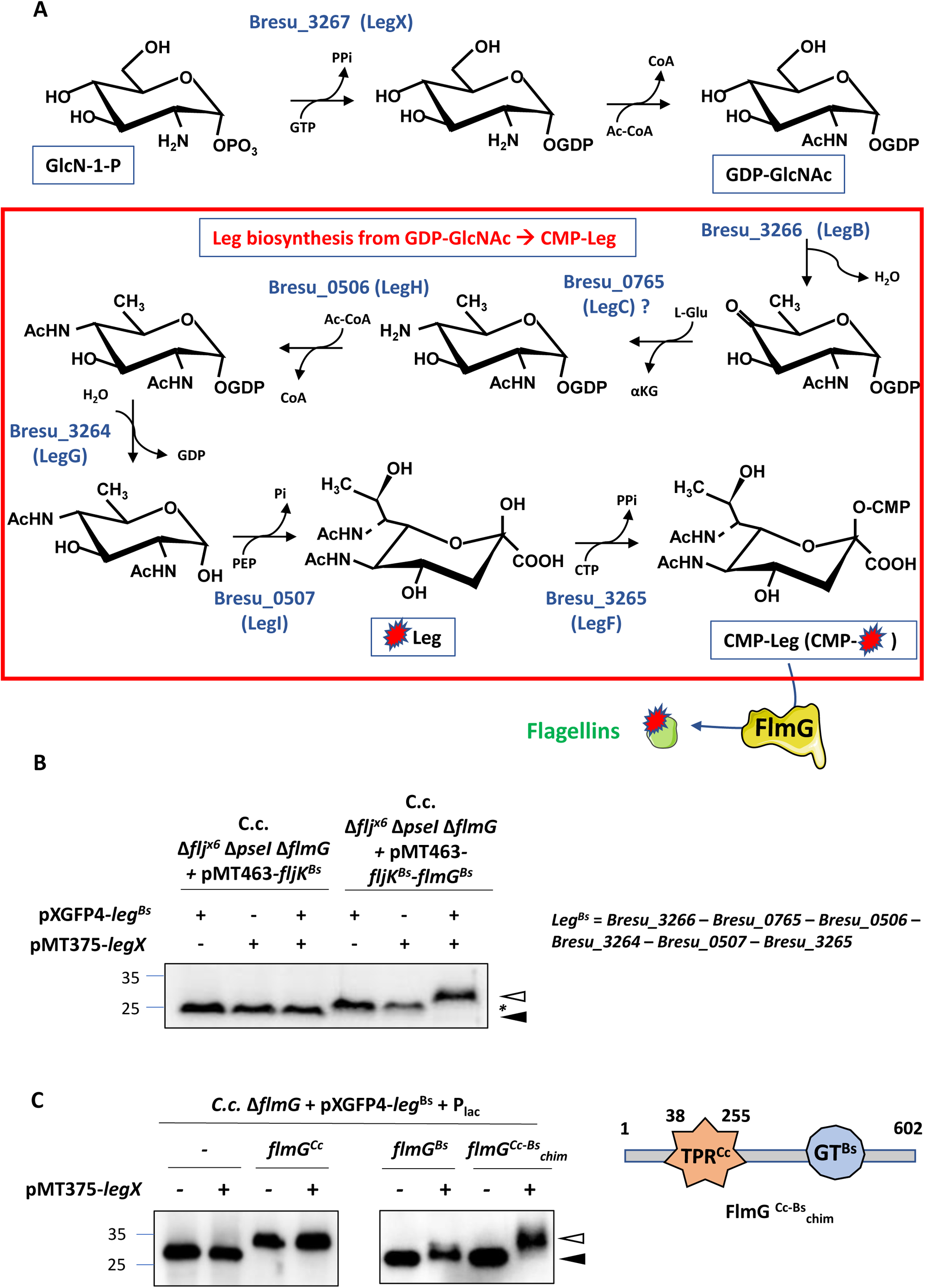
Predicted Leg biosynthetic pathway in *B. subvibrioides*. (A) Schematic of the legionaminic acid biosynthetic pathway as it has been described in *C. jejuni* and elucidated in this study. The pathway requires the addition of GDP into α-D-glucosamine-1-phosphate (GlcN-1P) by Bresu_3267. Then, the different steps are catalyzed by Bresu_3266, Bresu_0765, Bresu_0506, Bresu_3264, Bresu_0507 and Bresu_3265 to ultimately produced CMP-legionaminic acid. The activated legionaminic acid is then transferred to the flagellins by the glycosyltransferase FlmG. (B) Anti-FljK^Cc^ immunoblot analyses of whole cell lysates from *C. crescentus* mutant cultures expressing *fljK*^Bs^_syn_ (codon optimized for *E. coli*) and *flmG*^Bs^ from the replicative pMT463 plasmid, in the presence or absence of a compatible integrative plasmid carrying a *leg*^Bs^ synthetic operon (pXGFP4-*leg*^Bs^) and in the presence or absence of an additional compatible replicative plasmid carrying *Bresu_3267* (pMT375). In the presence of pXGFP4-*leg*^Bs^ (integrated at the *xylX* locus) and pMT375-*Bresu_3267*, FljK^Bs^_syn_ migration is shifted toward higher molecular mass, indicative of glycosylation. Note that in the *C*.*c*. Δ*flj*^x6^ background, the six flagellin-encoding genes have been deleted and the protein detected by the antibodies only corresponds to FljK^Bs^_syn_. The *leg*^Bs^ synthetic operon is composed of *Bresu_3266, Bresu_0765, Bresu_0506, Bresu_3264, Bresu_0507* (*legI*) and *Bresu_3265*. Molecular masses are indicated in kDa by the blue lines. Empty carets indicate the position of modified (glycosylated) flagellin, whereas filled carets mark unmodified flagellin. Asterisk indicates a modification of flagellin by FlmG^Bs^ that does not require Pse or Leg. (C) Anti-FljK^Cc^ immunoblot analyses of whole cell lysates from *C. crescentus* mutant cultures expressing FlmG^Cc^, FlmG^Bs^ and FlmG^Cc-Bs^_chim_ chimera from the replicative pSRK-Gm plasmid (P_*lac*_) in the presence of a compatible integrative plasmid carrying a *leg*^Bs^ synthetic operon (pXGFP4-*leg*^Bs^) and in the presence or absence of an additional compatible replicative plasmid carrying *Bresu_3267* (pMT375). Molecular masses are indicated in kDa by the blue lines. Empty carets indicate the position of modified (glycosylated) flagellin, whereas filled carets mark unmodified flagellin.

Previously we showed that an equivalent synthetic enzyme operon comprising six Pse biosynthesis enzymes was able to direct the synthesis of CMP-Pse from the UDP-GlcNAc precursor in a heterologous host [19]. On the basis of our failure with our corresponding synthetic Leg construct, we considered the possibility that Leg biosynthesis pathway might be incomplete in our heterologous host because the putative precursor, GDP-GlcNAc, is not naturally available in *C. crescentus* (and other bacteria that do not normally synthesize Leg). If true, then this essential biosynthetic activity might also be encoded in Leg biosynthesis gene clusters of *B. subvibrioides* or other Leg-producing bacteria. Upon inspection of the predicted Leg biosynthesis clusters in the genomes of the Gram-negative bacteria *A. baumannii* LAC-4 (GCA_000786735.1)[38] and *P. sp*. Irchel 3E13 (GCA_900187455.1), as well as that of the Gram-positive bacteria *Geobacillus kaustophilus* HTA426 [26, 39] and *Moorella humiferrea* DSM 23265 [40], we noted the presence of one gene encoding an ortholog of PtmE, an enzyme that was used in the enzymatic reconstitution of Leg biosynthesis *in vitro* using enzymes encoded in *C. jejuni* 11168[22]. In these experiments PtmE, a putative guanylyltransferase of GlcN-1-P (α-D-glucosamine-1-phosphate), promoted the production of GDP-GlcNAc *in vitro* (Figure 5A). An ortholog (Bresu_3267, henceforth LegX^*Bs*^) is also encoded adjacent to the genes encoding LegB, LegG and LegF (*Bresu_3266, Bresu_3264* and *Bresu_3265*) orthologs in the *B. subvibrioides* genome (see Figure 1 and 5A).

If LegX^*Bs*^ is indeed required for Leg biosynthesis in *B. subvibrioides*, Δ*legX* cells should recapitulate the motility and flagellin glycosylation defect reported above for Δ*legI* and Δ*flmG* cells. We engineered an in-frame deletion mutation in *legX* and found that the resulting Δ*legX* cells suffer from impaired motility (Figure 6A). Moreover, they neither glycosylate, nor export flagellin (Figure 6B) and TEM revealed only short flagellar filaments on the pole (Figure 6C), as for Δ*legI* and Δ*flmG* cells. If LegX indeed acts in Leg biosynthesis, then it might be possible to restore motility to Δ*legX* cells by expression of a LegX/PtmE ortholog (Figure 6A, 6B), similarly to the heterologous complementation of Δ*legI* cells. This was indeed the case: expression of the *M. humiferrea* LegX ortholog (MOHU_20790) from pSRK-Gm not only restored motility to Δ*legX* cells, but also flagellin glycosylation and export in a manner indistinguishable from the complementation with LegX^*Bs*^ (expressed from pSRK-Gm). As MOHU_20790 exhibits 54% similarity (36% identity) to LegX^*Bs*^ (Table S2), and the predicted fold of LegX (Figure 6D, right) closely resembles that of the nucleotidyltransferase PtmE (Figure 6D, left), we conclude that LegX enzymatic activity is required for motility and Leg biosynthesis in *B. subvibrioides* and that its function in motility can be conferred by LegX orthologs from phylogenetically distant bacteria, such as the Gram-positive bacterium *M. humiferrea*.

**Figure 6.**
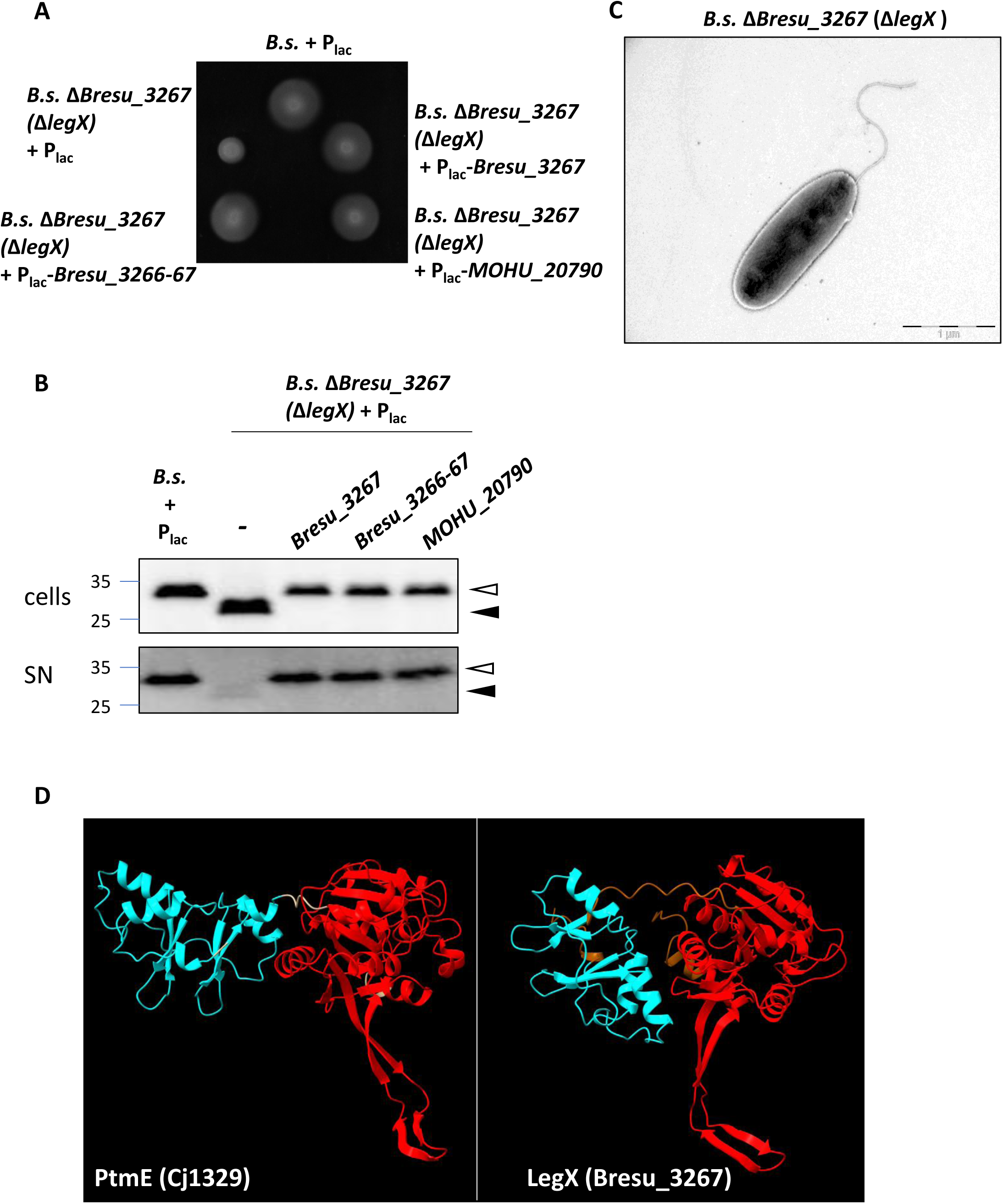
Deletion of *Bresu_3267* (*legX*) encoding a nucleotidyltransferase affects motility, flagellin glycosylation and secretion in *B. subvibrioides*. (A) Motility assay of Δ*Bresu_3267 (*Δ*legX*) *B*.*s*. cells compared to *WT B*.*s*. cells harbouring the empty pSRK-Gm vector (+P_*lac*_) or a complementing derivative with either *Bresu_3267, Bresu_3266-67* or *Moorella humiferrea MOHU_20790*. (B) Immunoblots probed with anti-FljK^*Cc*^ antibodies from cell lysates (cells) and supernatants (SN) of *WT B*.*s*. and Δ*Bresu_3267 (*Δ*legX*) cultures harbouring either the empty pSRK-Gm vector (+P_*lac*_) or a complementing plasmid expressing either Bresu_3267 (LegX), Bresu_3266-67 (LegB-LegX) or the LegX ortholog *M. humiferrea* MOHU_20790. Molecular size standards are indicated by the blue lines with the corresponding value in kDa. Empty carets indicate the position of modified (glycosylated) flagellin, whereas filled carets mark unmodified flagellin. (C) Images of Δ*Bresu_3267 (*Δ*legX*) *B*.*s*. cells analyzed by TEM. (D) Alphafold2 prediction of Bresu_3267 (LegX) from *Brevundimonas subvibrioides* and Cj1329 (PtmE) from *Campylobacter jejuni*. The common modular architecture reveals an N-terminal part containing two tandem repeats of the cystathionine beta-synthase domain superfamily (cyan) and the C-terminal part composed of the nucleotidyltransferase domain (red).

As the *legX* gene lies downstream of the predicted *legB* (*Bresu_3266*) gene, we also inactivated *legB* and observed that the motility of the corresponding mutant (Δ*legB*) is curbed (Figure 7A) and that flagellin glycosylation and export is defective (Figure 7B). However, complementation analyses with plasmids harboring either *legB* or *legB*-*legX* revealed that the Δ*legB* mutation is polar on *legX* expression, indicating that these two genes indeed form an operon (Figure 7A, 7B). We also inactivated the predicted *legH* gene (*Bresu_0506)* that lies upstream of *legI* (*Bresu_0507)*, but in this case there was no evidence of polarity (Figure 7D-7F), despite a similar apparent translational as inferred from the genome sequence. In summary, our analyses show that LegX, LegB and LegH are necessary for Leg- and FlmG-dependent flagellin glycosylation in *B. subvibrioides*. Importantly, LegX is an ideal marker to distinguish Leg from Pse biosynthesis pathways, often embedded in flagellar clusters [26] and, owing to its functional conservation, suitable for the establishment of glyco-profiling set-ups that rely on the motility defect of Δ*legX* cells as proxy.

**Figure 7.**
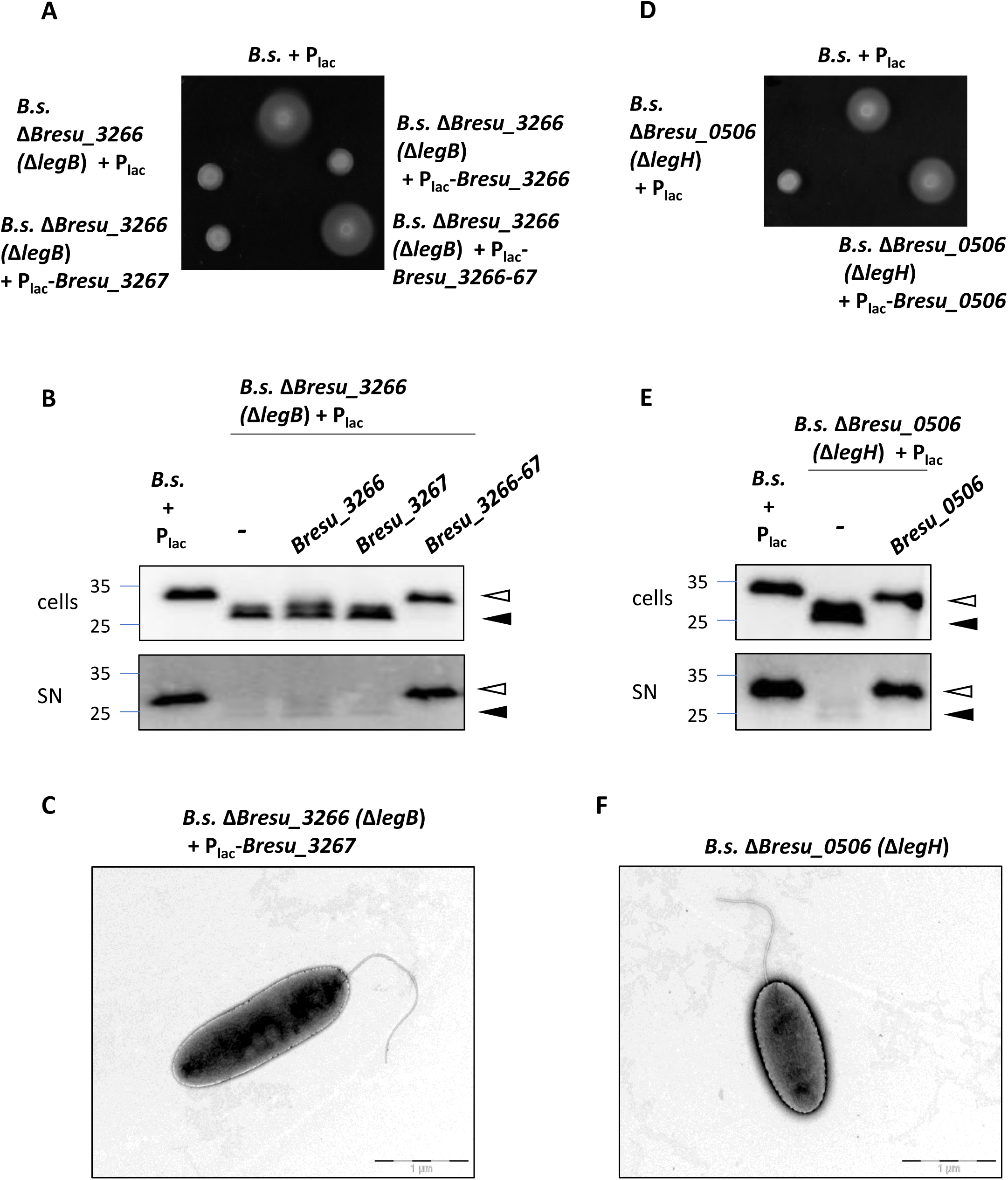
Role of Bresu_3266 (LegB) epimerase/dehydratase and Bresu_0506 (LegH) sialic O-acetyltransferase in flagellar motility of *B. subvibrioides* cells. (A) Motility assays of *WT B*.*s*. and Δ*Bresu_3266* (Δ*legB*) cells expressing Bresu_3266 (LegB), Bresu_3267 (LegX) or Bresu_3266-Bresu_3267 (LegB-LegX, Bresu_3266-67) from P_*lac*_ on plasmid pSRK-Gm. (B) Immunoblots probed with anti-FljK^Cc^ antibodies to reveal flagellins in cell extracts (cells) and supernatants (SN) of the strains described in (A). Empty carets indicate the position of modified (glycosylated) flagellin, whereas filled carets mark unmodified flagellin. (C) TEM images of Δ*Bresu_3266* (Δ*legB*) cells expressing *Bresu_3267* from P_*lac*_ on pSRK-Gm. (D) Motility assay of *WT B*.*s*. and Δ*Bresu_0506* (Δ*legH*) cells complemented with a plasmid expressing Bresu_0506 (LegH) from P_*lac*_ on pSRK-Gm. (E) Immunoblots probed with anti-FljK^Cc^ antibodies to reveal flagellins in cell extracts (cells) and supernatants (SN) of the strains described in (D). (F) TEM images of Δ*Bresu_0506* (Δ*legH*) cells.

### Reconstitution and rewiring of FlmG-dependent flagellin glycosylation

Having unveiled LegX as a critical component of the *B. subvibrioides* Leg-based motility system, we asked whether addition of the LegX enzyme would permit reconstitution of the Leg-dependent flagellin glycosylation by FlmG^*Bs*^ in our recombinant *C. crescentus* cells expressing the other six Leg biosynthesis enzymes. To this end, we transformed a compatible plasmid harboring *B. subvibrioides* LegX CDS (pMT375-*legX*) into the expression *C. crescentus* strains already described above that lack flagellins, PseI and FlmG and performed immunoblot to determine if FlmG^*Bs*^ can support the modification of FljK^*Bs*^ in Leg- and LegX-dependent manner. As shown in Figure 5B, FljK^*Bs*^ was converted to a substantially slower migrating species, a modification that was dependent on the presence of FlmG^*Bs*^ and all seven Leg biosynthesis enzymes (including LegX). Additionally, we observed a barely detectable change in mobility that is FlmG^*Bs*^-dependent, but requires neither Pse, nor Leg (see asterisk, Figure 5B). This change in FljK^*Bs*^ mobility may reflect a certain degree of promiscuity of FlmG towards other donor molecules that are transferred to FljK^*Bs*^.

We hypothesized that the specificity of FlmG enzymes towards Leg versus Pse likely resides in the C-terminal glycosyltransferase (GT-B domain [41]). This hypothesis is based on our previous finding that the N-terminal TPR domain of FlmG^*Cc*^ can bind FljK^*Cc*^, whereas the GT-B alone cannot [19]. Since FlmG^*Bs*^ shares this modular architecture based on sequence analysis, we wondered if a chimeric version of FlmG^*Cc*-*Bs*^ in which we substituted the GT-B domain from *C. crescentus* with that of *B. subvibrioides* would thus glycosylate *C. crescentus* flagellins with Leg. To this end, we used *C. crescentus* Δ*flmG* mutant cells harboring the synthetic six-gene Leg operon at the *xylX* locus. We first transformed these cells with pMT375-*legX*^*Bs*^ and then finally with pSRK-Gm variants expressing either FlmG^*Cc*^, FlmG^*Bs*^, or the chimeric FlmG^*Cc-Bs*^ version. As shown in Figure 5C, the chimeric FlmG^*Cc-Bs*^ was able to modify the *C. crescentus* flagellins in a manner that depended on the presence of LegX^*Bs*^, but it did not modify flagellin in cells producing only Pse (also observed in Figure S3A). Moreover, WT FlmG^*Bs*^ version did not support efficient flagellin modification in the Leg-producing *C. crescentus* cells regardless of whether LegX^*Bs*^ was present or not (Figure 5C). As control, FlmG^*Cc*^ also supported flagellin modification in this system (likely with Pse), because these Δ*flmG* cells produce both Pse and Leg, but only in the presence of pMT375-*legX*^*Bs*^.

In summary, exchanging the C-terminal GT-B domain enabled rewiring the glycosyltransferase specificity from Pse-accepting enzyme to a Leg-accepting enzyme, resulting in the modification of *C. crescentus* flagellins with Leg in cells recombinantly expressing at least seven Leg biosynthesis genes. The fact that such cells are non-motile (Figure S3B) indicates that additional factors exist in the flagellation pathway that exhibit specificity towards the glycosyl group that is joined to flagellins.

## DISCUSSION

### Insulated Leg- or Pse-dependent glycosylation pathways

The exquisite specificity in cellular glycosylation reactions are predetermined to ensure that the desired structures are decorated with the correct sugars. In as much as the underlying glycosyl donor and acceptor selectivity underlie biological function, biotechnological processes often necessitate relaxing these specificities, for example in engineering promiscuous glycosyltransferase (GT) enzymes that can be used to modify a desired target protein with a sugar of choice [42]. Such long-term goals are achievable, but ideally facilitated by the discovery and dissection of the determinants underpinning the GT specificities, including acceptor and donor. In addition to illuminating the molecular mechanism of FlmG fGTs, our work also opens the door towards biotechnological engineering of flagellin-based bio-glycoconjugates using Pse or Leg for example as simple vaccine [43-45] that could serve to combat Pse/Leg in infections by prior immunization not only for *A. baumannii* strains or other pathogens that decorate surfaces with Pse, but, importantly, also for those that contain Leg, including most clinical *A. baumannii* isolates [12, 34, 46].

Our genetic dissection of orthologous FlmG fGTs provided unprecedented insight into the donor sugar and acceptor protein specificities underlying protein glycosylation mechanisms. At the level of the donor, featuring a remarkable stereoisomer selectivity, we showed that the FlmGs from *C. crescentus* and *B. subvibrioides* evolved a strong preference for either Leg or Pse (Figures 2 and 4). Additionally, the stereoisomer specificity of the donor is already reflected in the biosynthesis pathway. Inactivation of the defining synthase enzymes for Leg or Pse, LegI and PseI, yields the same motility defects as the inactivation of the corresponding FlmG enzyme. LegI and PseI cannot substitute for one another in the two flagellation systems that we studied, indicating that the corresponding biosynthesis pathways are genetically (and therefore biochemically) insulated. However, the fact that different PseI orthologs can substitute for the endogenous *C. crescentus* enzyme and, in turn, LegI orthologs can substitute for the endogenous enzyme from *B. subvibrioides* when probing motility, underscores the specificity of the biosynthesis pathways for the two stereoisomers. This stringency lends itself for *in vivo* glyco-profiling using Δ*pseI* and Δ*legI* mutant strains of *C. crescentus* and *B. subvibrioides*, respectively, to functionally probe for Pse or Leg biosynthesis pathways identified in genome searches. Remarkably, such profiling assays not only permit distinction among strains and species, but are also discriminatory across larger phylogenetic distances, including the Gram-positive to Gram-negative divide and even the boundaries between eubacterial and archaeal kingdoms. By extension, having recognized the LegX/PtmE enzyme as a critical element in the Leg-specific enzymatic biosynthesis step (Figure 6) likewise offers another functional, but also a novel bioinformatic, criterion for the correct assignment and discrimination of predicted stereoisomer biosynthesis routes residing in ever-expanding genome databases. The current era of synthetic biology offers unlimited depth to which such synthetic genetic glyco-profiling approaches can be applied.

### Specificity determinants in flagellin glycosyltransferases

Our reconstituted Leg-dependent glycosylation of FljK^*Bs*^ by FlmG^*Bs*^ in *C. crescentus* Δ*pseI* cells using a synthetically assembled Leg-biosynthesis operon, complemented with LegX (Figure 5), allowed us to unambiguously establish the minimal set of components that are required to achieve protein glycosylation using a Leg-based system. The FlmG class of fGTs are suitable subjects for molecular dissection of the underlying specificity determinants because of their conspicuous two-domain architecture that is recognizable by simple primary structure (sequence) comparisons, even without tertiary structural analysis. In fact, the (predicted) bilobed FlmG structure [15] had previously prompted us to determine that the N-terminal TPR domain of FlmG^*Cc*^ confers flagellin (acceptor) recognition, whereas the GT-B domain cannot bind flagellin [19]. Hypothesizing that the GT-B domain could act as determinant for the donor, we considered a simple division of labor model between the two parts of FlmG accounting for the bipartite specificity. Proof for this notion came from the analysis of a chimeric form, FlmG^*Cc-Bs*^, in which the flagellin binding domain from FlmG^*Cc*^ was joined to the GT-B domain of FlmG^*Bs*^. Expression of this chimeric FlmG^*Cc-Bs*^ variant in *C. crescentus* Δ*flmG* cells that had been engineered to synthesize Leg resulted in the modification of the *C. crescentus* flagellin, whereas the *WT* version of FlmG^*Bs*^ had poor activity (Figure 5C). Conversely, FlmG^*Cc-Bs*^ was unable to support glycosylation of *C. crescentus* flagellins with Pse, likely because it no longer possesses the Pse-specific GT-B domain of FlmG^*Cc*^.

Similar dissection experiments should be conducted with the other class of fGTs that are wide-spread in bacteria, the Mafs [10, 26, 28, 47, 48], to reveal if analogous mechanisms and determinants underpin flagellin glycosylation in these systems. It stands to reason that donor and acceptor specificities exist in Mafs as well, however, the flagellin recognition determinants remain unknown. An X-ray structure determined for the Maf from *M. magneticum* [28] revealed a tripartite domain architecture with central GT-A domain bearing clear resemblance to the GT29 and GT42 family of sialyltransferases. The GT-A domain is a characteristic of the Mafs (also known as the signature MAF_flag10 domain) and is likely to confer Pse donor specificity. In fact, our glyco-profiling revealed the corresponding synthase of *M. magneticum* to have PseI activity in our motility assay (Figure S1) and our sequence analysis by BlastP easily discerned a complete (predicted) Pse-biosynthesis pathway encoded in its genome. While the flagellin binding determinant was not evident in the *M. magneticum* Maf structure, a weak structural similarity with flagellin and flagellin secretion chaperones may point to a C-terminal flagellin recognition determinant. However, it remains to be determined whether this region is necessary and sufficient for flagellin binding.

Khairnar *et al*. [26] provided evidence of some donor promiscuity in the Maf from *G. kaustophilus* that is encoded in flagellar locus. When this Maf was expressed in recombinant *Escherichia coli* cells that synthesize sialic acid, modification of the co-expressed *G. kaustophilus* flagellin was seen with sialic acid. However, it would be interesting to test if the efficiency of flagellin glycosylation by *G. kaustophilus* Maf is increased in a heterologous host producing Leg as *maf* gene is adjacent to Leg biosynthesis genes, including a LegX ortholog (Table S2) and our glyco-profiling in Figure 4 revealed that *G. kaustophilus* indeed encodes a LegI ortholog. Overall, it remains to be determined whether the Mafs are inherently more promiscuous than the FlmG enzymes.

### Leg- and Pse-based glycosylation in the (same) prokaryotic cell

The donor specificity observed with the two FlmG enzymes studies in our work and the possible specificity inferred for Maf-encoding gene clusters may ensure that the correct cellular structure is modified with the right donor. In our experiments when *C. crescentus* cell synthesizing Leg were used, FlmG^*Cc*^ had a clear preference to modify flagellin with Pse, rather than Leg. Thus, the terminal determinant of a given glycosylation pathway governs selectivity of CMP-Pse over CMP-Leg or vice versa. Our work also indicates that the biosynthesis pathways themselves are kept insulated by dedicated enzymes, perhaps to prevent the formation of Leg/Pse hybrid intermediates that would otherwise create too much chemical variability for the systems to function properly in their biological roles. Pse and Leg glycosylation systems are used for other cell surface structures, not only flagellins [1, 16, 49]. While Leg or Pse biosynthesis enzymes are often encoded in flagellar gene clusters, they can also occur within O-antigen or capsular gene clusters, sometimes even in the same genome. In this situation, a possible enzymatic interference of the Leg and Pse biosynthesis pathways must be avoided. In bacterial cells, it might be possible to restrict Leg or Pse synthesis to specific (mutually exclusive) growth conditions, but a perhaps more common solution is to design each biosynthetic pathway with specific chemical marks. An appealing hypothesis is that this is achieved with the synthesis of Leg from GDP-activated precursors, whereas Pse synthesis occurs from UDP-activated molecules [21, 22]. This dependency on GDP for Leg also comes at a price, since at least one specific conversion enzyme, such as LegX, is required to launch the biosynthesis pathway with the production of the activated precursor bearing the GDP mark (Figure 6).

For such chemical complexity in the biosynthesis of stereoisomers to evolve, these glycosylation systems must be of considerable value to cells, begging the question of their function. Do surface modifications with Pse or Leg just serve to generate epitopes or envelopes with different modifications or are there special physical or chemical properties associated with Pse or Leg. As members of the sialic family of molecules they certainly have the potential to function as innate immune modulators, but Leg or Pse also found on environmental bacteria that are not known to associate with eukaryotic cells. While Pse and Leg play an important role in *C. crescentus* and *B. subvibrioides* flagellation, respectively, many other flagellation systems exist that do not require Pse or Leg. On the basis of this fact, it is conceivable that glycosylation does not fulfill a conserved role in flagellar assembly in general, but we cannot exclude that it has been appropriated for regulatory purposes in some bacterial flagellation systems. We observed that modifying FljK^*Cc*^ with Leg in our recombinant *C. crescentus* system still did not restore motility, suggesting that the type of sugar modification does matter, possibly because other flagellin interacting proteins such as the FlaF secretion chaperone, capping proteins or unknown factors no longer interact or function properly with FljK^*Cc*^ that does not harbor the Pse modification. Alternatively, or additionally, it is conceivable that modification of the flagellar filament with Pse or Leg simply protects against infection by certain flagellotropic phages, in a manner analogous to that reported recently for pilus glycosylation in *Pseudomonas aeruginosa [50]*.

## EXPERIMENTAL PROCEDURES

### Strains and growth conditions

Bacterial strains used in this study are listed in Table S3. *C. crescentus* and *B. subvibrioides* strains were grown at 30°C in peptone-yeast extract (PYE) (2g/L bacto-peptone, 1g/L yeast extract, 1 mM MgSO_4_ and 0.5 mM CaCl_2_)[51]. *E. coli* S17-1 λ*pir* and EC100D were grown at 37°C in LB. Antibiotics were added to the medium at the following concentration (µg/mL in liquid/solid medium for *C. crescentus* and *B. subvibrioides*; µg/mL in liquid/solid medium for *E. coli*): nalidixic acid (20 only in solid medium for *C. crescentus* and *B. subvibrioides*), tetracycline (1/1; 10/10), kanamycin (5/20; 20/20), gentamycin (1/1; 25/25). Gene expression was induced when required with 50 µM vanillate, 0.3% D-xylose or 0.5 mM isopropyl-beta-D-thiogalactoside (IPTG) for *C. crescentus* and *B. subvibrioides* cultures. Electroporation, bi-parental mating and motility assays were performed as previously described in *C. crescentus* and *B. subvibrioides* [51, 52].

For motility assays, 1 µL of overnight cultures were spotted on soft (0.3%) agar plates with the corresponding antibiotics and inducers (IPTG or vanillate) and incubated for 3 days and 7 days for *C. crescentus* and *B. subvibrioides*, respectively.

### Immunoblots

For immunoblots, protein samples were prepared from cells harvested in the middle of the exponential growth phase (1 mL at OD_600nm_≈0.4). Proteins samples were separated on SDS polyacrylamide gel, transferred to polyvinylidene difluoride (PVDF) Immobilon-P membranes (Merck Millipore) and blocked in Tris-buffered saline (TBS) 0.1% Tween-20% and 5% dry milk [19]. The anti-FljK^Cc^ anti-serum (raised against His6-FljK expressed in *E. coli* [19]) was used at 1:10,000 dilution. Protein-primary antibody complexes were revealed with horseradish peroxidase-labeled donkey anti-rabbit antibodies (Jackson ImmunoResearch, West Grove, PA) and ECL detection reagents (Amersham, GE Healthcare, Glattbrugg, Switzerland).

### Negative stain transmission electron microscopy

Samples for negative stain TEM were prepared by first glow discharging 200-mesh copper, carbon-coated, formvar grids (EM Science, Hatfield, PA) for 1 min. 20 µL of exponential cultures of *B. subvibrioides* were applied to the grids and allowed to adsorb for 1 min before being washed three times in water, stained with 1% uranyl acetate for 1 min and washed with water for 30 sec. Negatively stained *B. subvibrioides* were imaged on a Tecnai 20 (FEI Company, Eindhoven, Netherland). Flagellum length measurement was performed using the ImageJ software.

### Derivation of nonulosonic acids (NulOs) with DMB and analysis on HPLC

We extracted NulOs from lyophilized purified flagella from culture supernatants. Briefly, 250 mL of an overnight culture (24 h for *B. subvibrioides*) was spun for 15 min at 8,000 r.p.m. at 4°C to remove cells. Shed flagella were then pelleted from the culture supernatant by ultracentrifugation at (27,000 r.p.m. 30 min, 15°C), washed with 50 mL water and pelleted again by ultracentrifugation. Purified flagella were resuspended in water and frozen at -80°C prior to lyophilization. 1,2-diamino-4,5-methylene dioxybenzene (DMB) was used to derivatize NulOs as previously described [28]. Briefly, dried glycoconjugates were hydrolyzed in 0.1 M trifluoroacetic acid for 2 h at 80°C to release NulOs. NulOs were coupled to DMB for 2 h at 50°C in the dark in a derivation solution (7 mM DMB; 1 M β-mercaptoethanol; 18 mM sodium hydrosulfite; 0.02 mM trifluoroacetic acid). NulO derivatives were separated isocratically on a C_18_ reverse-phase HPLC column (Thermo Scientific, Hypersil ODS, 4.6 mm by 250 mm, 5 µm) using acetonitrile/methanol/water (7:9:84 vol/vol/vol) mixture solvent and detected by a fluorimeter (Waters 2475, excitation wavelength λ_exc_=373 nm, emission wavelength λ_em_=448 nm.

### Strain and plasmid constructions

For in-frame deletions, bi-parental mating was used to deliver the corresponding pNPTS138 derivatives (listed in Table S3) into *B. subvibrioides* strains. Double recombination was selected by plating bacteria onto PYE plates supplemented with 3% sucrose. Putative mutants were confirmed by PCR using primers (listed in Table S4) external to the DNA fragments used for the pNPTS138 constructs.

pNK562: PCR was used to amplify two fragments flanking the *flmG* (*Bresu_2406*) ORF with Bs_flmG_del_1/Bs_flmG_del_2 and Bs_flmG_del_3/Bs_flmG_del_4. The PCR fragments were digested with *Mfe*I/*Bam*HI and *Bam*HI/*Hin*dIII, respectively and triple ligated into pNPTS138 restricted with *Eco*RI/*Hin*dIII.

pNK580: PCR was used to amplify two fragments flanking the *neuB* (*Bresu_0507*) ORF with Bs_neuB_del_1/Bs_neuB_del_2 and Bs_neuB_del_3/Bs_neuB_del_4. The PCR fragments were digested with *Eco*RI/*Bam*HI and *Bam*HI/*Hin*dIII, respectively and triple ligated into pNPTS138 restricted with *Eco*RI/*Hin*dIII.

pNK926: PCR was used to amplify two fragments flanking the *Bresu_3266* ORF with NK339/NK340 and NK341/342. The PCR fragments were digested with *Hin*dIII/*Bam*HI and *Bam*HI/*Eco*RI, respectively and triple ligated into pNPTS138 restricted with *Eco*RI/*Hin*dIII.

pNK1000: PCR was used to amplify two fragments flanking the *Bresu_3267* ORF with NK345/NK346 and NK347/348. The PCR fragments were digested with *Hin*dIII/*Kpn*I and *Kpn*I/*Eco*RI, respectively and ligated into pNPTS138 restricted with *Eco*RI/*Hin*dIII.

pNK1002: PCR was used to amplify two fragments flanking the *Bresu_0506* ORF with NK366/NK367 and NK368/369. The PCR fragments were digested with *Hin*dIII/*Bam*HI and *Bam*HI/*Eco*RI, respectively and ligated into pNPTS138 restricted with *Eco*RI/*Hin*dIII.

Inducible plasmids were constructed with a *Nde*I site overlapping the start codon and an *Xba*I site (or *Eco*RI site when mentioned) flanking the stop codon were constructed as follows:

pNK660: the *flmG* ORF was amplified by PCR with Bs-flmG-NdeI/ Bs-flmG-XbaI. The PCR fragment was digested by *Nde*I/*Xba*I and ligated into pSRK-Gm [37] restricted with *Nde*I/*Xba*I.

pNK631: the synthetic fragment encoding the *legI* (*Bresu_0507*) CDS, codon optimized for *E. coli* (see Table X), was subcloned into pSRK-Gm from pUCIDT plasmid using *Nde*I/*Xba*I.

pNK948: the *Bresu_3266* CDS was amplified by PCR with NK357/358. The PCR fragment was digested by *Nde*I/*Xba*I and ligated into pSRK-Gm restricted with *Nde*I/*Xba*I.

pNK950: the *Bresu_3267* CDS was amplified by PCR with NK359/360. The PCR fragment was digested by *Nde*I/*Xba*I and ligated into pSRK-Gm restricted with *Nde*I/*Xba*I.

pNK988: the *Bresu_3266-67* CDSs were amplified by PCR with NK357/360. The PCR fragment was digested by *Nde*I/*Xba*I and ligated into pSRK-Gm restricted with *Nde*I/*Xba*I.

pNK974: the *Bresu_0506* CDS was amplified by PCR with NK374/375. The PCR fragment was digested by *Nde*I/*Xba*I and ligated into pSRK-Gm restricted with *Nde*I/*Xba*I.

pNK957: the *Bresu_3267* CDS was amplified by PCR with NK359/360, digested by *Nde*I/*Xba*I and cloned into pMT375.

pSA228: the synthetic fragment encoding *fljK*^*Bs*^ (*Bresu_2638*, codon optimised for *C. crescentus*) CDS was digested by *Nde*I/*Eco*RI and ligated into pMT463 [53] restricted by *Nde*I/*Eco*RI.

pLT2043: the *neuB* CDS of *Pseudomonas irchel 3A5* was amplified with 3A5_PseI_NdeI/3A5_PseI_mfeI. The PCR fragment was digested by *Nde*I/*Mfe*I and ligated into pMT335 restricted with *Nde*I/*Eco*RI.

pLT2036: the *legI* CDS of *B. subvibrioides* was amplified with Bs_neuB_nde/Bs_neuB_eco. The PCR fragment was digested by *Nde*I/*Eco*RI and ligated into pMT335 [53] restricted with *Nde*I/*Eco*RI.

pLT2237: the *pseI* CDS of Kurthia was amplified with Ku_neuB_nde/Ku_neuB_eco. The PCR fragment was digested by *Nde*I/*Eco*RI and ligated into pMT335 restricted with *Nde*I/*Eco*RI.

pLT2262: the *pseI* CDS of *M. magneticum* was amplified with Mm_neuB_nde/Mm_neuB_eco. The PCR fragment was digested by *Nde*I/*Eco*RI and ligated into pMT335 restricted with *Nde*I/*Eco*RI.

pLT2263: the *pseI* CDS of *S. oneidensis* was amplified with So_neuB_nde/So_neuB_eco. The PCR fragment was digested by *Nde*I/*Eco*RI and ligated into pMT335 restricted with *Nde*I/*Eco*RI.

To create the vector co-expressing *fljK*^*Bs*^ and *flmG*^*Bs*^, the *flmG* ORF was amplified by PCR with primers Bs_flmG_rbs_Eco/Bs_flmG_Xba (with ribosome binding site and EcoRI site flanking the *flmG* start codon and XbaI site flanking the *flmG* stop codon) and digested by *Eco*RI/*Xba*I. The digested fragment was subcloned into pSA228[19] restricted by *Eco*RI/*Xba*I.

To express the *legX* ortholog from *Moorella humiferrea*, the *MOHU_20790* ORF (codon optimized for *E. coli*) was amplified by PCR using NK361 and M13(−48) primers from pUC-GW-MOHU_20790syn plasmid. After digestion by *Nde*I/*Xba*I, the fragment was ligated into pSRK-Gm restricted by *Nde*I/*Xba*I to generate pNK955.

pLT2295: to express the six enzyme *B. subvibrioides* legionaminic acid biosynthesis pathway in *C. crescentus*, a synthetic operon (codon-optimized for *E. coli*, see Table S4) encoding Bresu_3266, Bresu_0765, Bresu-0506, Bresu-3264, Bresu-0507 and Bresu-3265 was subcloned from pUC-GW plasmid to pXGFP4 using *Nde*I/*Xba*I.

To express *pseI*^Cj^, *legI*^Cj^ and *neuB*^Cj^, the corresponding CDSs were individually subcloned from pSA126, pSA47 and pSA48 [19] to pSRK-Gm using *Nde*I/*Xba*I to generate pNK991, pNK992 and pNK994, respectively.

To express heterologous PseI or LegI orthologs, the synthetic fragments harboring the CDSs (codon optimized for *E. coli*, see Table S4) were subcloned from pUC-GW or pUCIDT plasmids to pSRK-Gm using *Nde*I/*Xba*I (Table S3).

## ACKNOWLEDGEMENTS

We thank Laurence Degeorges for excellent technical assistance and Silvia Ardissone for the plasmids expressing *B. subvibrioides* flagellins and FlmG. We also thank Bohumil Maco for the help with TEM experiments. Funding support was from the Swiss National Science Foundation (31003A_182576), the University of Geneva (DIP) and UNITEC (InnogapR23-24) to P.H.V, and a Swisslife Foundation (Jubiläumsstiftung) grant for medical research to N.K..

## FIGURE LEGENDS

**Figure S1.**
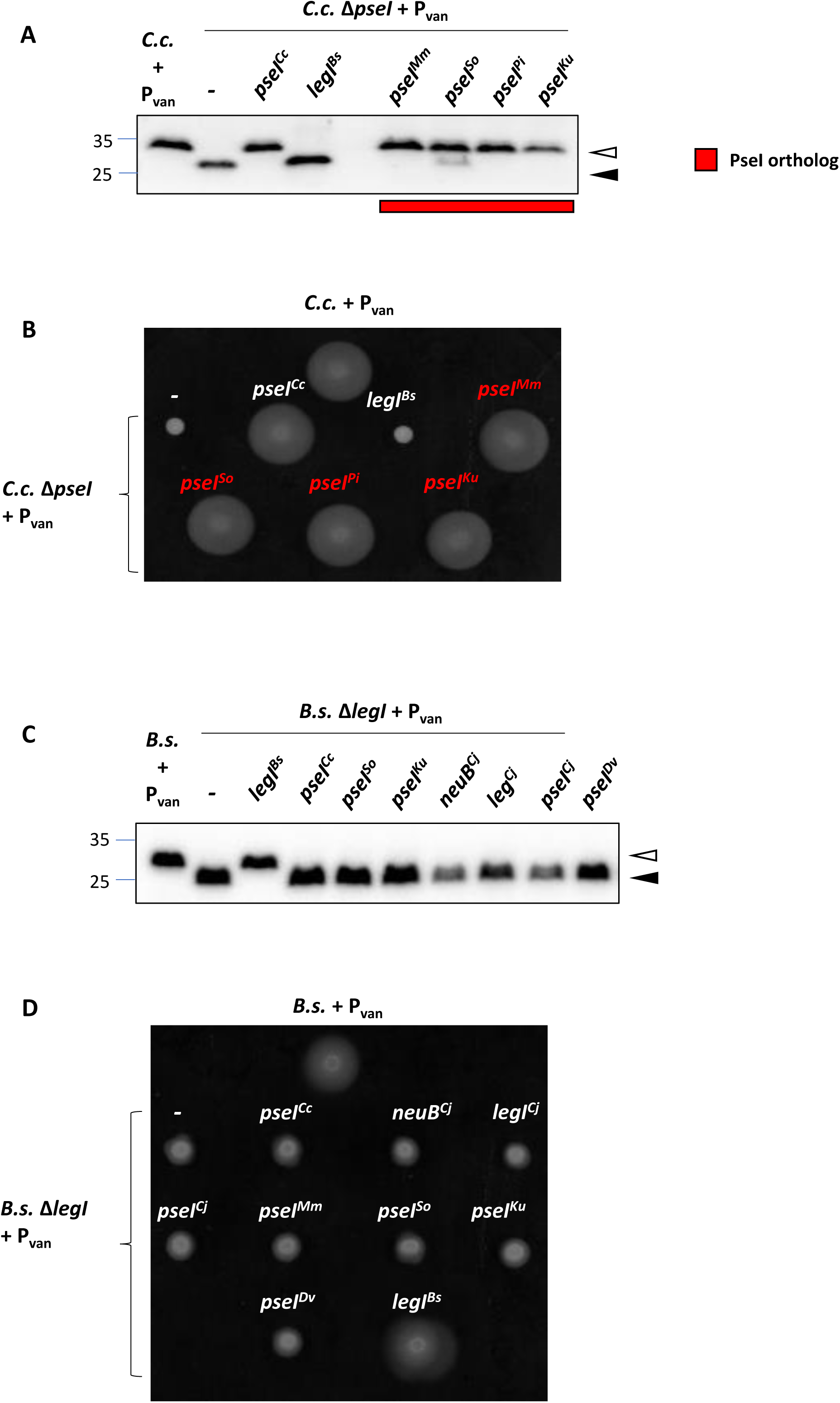
Complementation of *C. crescentus* Δ*pseI* cells and *B. subvibrioides* Δ*legI* cells with plasmids expressing PseI-like synthase (sequence) orthologs. (A) Immunoblots probed with anti-FljK^Cc^ antibodies from extracts of *C*.*c*. Δ*pseI* cells expressing the orthologs from the vanillate-inducible P_*van*_ promoter on plasmid pMT335 (+P_*van*_). Molecular masses are indicated by the blue lines in kDa. Empty carets indicate the position of modified (glycosylated) flagellin, whereas filled carets mark unmodified flagellin. (B) Motility assay of the strains described in (A). (C) Immunoblot performed on *B*.*s*. Δ*legI* cells harboring CDSs (*pseI*^*Cc*^, *pseI*^*So*^, *pseI*^*Ku*^, *pseI*^*Cj*^) for PseI-like proteins, LegI-like proteins (*legI*^*Bs*^, *legI*^*Cj*^) or a NeuB-like synthase (*neuB*^*Cj*^) under P_van_ control. Blue lines indicate the molecular masses in kDa. Empty carets indicate the position of modified (glycosylated) flagellin, whereas filled carets mark unmodified flagellin. (D) Motility assay of the strains described in (C).

**Figure S2.**
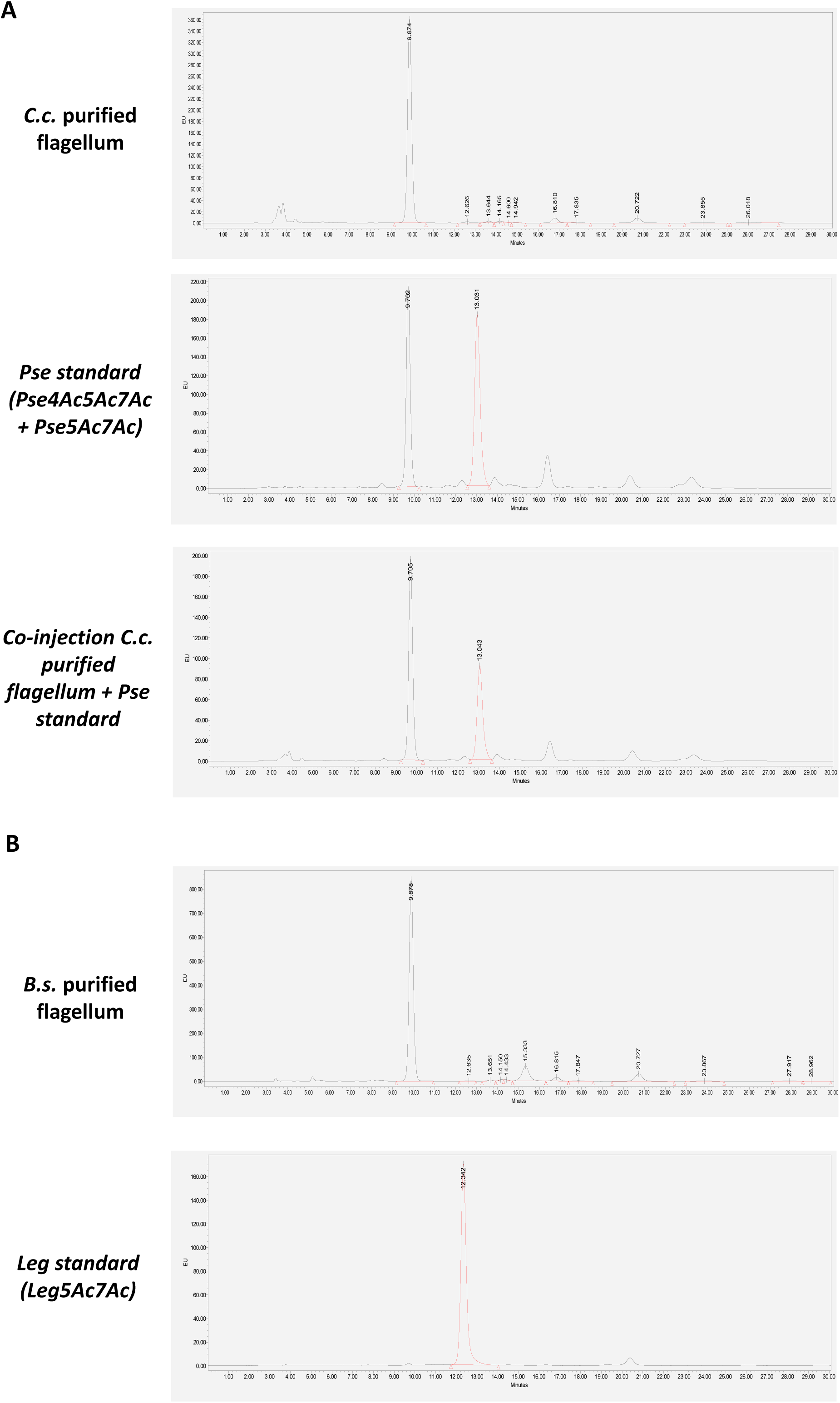
Pseudaminic acid is detected in *C. crescentus* purified flagella. (A) Chromatograms of RP-HPLC-FL experiments performed on purified flagella from *C. crescentus* WT cells. The potential nonulosonic acids (NulOs) produced by *C. crescentus* and present on the flagellum have been derivatized by DMB, a fluorogenic reagent that shows high specificity for NulOs and analyzed by reverse-phase high performance liquid chromatography coupled to fluorescence (RP-HPLC-FL). One major peak (retention times of 9.8 min) is detected in purified flagellum. This retention time perfectly overlaps with the first peak observed for pseudaminic acid standard composed of Pse4Ac5Ac7Ac and Pse5Ac7Ac isolated from the capsule of *A. baumannii* NIPH329 when co-injected. (B) Chromatograms of RP-HPLC-FL experiments performed on purified flagella from *B. subvibrioides* WT cells. One major peak (retention times of 9.8 min) and one minor peak (15.4 min) are detected in purified flagellum. These retention times are different to the Leg standard (Leg5Ac7Ac) used and isolated from the *A. baumannii* LUH5533 capsule. Chromatograms are representative of three independent experiments.

**Figure S3.**
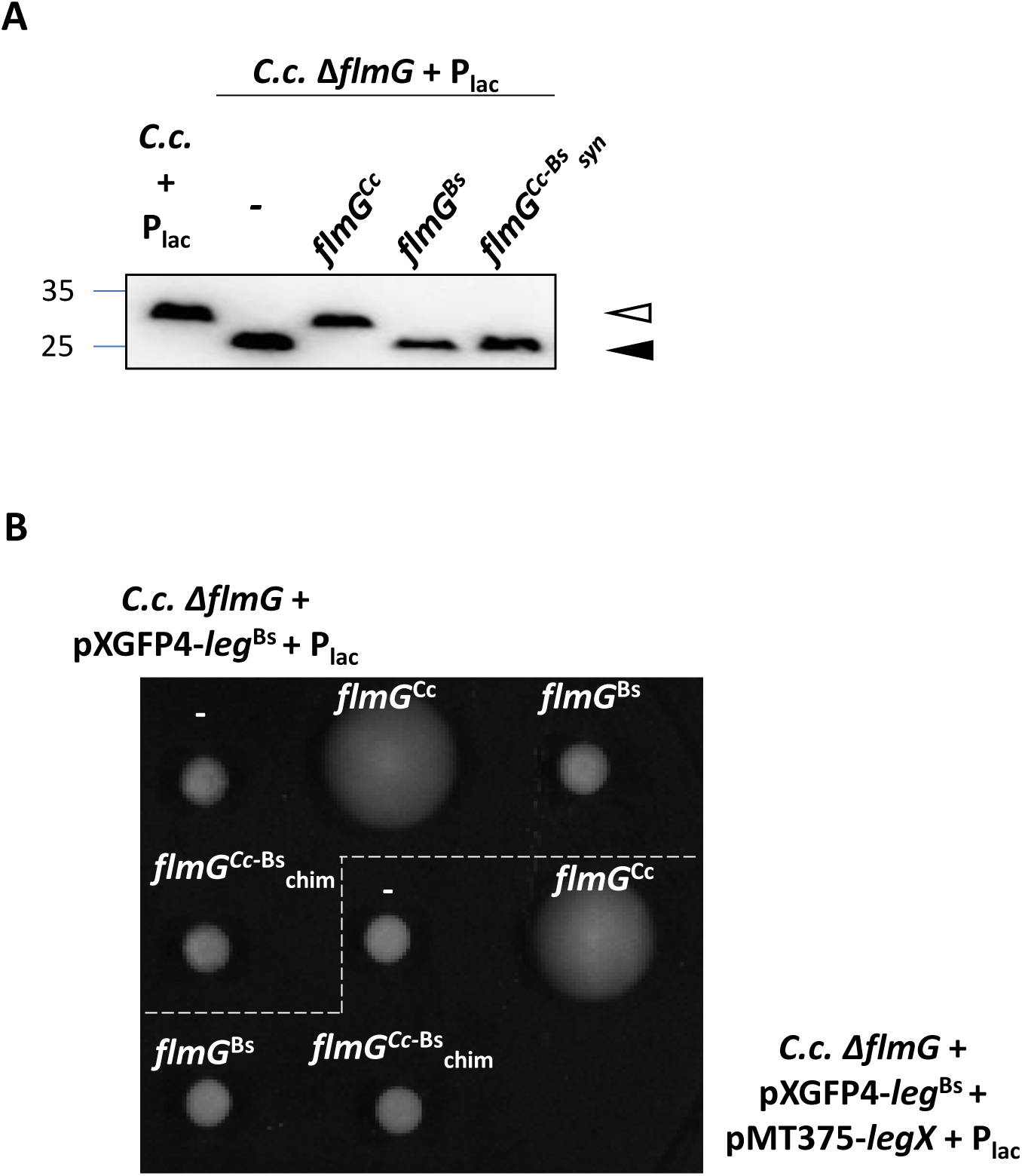
FlmG orthologs are not interchangeable between *C. crescentus*. and *B. subvibrioides*. (A) Immunoblots probed with antibodies to FljK^*Cc*^ antibodies to reveal flagellin in lysates of *C*.*c*. Δ*flmG* cells expressing *flmG* from *C*.*c*. and *B*.*s*. or a gene that codes for a chimeric FlmG composed of the N-terminal part of FlmG^*Cc*^ (harboring the TPR domain that promotes FlmG-Flagellins interaction) and the C-terminal part of FlmG^*Bs*^ (carrying the glycosyltransferase domain) from P_*lac*_ on plasmid pSRK-Gm. Empty carets indicate the position of modified (glycosylated) flagellin, whereas filled carets mark unmodified flagellin. (B) Motility assays of the strains described in Figure 5C: *C. crescentus* mutant expressing FlmG^Cc^, FlmG^Bs^ and FlmG^Cc-Bs^_chim_ chimera from the replicative pSRK-Gm plasmid (P_*lac*_) in the presence of a compatible integrative plasmid carrying a *leg*^Bs^ synthetic operon (pXGFP4-*leg*^Bs^) and in the presence or absence of an additional compatible replicative plasmid carrying *Bresu_3267* (pMT375).

## Notes

### Competing Interest Statement

The authors have declared no competing interest.

## REFERENCES

1. McDonald, N.D., and Boyd, E.F. (2021). Structural and Biosynthetic Diversity of Nonulosonic Acids (NulOs) That Decorate Surface Structures in Bacteria. Trends Microbiol 29, 142–157.

2. Varki, A., Schnaar, R.L., and Schauer, R. (2015). Sialic Acids and Other Nonulosonic Acids. In Essentials of Glycobiology, rd, A. Varki, R.D. Cummings, J.D. Esko, P. Stanley, G.W. Hart, M. Aebi, A.G. Darvill, T. Kinoshita, N.H. Packer, et al., eds. (Cold Spring Harbor (NY)), pp. 179–195.

3. Zunk, M., and Kiefel, M.J. (2014). The occurrence and biological significance of the [small alpha]-keto-sugars pseudaminic acid and legionaminic acid within pathogenic bacteria. RSC Advances 4, 3413–3421.

4. Le Quere, A.J., Deakin, W.J., Schmeisser, C., Carlson, R.W., Streit, W.R., Broughton, W.J., and Forsberg, L.S. (2006). Structural characterization of a K-antigen capsular polysaccharide essential for normal symbiotic infection in Rhizobium sp. NGR234: deletion of the rkpMNO locus prevents synthesis of 5,7-diacetamido-3,5,7,9-tetradeoxy-non-2-ulosonic acid. J Biol Chem 281, 28981–28992.

5. Knirel, Y.A., Shashkov, A.S., Tsvetkov, Y.E., Jansson, P.E., and Zahringer, U. (2003). 5,7-diamino-3,5,7,9-tetradeoxynon-2-ulosonic acids in bacterial glycopolymers: chemistry and biochemistry. Adv Carbohydr Chem Biochem 58, 371–417.

6. Tomek, M.B., Janesch, B., Maresch, D., Windwarder, M., Altmann, F., Messner, P., and Schaffer, C. (2017). A pseudaminic acid or a legionaminic acid derivative transferase is strain-specifically implicated in the general protein O-glycosylation system of the periodontal pathogen Tannerella forsythia. Glycobiology 27, 555–567.

7. Horzempa, J., Dean, C.R., Goldberg, J.B., and Castric, P. (2006). Pseudomonas aeruginosa 1244 pilin glycosylation: glycan substrate recognition. J Bacteriol 188, 4244–4252.

8. Thibault, P., Logan, S.M., Kelly, J.F., Brisson, J.R., Ewing, C.P., Trust, T.J., and Guerry, P. (2001). Identification of the carbohydrate moieties and glycosylation motifs in Campylobacter jejuni flagellin. J Biol Chem 276, 34862–34870.

9. Schirm, M., Soo, E.C., Aubry, A.J., Austin, J., Thibault, P., and Logan, S.M. (2003). Structural, genetic and functional characterization of the flagellin glycosylation process in Helicobacter pylori. Mol Microbiol 48, 1579–1592.

10. Meng, X., Boons, G.J., Wosten, M., and Wennekes, T. (2021). Metabolic Labeling of Legionaminic acid in Flagellin Glycosylation of Campylobacter jejuni Identifies Maf4 as a Putative Legionaminyl Transferase. Angew Chem Int Ed Engl.

11. Andolina, G., Wei, R., Liu, H., Zhang, Q., Yang, X., Cao, H., Chen, S., Yan, A., Li, X.D., and Li, X. (2018). Metabolic Labeling of Pseudaminic Acid-Containing Glycans on Bacterial Surfaces. ACS Chemical Biology 13, 3030–3037.

12. Wei, R., Yang, X., Liu, H., Wei, T., Chen, S., and Li, X. (2021). Synthetic Pseudaminic-Acid-Based Antibacterial Vaccine Confers Effective Protection against Acinetobacter baumannii Infection. ACS Cent Sci 7, 1535–1542.

13. Nedeljkovic, M., Sastre, D.E., and Sundberg, E.J. (2021). Bacterial Flagellar Filament: A Supramolecular Multifunctional Nanostructure. International Journal of Molecular Sciences 22, 7521.

14. Chevance, F.F., and Hughes, K.T. (2008). Coordinating assembly of a bacterial macromolecular machine. Nature reviews. Microbiology 6, 455–465.

15. Kint, N., Unay, J., and Viollier, P.H. (2022). Specificity and modularity of flagellin nonulosonic acid glycosyltransferases. Trends Microbiol 30, 109–111.

16. Szymanski, C.M., and Wren, B.W. (2005). Protein glycosylation in bacterial mucosal pathogens. Nature reviews. Microbiology 3, 225–237.

17. Valguarnera, E., Kinsella, R.L., and Feldman, M.F. (2016). Sugar and Spice Make Bacteria Not Nice: Protein Glycosylation and Its Influence in Pathogenesis. Journal of Molecular Biology 428, 3206–3220.

18. Harding, C.M., and Feldman, M.F. (2019). Glycoengineering bioconjugate vaccines, therapeutics, and diagnostics in E. coli. Glycobiology 29, 519–529.

19. Ardissone, S., Kint, N., and Viollier, P.H. (2020). Specificity in glycosylation of multiple flagellins by the modular and cell cycle regulated glycosyltransferase FlmG. Elife 9.

20. Friedrich, V., Janesch, B., Windwarder, M., Maresch, D., Braun, M.L., Megson, Z.A., Vinogradov, E., Goneau, M.F., Sharma, A., Altmann, F., et al. (2017). Tannerella forsythia strains display different cell-surface nonulosonic acids: biosynthetic pathway characterization and first insight into biological implications. Glycobiology 27, 342–357.

21. Schoenhofen, I.C., McNally, D.J., Brisson, J.R., and Logan, S.M. (2006). Elucidation of the CMP-pseudaminic acid pathway in Helicobacter pylori: synthesis from UDP-N-acetylglucosamine by a single enzymatic reaction. Glycobiology 16, 8C–14C.

22. Schoenhofen, I.C., Vinogradov, E., Whitfield, D.M., Brisson, J.R., and Logan, S.M. (2009). The CMP-legionaminic acid pathway in Campylobacter: biosynthesis involving novel GDP-linked precursors. Glycobiology 19, 715–725.

23. Lewis, A.L., Desa, N., Hansen, E.E., Knirel, Y.A., Gordon, J.I., Gagneux, P., Nizet, V., and Varki, A. (2009). Innovations in host and microbial sialic acid biosynthesis revealed by phylogenomic prediction of nonulosonic acid structure. Proc Natl Acad Sci U S A 106, 13552–13557.

24. Vieira, A.Z., Raittz, R.T., and Faoro, H. (2021). Origin and evolution of nonulosonic acid synthases and their relationship with bacterial pathogenicity revealed by a large-scale phylogenetic analysis. Microb Genom 7.

25. Chidwick, H.S., and Fascione, M.A. (2020). Mechanistic and structural studies into the biosynthesis of the bacterial sugar pseudaminic acid (Pse5Ac7Ac). Org Biomol Chem 18, 799–809.

26. Khairnar, A., Sunsunwal, S., Babu, P., and Ramya, T.N.C. (2020). Novel serine/threonine-O-glycosylation with N-acetylneuraminic acid and 3-deoxy-D-manno-octulosonic acid by bacterial flagellin glycosyltransferases. Glycobiology 31, 288–306.

27. Bubendorfer, S., Ishihara, M., Dohlich, K., Heiss, C., Vogel, J., Sastre, F., Panico, M., Hitchen, P., Dell, A., Azadi, P., et al. (2013). Analyzing the modification of the Shewanella oneidensis MR-1 flagellar filament. PLoS One 8, e73444.

28. Sulzenbacher, G., Roig-Zamboni, V., Lebrun, R., Guerardel, Y., Murat, D., Mansuelle, P., Yamakawa, N., Qian, X.X., Vincentelli, R., Bourne, Y., et al. (2018). Glycosylate and move! The glycosyltransferase Maf is involved in bacterial flagella formation. Environ Microbiol 20, 228–240.

29. Faulds-Pain, A., Birchall, C., Aldridge, C., Smith, W.D., Grimaldi, G., Nakamura, S., Miyata, T., Gray, J., Li, G., Tang, J.X., et al. (2011). Flagellin redundancy in Caulobacter crescentus and its implications for flagellar filament assembly. J Bacteriol 193, 2695–2707.

30. Montemayor, E.J., Ploscariu, N.T., Sanchez, J.C., Parrell, D., Dillard, R.S., Shebelut, C.W., Ke, Z., Guerrero-Ferreira, R.C., and Wright, E.R. (2021). Flagellar Structures from the Bacterium Caulobacter crescentus and Implications for Phage varphi CbK Predation of Multiflagellin Bacteria. J Bacteriol 203.

31. Ely, B., Ely, T.W., Crymes, W.B., Jr., and Minnich, S.A. (2000). A family of six flagellin genes contributes to the Caulobacter crescentus flagellar filament. Journal of bacteriology 182, 5001–5004.

32. Butaite, E., Baumgartner, M., Wyder, S., and Kummerli, R. (2017). Siderophore cheating and cheating resistance shape competition for iron in soil and freshwater Pseudomonas communities. Nat Commun 8, 414.

33. Marks, M.E., Castro-Rojas, C.M., Teiling, C., Du, L., Kapatral, V., Walunas, T.L., and Crosson, S. (2010). The genetic basis of laboratory adaptation in Caulobacter crescentus. J Bacteriol 192, 3678–3688.

34. Kenyon, J.J., Arbatsky, N.P., Shneider, M.M., Popova, A.V., Dmitrenok, A.S., Kasimova, A.A., Shashkov, A.S., Hall, R.M., and Knirel, Y.A. (2019). The K46 and K5 capsular polysaccharides produced by Acinetobacter baumannii NIPH 329 and SDF have related structures and the side-chain non-ulosonic acids are 4-O-acetylated by phage-encoded O-acetyltransferases. PLoS One 14, e0218461.

35. Shashkov, A.S., Senchenkova, S.N., Popova, A.V., Mei, Z., Shneider, M.M., Liu, B., Miroshnikov, K.A., Volozhantsev, N.V., and Knirel, Y.A. (2015). Revised structure of the capsular polysaccharide of Acinetobacter baumannii LUH5533 (serogroup O1) containing di-N-acetyllegionaminic acid. Russian Chemical Bulletin 64, 1196–1199.

36. Nedeljkovic, M., Postel, S., Pierce, B.G., and Sundberg, E.J. (2021). Molecular Determinants of Filament Capping Proteins Required for the Formation of Functional Flagella in Gram-Negative Bacteria. Biomolecules 11.

37. Khan, S.R., Gaines, J., Roop, n., R Martin, and Farrand, S.K. (2008). Broad-host-range expression vectors with tightly regulated promoters and their use to examine the influence of TraR and TraM expression on Ti plasmid quorum sensing. Appl Environ Microbiol 74, 5053–5062.

38. Ou, H.-Y., Kuang, S.N., He, X., Molgora, B.M., Ewing, P.J., Deng, Z., Osby, M., Chen, W., and Xu, H.H. (2015). Complete genome sequence of hypervirulent and outbreak-associated Acinetobacter baumannii strain LAC-4: epidemiology, resistance genetic determinants and potential virulence factors. Scientific Reports 5, 8643.

39. Takami, H., Takaki, Y., Chee, G.-J., Nishi, S., Shimamura, S., Suzuki, H., Matsui, S., and Uchiyama, I. (2004). Thermoadaptation trait revealed by the genome sequence of thermophilic Geobacillus kaustophilus. Nucleic Acids Research 32, 6292–6303.

40. Poehlein, A., Keyl, A., Milsch, J.C., and Daniel, R. (2018). Draft Genome Sequence of the Thermophilic Acetogen Moorella humiferrea DSM 23265. Genome Announcements 6, e00357–00318.

41. Breton, C., Šnajdrová, L., Jeanneau, C., Koča, J., and Imberty, A. (2005). Structures and mechanisms of glycosyltransferases. Glycobiology 16, 29R–37R.

42. Keys, T.G., and Aebi, M. (2017). Engineering protein glycosylation in prokaryotes. Current Opinion in Systems Biology 5, 23–31.

43. Cuccui, J., and Wren, B. (2015). Hijacking bacterial glycosylation for the production of glycoconjugates, from vaccines to humanised glycoproteins. J Pharm Pharmacol 67, 338–350.

44. Kay, E., Cuccui, J., and Wren, B.W. (2019). Recent advances in the production of recombinant glycoconjugate vaccines. npj Vaccines 4, 16.

45. Micoli, F., Adamo, R., and Costantino, P. (2018). Protein Carriers for Glycoconjugate Vaccines: History, Selection Criteria, Characterization and New Trends. Molecules 23.

46. Kenyon, J.J., and Hall, R.M. (2013). Variation in the complex carbohydrate biosynthesis loci of Acinetobacter baumannii genomes. PLoS One 8, e62160.

47. Parker, J.L., Lowry, R.C., Couto, N.A., Wright, P.C., Stafford, G.P., and Shaw, J.G. (2014). Maf-dependent bacterial flagellin glycosylation occurs before chaperone binding and flagellar T3SS export. Mol Microbiol 92, 258–272.

48. van Alphen, L.B., Wuhrer, M., Bleumink-Pluym, N.M.C., Hensbergen, P.J., Deelder, A.M., and van Putten, J.P.M. (2008). A functional Campylobacter jejuni maf4 gene results in novel glycoforms on flagellin and altered autoagglutination behaviour. Microbiology (Reading) 154, 3385–3397.

49. Nothaft, H., and Szymanski, C.M. (2010). Protein glycosylation in bacteria: sweeter than ever. Nature reviews. Microbiology 8, 765–778.

50. Harvey, H., Bondy-Denomy, J., Marquis, H., Sztanko, K.M., Davidson, A.R., and Burrows, L.L. (2018). Pseudomonas aeruginosa defends against phages through type IV pilus glycosylation. Nat Microbiol 3, 47–52.

51. Ely, B. (1991). Genetics of Caulobacter crescentus. Methods Enzymol. 204, 372–384.

52. Curtis, P.D., and Brun, Y.V. (2014). Identification of essential alphaproteobacterial genes reveals operational variability in conserved developmental and cell cycle systems. Mol Microbiol.

53. Thanbichler, M., Iniesta, A.A., and Shapiro, L. (2007). A comprehensive set of plasmids for vanillate- and xylose-inducible gene expression in Caulobacter crescentus. Nucleic Acids Research 35, e137.

