## Supplement tables for "Genetic glyco-profiling and rewiring of insulated flagellin glycosylation pathways"

**Table S1:** Comparison of the PseI/LegI synthases tested in this study to *Campylobacter jejuni* and *Caulobacter crescentus* proteins.

|  | ***Campylobacter jejuni NCTC11168*** | | | ***Caulobacter crescentus*** |
| --- | --- | --- | --- | --- |
|  | NeuB1 (Cj1141) | NeuB2 (Cj1327) | NeuB3 (Cj1317) | PseI (CCNA0296) |
| N-acetylneuraminate synthase | Sialic acid | Legionaminic acid | Pseudaminic acid | Pseudaminic acid |
| **Gram-negative** | | | | |
| *Caulobacter crescentus* | 32% (49%) | 35% (53%) | 43% (59%) | 100% |
| *Brevundimonas subvibrioides* | 30% (49%) | 42% (64%) | 32% (52%) | 36% (51%) |
| *Brevundimonas lutea* | 31% (50%) | 44% (66%) | 35% (51%) | 37% (55%) |
| *Brevundimonas viscosa* | 29% (47%) | 46% (63%) | 33% (50%) | 37% (49%) |
| *Magnetospirillum magneticum* | 31% (48%) | 36% (51%) | 45% (51%) | 56% (69%) |
| *Leptospira interrogans* | 31% (52%) | 35% (56%) | 44% (62%) | 55% (71%) |
| *Treponema pallidum* | 26% (44%) | 28% (43%) | 26% (41%) | 34% (49%) |
| *Treponema denticola* | 26% (46%) | 29% (47%) | 27% (46%) | 29% (47%) |
| *Acinetobacter baumannii ACICU* | 33% (51%) | 39% (57%) | 46% (62%) | 54% (67%) |
| *Acinetobacter baumannii LUH* | 30% (49%) | 46% (64%) | 30% (51%) | 30% (48%) |
| *Acinetobacter baumannii LAC-4* | 27% (46%) | 41% (62%) | 31% (51%) | 31% (47%) |
| *Legionella pneumophila subsp. Pneumophila Str. Philadelphia 1* | 32% (50%) | 46% (62%) | 34% (54%) | 34% (54%) |
| *Pseudomonas aeruginosa Y82* | 29% (46%) | 46% (60%) | 31% (47%) | 33% (47%) |
| *Pseudomonas aeruginosa PAPS475* | 34% (52%) | 36% (56%) | 46% (62%) | 59% (71%) |
| *Pseudomonas sp. Irchel 3A5* | 34% (51%) | 37% (54%) | 46% (62%) | 59% (71%) |
| *Shewanella oneidensis* | 27% (42%) | 32% (49%) | 31% (49%) | 30% (40%) |
| *Shewanella japonica* | 30% (49%) | 34% (54%) | 42% (60%) | 48% (63%) |
| **Gram-positive** | | | | |
| *Bacillus subtilis PY79* | 29% (48%) | 38% (52%) | 34% (51%) | 33% (51%) |
| *Kurthia* | 30% (51%) | 36% (56%) | 47% (63%) | 52% (69%) |
| *Kurthia sibirica* | 31% (53%) | 36% (56%) | 47% (65%) | 51% (68%) |
| *Clostridium botulinum* | 32% (54%) | 38% (60%) | 33% (52%) | 31% (51%) |
| *Geobacillus kaustophilus* | 34% (52%) | 50% (65%) | 36% (54%) | 31% (45%) |
| *Moorella humiferrea* | 35% (56%) | 54% (71%) | 38% (59%) | 38% (55%) |
| **Actinobacteria** | | | | |
| *Dermabacter vaginalis* | 28% (50%) | 34% (57%) | 40% (59%) | 51% (65%) |
| *Mycobacterium sp. KBS0706* | 31% (49%) | 37% (54%) | 46% (62%) | 66% (81%) |
| ***Archaea*** | | | | |
| *Halorubrum sp. PV6* | 32% (53%) | 43% (64%) | 36% (53%) | 35% (53%) |
| *Methanobrevibacter smithii* | 31% (51%) | 38% (57%) | 50% (70%) | 44% (63%) |

Percentage of identity and similarity (in parenthesis) between the different PseI/LegI/NeuB synthase homologs tested in this study. The enzymes that restore the motility of the *C. crescentus* Δ*pseI* cells and the *B. subvibrioides* Δ*legI* cells are highlighted in blue and in orange, respectively*.* The enzymes that do not restore motility are highlighted in white. *Caulobacter crescentus* (YP_002518334.1); *Brevundimonas subvibrioides* (WP_013267925); *Brevundimonas lutea* (WP_122465405.1); *Brevundimonas viscosa* (WP_092306097.1); *Magnetospirillum magneticium* (WP_011383156.1); *Leptospira interrogans* (WP_001033784.1); *Dermabacter vaginalis* (WP_223845738.1); *Treponema pallidum* (WP_010882009.1); *Treponema denticola* (WP_002682269.1); *Acinetobacter baumannii* ACICU (WP_000037963.1); *Acinetobacter baumannii* LUH (WP_071211480.1); *Acinetobacter baumannii* LAC-4 (WP_002063388); *Legionella pneumophila subsp. Pneumophila Str. Philadelphia* (Q5ZXH9.1); *Pseudomonas aeruginosa* Y82 (WP_003150869.1); *Pseudomonas aeruginosa* PASP475 (WP_171959540); *Pseudomonas sp. Irchel* (WP_095094799.1); *Shewanella oneidensis* (WP_011073144.1); *Shewanella japonica* (WP_080916269.1); *Kurthia* (WP_068455558.1); *Kurthia sibirica* (WP_109305561.1); *Clostridium botulinum* (WP_012100560.1); *Geobacillus kaustophilus* (WP_011232593.1); *Moorella humiferrea* (MBE3573281.1); *Mycobacterium sp. KBS0706* (WP_138891265); *Halorubrum sp. PV6* (WP_127117277.1); *Methanobrevibacter smithii* (WP_004033464.1).

**Table S2:** Comparison of PtmE/LegX orthologs to *Campylobacter jejuni* and *Brevundimonas subvibrioides* proteins.

|  | ***Campylobacter jejuni NCTC11168*** | ***Brevundimonas subvibrioides*** |
| --- | --- | --- |
|  | PtmE (Cj1329) | Bresu_3267 |
| **Gram-negative** | | |
| *Brevundimonas subvibrioides** | 36% (60%) | 100% (100%) |
| *Brevundimonas lutea** | 32% (60%) | 39% (59%) |
| *Brevundimonas viscosa** | 39% (59%) | 47% (62%) |
| *Treponema denticola* | No hit | No hit |
| *Acinetobacter baumannii LUH** | 39% (63%) | 42% (59%) |
| *Acinetobacter baumannii LAC-4** | 37% (63%) | 41% (60%) |
| *Legionella pneumophila subsp. Pneumophila Str. Philadelphia 1* | No hit | 30% (42%)$ |
| *Pseudomonas aeruginosa Y82** | 34% (61%) | 41% (61%) |
| **Gram-positive** | | |
| *Bacillus subtilis PY79* | 24% (45%)$ | 24% (46%)$ |
| *Clostridium botulinum** | 38% (65%) | 34% (59%) |
| *Geobacillus kaustophilus** | 41% (67%) | 44% (63%) |
| *Moorella humiferrea** | 33% (56%) | 36% (54%) |
| ***Archaea*** | | |
| *Halorubrum sp. PV6* | 28% (49%)$ | 34% (46%)$ |

Percentage of homology and similarity (in bracket) between the different nucleotidyltransferases present in the species that harbor proven/predicted legionaminic acid synthases used in this study. The enzymes that restore the motility of the *B. subvibrioides* Δ*legI* cells are highlighted in orange*.* The enzymes that do not restore motility are highlighted in white. The asterisk (*) indicates that the nucleotidyltransferase-encoding gene is genetically clustered with genes coding for enzymes predicted to be involved in legionaminic acid production. The dollar symbol ($) indicates the corresponding sequence similarity of less than 65% of the protein length. When the best-hits are different between the two species they are separated with the slash. *Brevundimonas subvibrioides* (WP_013270673.1), *Brevundimonas lutea* (WP_122465403.1), *Brevundimonas viscosa* (WP_092306095.1), *Acinetobacter baumannii LUH* (AHB32582.2), *Acinetobacter baumannii LAC-4* (AIY39058.1), *Legionella pneumophila subsp. Pneumophila Str. Philadelphia 1* (WP_010946056.1), *Pseudomonas aeruginosa Y82* (WP_003150871.1), *Bacillus subtilis* PY79 (WP_003244408.1 /WP_009968049.1), *Clostridium botulinum* (WP_003359147.1), *Geobacillus kaustophilus* (WP_011232591.1), *Moorella humiferrea* (WP_106005998.1), *Halorubrum sp. PV6* (WP_127117239.1/ WP_127117240.1).

**Table S3: Strains and plasmids used in this study.**

| **Strains** | **Relevant characteristics** | **Origins** |
| --- | --- | --- |
| ***C. crescentus*** |  |  |
| NA1000 | Synchronizable derivative of wild type strain CB15 | (Evinger and Agabian 1977) |
| Δ*neuB* | NA1000 derivative with in-frame deletion of *neuB* | (Ardissone et al. 2020) |
| Δ*flmG* | NA1000 derivative with in-frame deletion of *flmG* | (Ardissone et al. 2020) |
| Δ*fl^jx6^* | NA1000 derivative with in-frame deletions of *fljJ*, *fljK*, *fljL* and *fljMNO* | (Faulds-Pain et al. 2011) |
| Δ*fl^jx6^ ΔneuB ΔflmG* | NA1000 derivative with in-frame deletions of *fljJ, fljK, fljL, fljMNO, neuB* and *flmG* | This work |
| ***B. subvibrioides*** |  |  |
| ATCC15264 | Wild-type |  |
| Δ*neuB* | ATCC15264 derivative with in frame-deletion of *neuB* (Bresu_0507) | This work |
| Δ*flmG* | ATCC15264 derivative with in frame-deletion of *flmG* (*Bresu_2406*) | This work |
| Δ*flmG* Δ*neuB* | ATCC15264 derivative with in frame-deletion of *neuB* and *flmG* | This work |
| Δ*Bresu_3266* | ATCC15264 with in frame-deletion of *Bresu_3266* | This work |
| Δ*Bresu_3267* | ATCC15264 with in frame-deletion of *Bresu_3267* | This work |
| Δ*Bresu_0506* | ATCC15264 with in frame-deletion of *Bresu_0506* | This work |
| ***E. coli*** |  |  |
| EC100D | Cloning strain | Epicentre Technologies |
| S17.1 λ*pir* | For plasmid mobilization | (Simon et al. 1983) |
| **Plasmids** | **Relevant characteristics** | **Origins** |
| pNPTS138 | Two-part selection in in-frame deletion vector, *oriT*, *sacB*, Kan^R^ | Alley MRK, unpublished |
| pSRK-Gm | pBBR1MCS-5 derived broad host range vector containing *lac* promoter, *lacI*^q^, l*acZ*α^+^; Gm^R^ | (Khan et al. 2008) |
| pMT463 | Medium copy number plasmid for inducible expression; P*_xyl_*, Gm^R^ | (Thanbichler et al. 2007) |
| pMT375 | Low copy number plasmid for inducible expression; P_xyl_, Tet^R^ | (Thanbichler et al. 2007) |
| pMT335 | High copy number plasmid for inducible expression; P_van_, Gm^R^ | (Thanbichler et al. 2007) |
| pXGFP4 | Integrative plasmid for inducible expression; P_xyl_, KanR | (Thanbichler et al. 2007) |
| pSA37 | pNPTS138 derivative carrying the in-frame deletion of *flmG* for *C. crescentus,* Kan^R^ | (Ardissone et al. 2020) |
| pNK580 | pNPTS138 derivative carrying the in-frame deletion of *neuB* for *B. subvibrioides*, Kan^R^ | This work |
| pNK562 | pNPTS138 derivative carrying the in-frame deletion of *flmG* for *B. subvibrioides*, Kan^R^ | This work |
| pNK926 | pNPTS138 derivative carrying the in-frame deletion of *Bresu_3266* for *B. subvibrioides*, Kan^R^ | This work |
| pNK1000 | pNPTS138 derivative carrying the in-frame deletion of *Bresu_3267* for *B. subvibrioides*, Kan^R^ | This work |
| pNK1002 | pNPTS138 derivative carrying the in-frame deletion of *Bresu_0506* for *B. subvibrioides*, Kan^R^ | This work |
| pNK948 | pSRK derivative carrying the *B. subvibrioides Bresu_3266* ORF, Gm^R^ | This work |
| pNK950 | pSRK derivative carrying the *B. subvibrioides Bresu_3267* ORF, Gm^R^ | This work |
| pNK988 | pSRK derivative carrying the *B. subvibrioides Bresu_3266* and *Bresu_3267* ORFs, Gm^R^ | This work |
| pNK974 | pSRK derivative carrying the *B. subvibrioides Bresu_0506* ORF, Gm^R^ | This work |
| pNK957 | pMT375 derivative carrying the *B. subvibrioides Bresu_3267* ORF, Tet^R^ | This work |
| pSA228 | pMT463 derivative carrying the synthetic *B. subvibrioides fljK* (*Bresu_2638*) ORF codon optimized for *E. coli*, Gm^R^ | This work |
| pSA462 | pMT463 derivative carrying the synthetic *B. subvibrioides fljK* ORF codon optimized for *E. coli* and the *B. subvibrioides flmG* ORF, Gm^R^ | This work |
| pLT2295 | pXGFP4 derivative carrying the synthetic *B. subvibrioides Bresu_3266*, *Bresu_0765*, *Bresu_0506*, *Bresu_3264*, *neuB* (*Bresu_0507*) and *Bresu_3265* ORFs codon optimized for *E. coli*, Kan^R^ | This work |
| pNK955 | pSRK derivative carrying the synthetic *M. humiferrea* *MOHU_20790* ORF codon optimized for *E. coli*, Gm^R^ | This work |
| pNK660 | pSRK derivative carrying the *B. subvibrioides flmG* ORF, Gm^R^ | This work |
| pSA96 | pSRK derivative carrying the *C. crescentus* *flmG* ORF, Gm^R^ | (Ardissone et al. 2020) |
| pNK628 | pSRK derivative carrying the synthetic *L. interrogans neuB* ORF codon optimized for *E. coli*, Gm^R^ | This work |
| pNK631 | pSRK derivative carrying the synthetic *B. subvibrioides neuB* ORF codon optimized for *E. coli*, Gm^R^ | This work |
| pNK649 | pSRK derivative carrying the synthetic *flmG^Cc-Bs^* chimera ORF codon optimized for *E. coli*, Gm^R^ | This work |
| pNK661 | pSRK derivative carrying the synthetic *M. smithii neuB* ORF codon optimized for *E. coli*, Gm^R^ | This work |
| pNK663 | pSRK derivative carrying the synthetic *B. viscosa neuB* ORF codon optimized for *E. coli*, Gm^R^ | This work |
| pNK665 | pSRK derivative carrying the synthetic *B. lutea neuB* ORF codon optimized for *E. coli*, Gm^R^ | This work |
| pNK668 | pSRK derivative carrying the synthetic *M. humiferrea neuB* ORF codon optimized for *E. coli*, Gm^R^ | This work |
| pNK663 | pSRK derivative carrying the synthetic *B. viscosa neuB* ORF codon optimized for *E. coli*, Gm^R^ | This work |
| pNK707 | pSRK derivative carrying the synthetic *P. aeruginosa PASP475 neuB* ORF codon optimized for *E. coli*, Gm^R^ | This work |
| pNK708 | pSRK derivative carrying the synthetic *A. baumannii LAC-4 neuB* ORF codon optimized for *E. coli*, Gm^R^ | This work |
| pNK709 | pSRK derivative carrying the synthetic *K. sibirica neuB* ORF codon optimized for *E. coli*, Gm^R^ | This work |
| pNK710 | pSRK derivative carrying the synthetic *S. japonicum neuB* ORF codon optimized for *E. coli*, Gm^R^ | This work |
| pNK711 | pSRK derivative carrying the synthetic *Mycobacterium sp. neuB* ORF codon optimized for *E. coli*, Gm^R^ | This work |
| pNK715 | pSRK derivative carrying the synthetic *A. baumannii ACICU neuB* ORF codon optimized for *E. coli*, Gm^R^ | This work |
| pNK716 | pSRK derivative carrying the synthetic *P. aeruginosa Y82 neuB* ORF codon optimized for *E. coli*, Gm^R^ | This work |
| pNK720 | pSRK derivative carrying the synthetic *A. baumannii LUH5551 neuB* ORF codon optimized for *E. coli*, Gm^R^ | This work |
| pNK735 | pSRK derivative carrying the synthetic *T. pallidum neuB* ORF codon optimized for *E. coli*, Gm^R^ | This work |
| pNK833 | pSRK derivative carrying the synthetic *L. pneumophila neuB* ORF codon optimized for *E. coli*, Gm^R^ | This work |
| pNK835 | pSRK derivative carrying the synthetic *T. denticola neuB* ORF codon optimized for *E. coli*, Gm^R^ | This work |
| pNK837 | pSRK derivative carrying the synthetic *Halorubrum sp. PV6* *neuB* ORF codon optimized for *E. coli*, Gm^R^ | This work |
| pNK900 | pSRK derivative carrying the synthetic *G. kaustophilus neuB* ORF codon optimized for *C. crescentus*, Gm^R^ | This work |
| pNK915 | pSRK derivative carrying the synthetic *C. botulinum neuB* ORF codon optimized for *E. coli*, Gm^R^ | This work |
| pNK934 | pSRK derivative carrying the synthetic *B. subtilis PY79 neuB* ORF codon optimized for *E. coli*, Gm^R^ | This work |
| pNK629 | pSRK derivative carrying the synthetic *D. vaginalis neuB* ORF codon optimized for *E. coli*, Gm^R^ | This work |
| pNK991 | pSRK derivative carrying the *C. jejuni neuB1*/*sia* ORF, Gm^R^ | This work |
| pNK992 | pSRK derivative carrying the *C. jejuni neuB2*/*legI* ORF, Gm^R^ | This work |
| pNK994 | pSRK derivative carrying the *C. jejuni neuB3*/*pseI* ORF, Gm^R^ | This work |
| pSA53 | pMT335 derivative carrying the *C. crescentus* *neuB* ORF | (Ardissone et al. 2020) |
| pLT2237 | pMT335 derivative carrying the *Kurthia neuB* ORF, Gm^R^ | This work |
| pSA126 | pMT335 derivative carrying the *C. jejuni neuB1*/*sia* ORF, Gm^R^ | (Ardissone et al. 2020) |
| pSA47 | pMT335 derivative carrying the *C. jejuni neuB2*/*legI* ORF, Gm^R^ | (Ardissone et al. 2020) |
| pSA48 | pMT335 derivative carrying the *C. jejuni neuB3*/*pseI* ORF, Gm^R^ | (Ardissone et al. 2020) |
| pLT2262 | pMT335 derivative carrying the *M. magneticum neuB* ORF, Gm^R^ | This work |
| pLT2263 | pMT335 derivative carrying the *S. oneidensis neuB* ORF, Gm^R^ | This work |
| pLT2043 | pMT335 derivative carrying the *P. irschel 3A5* ORF, Gm^R^ | This work |
| pLT2036 | pMT335 derivative carrying the *B. subvibrioides neuB* ORF, Gm^R^ | This work |

Ardissone, S., N. Kint, and P. H. Viollier. 2020. Specificity in glycosylation of multiple flagellins by the modular and cell cycle regulated glycosyltransferase FlmG. Elife **9**.

Evinger, M., and N. Agabian. 1977. Envelope-associated nucleoid from Caulobacter crescentus stalked and swarmer cells. J Bacteriol **132**:294-301.

Faulds-Pain, A., C. Birchall, C. Aldridge, W. D. Smith, G. Grimaldi, S. Nakamura, T. Miyata, J. Gray, G. Li, J. X. Tang, K. Namba, T. Minamino, and P. D. Aldridge. 2011. Flagellin redundancy in Caulobacter crescentus and its implications for flagellar filament assembly. J Bacteriol **193**:2695-2707.

Khan, S. R., J. Gaines, R. M. Roop, 2nd, and S. K. Farrand. 2008. Broad-host-range expression vectors with tightly regulated promoters and their use to examine the influence of TraR and TraM expression on Ti plasmid quorum sensing. Appl Environ Microbiol **74**:5053-5062.

Simon, R., U. Priefer, and A. Pühler. 1983. A broad host range mobilization system for in vivo genetic engineering: Transposon mutagenesis in Gram negative bacteria. Bio/Technology **1**:784-791.

Thanbichler, M., A. A. Iniesta, and L. Shapiro. 2007. A comprehensive set of plasmids for vanillate- and xylose-inducible gene expression in Caulobacter crescentus. Nucleic Acids Research **35**:e137.

Table S4. Oligonucleotides and synthetic genes used in this study

| **Primer** | **Sequence 5’-3’ (Restriction sites underlined)** |
| --- | --- |
| Bs_flmG_del_1 | aaaaaaCAATTGTGCATCAGCGCGCGTGTGA |
| Bs_flmG_del_2 | aaaaaaGGATCCGGTGTCGATGGCCTTTCGCTGGGCCTT |
| Bs_flmG_del_3 | aaaaaaGGATCCTCCTGTGCGCTGGACCTGGTGGT |
| Bs_flmG_del_4 | aaaaaaAAGCTTGAGACGGGGTCGCTGGCGGGAAA |
| Bs_neuB_del_1 | aaaaaaGAATTCGGGTGTTCGGCAAGCCGCTGGAAGA |
| Bs_neuB_del_2 | aaaaaaGGATCCGAAGGTCTGGAACTTGCGGCGT |
| Bs_neuB_del_3 | aaaaaaGGATCCTTCGCCGCGCGCGACATCGCGGCA |
| Bs_neuB_del_4 | aaaaaaAAGCTTCGCGGATCGGAAAGGACGGGCGA |
| NK339 | aaaaaaGGATCCGTCCGGTCGCCCTCCATG |
| NK340 | aaaaaaAAGCTTCACGAACCCGCGGGTCG |
| NK341 | aaaaaGGATCCCCCGAGGAGTATGCGGTG |
| NK342 | aaaaaaGAATTCGTCGCCGTTCAGGACCAC |
| NK345 | aaaaaaAAGCTTCGTACGAGGCCCCGGAG |
| NK346 | aaaaGGTACCGCGCGGCTCACACCGC |
| NK347 | aaaaaaGGTACCGACGGCCGGACCGACTG |
| NK348 | aaaaaaGAATTCCGATGATGTGGTCCGCCC |
| NK357 | aaaaaaCATATGGAGGGCGACCGGACGAC |
| NK358 | aaaaaaTCTAGATCACACCGCATACTCCTCG |
| NK359 | aaaaaaCATATGGTGAGCCGCGCCGATCC |
| NK360 | aaaaaaTCTAGATCAGTCGGTCCGGCCGTC |
| NK361 | aaaaaaCATATGGGCTGGCATGATGACAAGTTAG |
| NK366 | aaaaaaAAGCTTGATGCCGTATCGCGAGAAG |
| NK367 | aaaaaaGGATCCGTGCCTCGACATGGCCC |
| NK368 | aaaaaaGGATCCCAATGACCGAGCGGGTCTTC |
| NK369 | aaaaaaGAATTCCCAGGGCGTTGTCGAGG |
| NK374 | aaaaaaCATATGTCGAGGCACGGGCCCAC |
| NK375 | aaaaaaTCTAGATCATTGAGCCTCCAGAGGAC |
| Bs_flmG_rbs_Eco | GAATTCAGGAGGTAAAAAAATGACCCACGTCACCCCCT |
| Bs_flmG_Xba | aaaaaaTCTAGATTAGGCGGCGTTCCCGACCT |
| Bs_flmG_NdeI | aaaaaaCATATGACCCACGTCACCCCCT |
| 3A5_PseI_ndeI | aaaaaaCATATGAGTCCATCGATATCCATAGCA |
| 3A5_PseI_mfeI | aaaaaaCAATTGTCACTGTTTGAGCTTCACATTCT |
| Bs_neuB_nde | aaaaaaCATATGACCGAGCGGGTCTTCTTCAT |
| Bs_neuB_eco | aaaaaaGAATTCAGACCAGGTCGGTCGGCGCCAGCTT |
| Ku_neuB_nde | aaaaaaCATATGACAATTAAAGTAGAACAATATGAAA |
| Ku_neuB_eco | aaaaaaGAATTCTTAAATTAAGTCGTCAAATGTAATA |
| So_neuB_nde | aaaaaaCATATGACTAATCATTCCCCCCTATTTATC |
| So_neuB_eco | aaaaaaGAATTCATTCTGATACTCCAAACTTTTCTCTTTG |
| Mm_neuB_nde | aaaaaaCATATGACCGAGATTTCCATCGCCGGA |
| Mm_neuB_eco | aaaaaaGAATTCTACTCGATCATCGACCAGTCGAA |
| **Synthetic DNA** (from Integrated DNA Technologies, Coralville, Iowa, USA) | |
| *B. subvibrioides fljK*, codon optimised for *C. crescentus* (5’-3’) | CATATGGCGCTGAACTCGGTCAACACCAACTCCGGCGCCCTGGTGGCCCTGCAGAACCTGCAGTCGACCAACTCCGAACTGGCCACCGTCCAGTCCCGCATCAACACGGGCAAGAAGGTGAACTCGGCCAAGGATAACGGCGCGGTCTGGGCCATCGCCCAGGGCCAGCGCTCGGAGGTGAATGCCCTGGGCGCCGTGAAGGATAGCCTGGCCCGTGGCTCCTCCGCCGTGGACGTGAGCATCGCCGCGGGCGAATCGGTGTCCGACCTGCTGCTCCAGCTGAAGGAGAAGGCGCTGAGCGCGACCGACAAGAGCCTGACGACGGCGGCCCGGACGGCCCTGAACGAAGACTTCAAGGCCATCCGCGACCAGATCACCACCGTCGTGACCAACGCCAAGTTCAACGGCGTCAACCTGCTGGACAATAGCACCGGCACCGGGGGCTACAAGGCGCTCTCGAACACCGCCGGCTCGACCATCAAGGTCGCCGGCGAAAACCTGTCGCTGGGCGGCACGAACGTCACCGTCGCCACGACGACCACCATCGGCACCAGCACGCTGGCGACCACCGCGCTGGGTCTGGTCAATGCCAGCATCGACAAGGTGTCGGCCTCCCTGGCCCGCCTGGGCACGGGTGCGAAGGCCCTGGACACCCACAGCACGTTCGTCGGCAAGCTGAGCGACGCGCTGGAGAACGGTATCGGCAACCTGGTCGACGCGGACCTGGCGAAGGAGTCCGCGCGCCTGCAGTCGCTGCAGACCAAGCAGCAGCTGGGGGTGCAGGCGCTGTCGATCGCCAACCAGTCGAGCTCGATCCTGCTGGGCCTGTTCCGCTGAATTCTAGA |
| *B. subvibrioides neuB,* codon optimised *for E. coli* (5’-3’) | AGATCTGCTAGCCCAGCCACAGGCCCGTGCCGGGATCGAAGGCGAAGCGGCAGCCGATCAGGCGGAACTGGGCGCGGGCCATGGTCTCGAACAGGGCCGTCAGGTCGCGGGCGGCGTCCAGGTCGTCGTGGTCCAGCACCAGGCAGCGGATCTGCAGCCAGCCGTGGTCGGGCAGCAGGTAGAAGGCGCCCTCGTCCTGATCCTCGCCCGAAACCTCCAGCCCCCGGTCGATGGCTTCGACGACATAGCCGGCCGCGCGGCAGGTGTCGGTGAGCGCGGCCAGCAGGGCGGCTTCCTGGTCAGGGGTCAGGTCGGTCATGGGCAAGAGGTCCAGGTCGTGGTTTGTCGGCGGCTTCTAGCATGGACCGCCCGCGCCCGTGAGGCCGAGGATTTCGCGCTGGTCAGACAACCTACTTGCCGTCCCCACATGTTAGCGCTACCAAGTGCCGACGAACGCGCGCCGCCGACGGTGTCGGCGCTTCAGACGCTCGAGTTTTGGGGAGACGACGCCATATGACGGAGCGTGTTTTCTTCATCGCGGAGGCAGGTGTCAACCATAATGGGGACTTAGACCTGGCACTTCGCTTAGTGGACGTAGCTCATCGCGCCGGTGCAGACGCTGTAAAGTTTCAGACCTTTACCACCGATCGTGTAATTGCAGTGGATGCACCGCTTGCTGAGTACCAAAAAGCCAACGCAGGAGCACCGTCAGCGGTGGAGATGCTGAAAGGATTCGAACTTCCACGTGCTGATTTTGCCCGCATCGCCGACCATTGTCGTGCTATTGGCATTGAGTTTATGTCATCACCCTTCGACTTGGAGTCAGCGGCTTTCCTGGCCGGGATTGGGATGACTAAATTCAAACTTGCCTCGGGCGAGATTACTCACGCTCCATTTGTCCGTGGAGTGGCTCGCTTGGCAGCTGACGCTGGTGGGAGCGTGATCTTGTCAACCGGAATGTCTACCTTGGATGAGGTCGCGGGCGCGGTCGGCTGGATCGAGGGTGAAGGTTGTCGTGATTTGACAATCCTTCATTGCGTGTCAAATTACCCTGCTCCCGCAAAGGACGCGAACTTACGCGCCATGGACACGCTTCGCGAAGCCTTCGGTCATCCTGTGGGATGGTCCGACCACACATTAGGTGACGCAGTGTCGTTAGCCGCCGTAGCCCGCGGCGCCCGTGTCGTTGAAAAACATTTTACCCTTGACAACGCACTTGCAGGTCCCGATCATGCAATGTCCATGGACCCCGACGGGTTAGCACGTTTGATTGCGGGCCTTCGTACAGTTGAGGCGTCACTTGGCGACGGGATTAAACGCCCAGTGGAAGCAGAACGTGAAATCATGACAGTCGCCCGCCGTTCGTTATTTGCAGCGCGTGACATCGCCGCAGGTGCGGTGGTGACTGAAGCAGACTTAATCGCATTGCGCCCCGGCGATGGCATTAGCGCGGCTCGCCATGCTGACGTAGTTGGTCGCACAGCCGCACGTGCTTTACCCGCAGGGACCAAGTTAGCGCCGACTGACTTGGTATGAAACTAGTACTTAAGAAGGAGAGAATTCATGTCTAGAAAAGAGCTCGCGGCCGCGGATCC |
| *B. lutea neuB,* codon optimised for *E. coli* (5’-3’) | AGATCTGCTAGCCCAGCCACAGGCCCGTGCCGGGATCGAAGGCGAAGCGGCAGCCGATCAGGCGGAACTGGGCGCGGGCCATGGTCTCGAACAGGGCCGTCAGGTCGCGGGCGGCGTCCAGGTCGTCGTGGTCCAGCACCAGGCAGCGGATCTGCAGCCAGCCGTGGTCGGGCAGCAGGTAGAAGGCGCCCTCGTCCTGATCCTCGCCCGAAACCTCCAGCCCCCGGTCGATGGCTTCGACGACATAGCCGGCCGCGCGGCAGGTGTCGGTGAGCGCGGCCAGCAGGGCGGCTTCCTGGTCAGGGGTCAGGTCGGTCATGGGCAAGAGGTCCAGGTCGTGGTTTGTCGGCGGCTTCTAGCATGGACCGCCCGCGCCCGTGAGGCCGAGGATTTCGCGCTGGTCAGACAACCTACTTGCCGTCCCCACATGTTAGCGCTACCAAGTGCCGACGAACGCGCGCCGCCGACGGTGTCGGCGCTTCAGACGCTCGAGTTTTGGGGAGACGACGCCATATGGCGCAAAGTGATGTATTCATTATTGCTGAGGCGGGTGTAAATCACAATGGCGATATGGATTTGGCGCGTCGCTTGATTGATGTGGCTGCCGAAGCAGGCGCTGACGCGGTAAAATTCCAAACTTTCACGAGTGCCTCCCTGATGAGCGATGCCGCTCCTATGGCGGAATATCAGAAGGCCGGTGCTGAGGCCGATAGTGCGGTGGAAATGATTCGTGCCCTTGAGCTTGGACATGACCAATTTGCGGAATTGGCGCGCCATTGCGAGCAACGCGGTATCTTATTCCTTAGCACCGCGTTCGACATTGATTCGGCCCGCTTCTTGCATGGTTTAGGGATGCGTCGCTTTAAGATCCCTTCCGGAGAAATTACTAATGCACCTCTTATTCGCCGTATCGCGCGTATGGCGGATCAAGTTATCGTATCAACAGGAATGGCGACTTTGGCCGATGTGGAAGCAGCCGTTGGATGGATTGAGGCAGAAGGAGTTGAAGACATCGTAATTCTTCACTGCGTAAGTAATTACCCGGCGCCACCAGAAGCGGCAAATCTTCGTGCCATGGACACTTTACGCGCTGCCTTCGGACATCCCGTCGGGTGGTCAGACCACACGTTAGGTGACGCTGTTACCTTAGCCGCTGTGGCACGCGGCGCAACGGTGATCGAAAAACATTTCACCTTGGATACAGCACTGCCAGGGCCCGATCATGCAATGAGTTTGAATCCGGATGAACTTCGTGCGATGGTAGCCCGCATTCGCGACGTAGAAGCGTCATTGGGCTCCGGTCGCAAGGCTCCCACGGAGGCGGAACGTGAGACCGCAGTTGTCGCTCGCCGCTCTCTGTTTGCCCGTGTCGCTATTGCTGCTGGACAGACGGTAACGGAGGAAGAGTTGATCGCGTTACGCCCGGGAGACGGTCTGTCTCCAGCCCGTCTGGACGACATCGTCGGCCGCCCGGCTGCTCGTGACATTGCTGCCGGCGCAAAGCTTGGATGGAATGATTTAGCGTGAAACTAGTACTTAAGAAGGAGAGAATTCATGTCTAGAAAAGAGCTCGCGGCCGCGGATCC |
| *B. viscosa neuB,* codon optimised for *E. coli* (5’-3’) | AGATCTGCTAGCCCAGCCACAGGCCCGTGCCGGGATCGAAGGCGAAGCGGCAGCCGATCAGGCGGAACTGGGCGCGGGCCATGGTCTCGAACAGGGCCGTCAGGTCGCGGGCGGCGTCCAGGTCGTCGTGGTCCAGCACCAGGCAGCGGATCTGCAGCCAGCCGTGGTCGGGCAGCAGGTAGAAGGCGCCCTCGTCCTGATCCTCGCCCGAAACCTCCAGCCCCCGGTCGATGGCTTCGACGACATAGCCGGCCGCGCGGCAGGTGTCGGTGAGCGCGGCCAGCAGGGCGGCTTCCTGGTCAGGGGTCAGGTCGGTCATGGGCAAGAGGTCCAGGTCGTGGTTTGTCGGCGGCTTCTAGCATGGACCGCCCGCGCCCGTGAGGCCGAGGATTTCGCGCTGGTCAGACAACCTACTTGCCGTCCCCACATGTTAGCGCTACCAAGTGCCGACGAACGCGCGCCGCCGACGGTGTCGGCGCTTCAGACGCTCGAGTTTTGGGGAGACGACGCCATATGTCCCGTGTCTTCGTAATTGCTGAAGCAGGCGTCAATCACGACGGGAGCCTTGATGACGCTCTACGTATGGTTGACGTAGCAGCTGAAGCCGGTGCCGACGCCGTCAAATTTCAGACCTTTGACGCAGCTCAACTGGTAACTCGTCGTGCAGCGAAAGCGGCTTATCAGGCACGCAATACAGGGGCGGATGATGGGCAGTTAGCGATGTTGCGCCGTCTTGAATTAGACCGTGACGCCCACTGCACTCTGGCCCGCGCGGCCGAGGCCAAAGGGGTACGCTTTATGTCAACTGCTTTCGACATGGATTCCCTGGACTTTCTGGCAGGTCTGGACATGCCCGCTATCAAAATCCCATCCGGAGACTTAACATGGGGACCGATGTTGCTCAAGGCCGCTCGTTTAGGGCTTCCACTTATCGTAAGCACTGGTATGGCAACACTGGGAGAGATTGAGGAGGCTTTACAGGTTATCGCGTTTGCACTTACTCGTGACGGGCTTCCGGCCGGAAGTCAAGAGCTTAAGGCCGCTTTTGCAGAGCCACAAGCTCAAGCAGCCTTACGTGATCGCGTAACGTTATTGCATTGCACTACAGAGTATCCCGCTCCCCTTCGCACTGTAAACCTGCGTGCCATGGATCTGATGCGTGAGACTTTTGGTTTACCGGTAGGTTACTCGGATCATACTCTTGGGACCACCGTGGCGATCGCGGCGGCTGCTCGCGGCGCTACGGTCGTCGAAAAACACTTCACGCTTGATCGTGCTCGTCCCGGTCCTGACCACGCCGCATCGCTGGAACCGGACGAACTTGCCGCAATGGTTGCTGCCGTTCGTGAGGTAGAGGCTGCACTTGGTGAGGCACGTAAAGGTCCGGCAGCTGAGGAAATCAGTAATCGCGCCATCGCCCGCCGTAGCTTGGTGGCTGCTCGTCCAGTGGCTGCCGGGGAGCCCTTCTCTCTGGACAACTTGACTGCAAAGCGTCCCGCGGACGGATTGTCACCGATGGAAGTCTGGCCCCTTCTTGGACAACCAGCTGCTCGTGACTGGGCAGAAGACGAGGCGATCACCGCTTGAAACTAGTACTTAAGAAGGAGAGAATTCATGTCTAGAAAAGAGCTCGCGGCCGCGGATCC |
| *T. pallidum neuB,* codon optimised for *E. coli* (5’-3’) | CAATTGAGATCTGCTAGCCCAGCCACAGGCCCGTGCCGGGATCGAAGGCGAAGCGGCAGCCGATCAGGCGGAACTGGGCGCGGGCCATGGTCTCGAACAGGGCCGTCAGGTCGCGGGCGGCGTCCAGGTCGTCGTGGTCCAGCACCAGGCAGCGGATCTGCAGCCAGCCGTGGTCGGGCAGCAGGTAGAAGGCGCCCTCGTCCTGATCCTCGCCCGAAACCTCCAGCCCCCGGTCGATGGCTTCGACGACATAGCCGGCCGCGCGGCAGGTGTCGGTGAGCGCGGCCAGCAGGGCGGCTTCCTGGTCAGGGGTCAGGTCGGTCATGGGCAAGAGGTCCAGGTCGTGGTTTGTCGGCGGCTTCTAGCATGGACCGCCCGCGCCCGTGAGGCCGAGGATTTCGCGCTGGTCAGACAACCTACTTGCCGTCCCCACATGTTAGCGCTACCAAGTGCCGACGAACGCGCGCCGCCGACGGTGTCGGCGCTTCAGACGCTCGAGTTTTGGGGAGACGACGCCATATGTTCACGTGCGGAGGCCGTTGCTTTCGCCCCGACGCCGACATCTTAACCATTGCGGAAATCGGCTCAGCTCACGCCGGGTCTTTCGACCGCGCACGTGCATTAATCGATGCAGCGGCTGATGCTGCTGCGGCTGCCGTCAAATTCCAGCTGATTTACGCACACGAAATCTTACACCCTTTAACCGGGGCTGTGCGCCTGCCCTCAGGCGCGGTTTCGTTATACCAACGCTTTGAAGAACTTGAAGTGCCTTTATCCTTCTATGCACAGTGCTTCAACCACGCACGCAGCCGCGGCATGCTGGTCGGGATCAGCCCATTCGGACCACGTTCCGCCACGGAAGCACTTGCGTTGAAACCGGACTTCTTAAAAGTAGCCTCACCTGAGTTGAATTATCCGACGCTGATTAGTACGTTAGCTGCTGCTGAGTTGCCTCTTATTTTGTCATCCGGCGTTTGTCTGCTGAAAGAAATCGAGGGAGCGTTGGCGCAGTGCCGTCAGTACACCAAACAGGGGTCGAGTCATGCATTACTTCACTGTATCACAGCGTACCCAGCACCTGAGACGGAGTACAATTTGGCATTGTTACCCGCCTTGGCAACTATTTTCAACATCAACGTGGGTGTGAGTGACCATAGCGTAGACCCTTTATTAGTCCCTTTATTAGCACGCGCCCACGGCGCATGTATCGTAGAGAAACATATTTGTCTGAGCCGCACCGATGCGGGGCTTGACGACAGTATCGCTCTGGACCCTGCTGATTTTCGCACAATGACTGCGGCGTTAAACTCATGTGCGCGCCGCAGTCCGTCGCAGATTATTAGCTTTTTACACGAACGTGGGTATGCTCCCCACGTTGTGCGCGCTGTAATCGGCTCAGGGGAAAAAGTATTGGCGCCCTCAGAGCGCGCACACTACCAGAAGTCCAATCGCTCCTTGCACTATTTGCACGCCTACCCTCGCGGGACCGTCTTACAGAAGGAAAACCTGGTCATCGTACGTAGTGAAGCGAACCTGAGCGCCGGTGAAGCTCCCGAGCATGCAAATTTATTTGTTGGAGCTGTATTACAGCGTTCGGTCCACGCAGGTGAGGGTGCGCGCTTTGGCGACATTATCCAGAAGGGCACTTGTTTATGAGCTCGAATTCTAGAGCGGCCGCATACGTA |
| *T. denticola neuB,* codon optimised for *E. coli* (5’-3’) | GGATCCCAATTGAACTAGTACCAGCCACAGGCCCGTGCCGGGATCGAAGGCGAAGCGGCAGCCGATCAGGCGGAACTGGGCGCGGGCCATGGTCTCGAACAGGGCCGTCAGGTCGCGGGCGGCGTCCAGGTCGTCGTGGTCCAGCACCAGGCAGCGGATCTGCAGCCAGCCGTGGTCGGGCAGCAGGTAGAAGGCGCCCTCGTCCTGATCCTCGCCCGAAACCTCCAGCCCCCGGTCGATGGCTTCGACGACATAGCCGGCCGCGCGGCAGGTGTCGGTGAGCGCGGCCAGCAGGGCGGCTTCCTGGTCAGGGGTCAGGTCGGTCATGGGCAAGAGGTCCAGGTCGTGGTTTGTCGGCGGCTTCTAGCATGGACCGCCCGCGCCCGTGAGGCCGAGGATTTCGCGCTGGTCAGACAACCTACTTGCCGTCCCCACATGTTAGCGCTACCAAGTGCCGACGAACGCGCGCCGCCGACGGTGTCGGCGCTTCAGACGCTCGAGTTTTGGGGAGACGACGCCATATGTGTGGCAAGTACGGTTATTTTAAGTTAGGGTCCGAAATTTTCAGTGTAGAGAACCCCTTGACAATTGCTGAGATTGGTACGTCCCATAACGGAAGCATCCAAAAAGCACGCAATTTGATTGACGTGGCCGCCGAGGCGGGGGCTAAAGCGGTCAAATTCCAGATTGTCTATGCAGATGAGATCTTGCATCCGAACACAGGATATGTCGACTTGCCTACTGGTAAAATCCCGTTGTATGATCGTTTTAAGTCGTTGGAACTGCCGGTTAGTTTTTATAAAGAGTTGGCGGAGTATTCCCGTTCAAAAAAACTGTTGTTTTCAGCATCGCCATTCGGATTTCGCTCGGCCGAGGAGTTAGCTGCATTAAAGCCCGACTTTATTAAGATCGCGAGTCCCGAGTTGAATTATGTTCAATTGCTGAAGTACTGCGCTGGCTTCAACATTCCAATGATTCTGTCTAGCGGGGTATCTATTTTGAAAGATATTGAGAAAGCTGTCAGCTTCTTACGCAGCGAAAACAAAGATCTGCCTCTTGCTCTTCTGCACTGCATTACTTCATATCCAGCACCCGAAAAAGAATATAATGTTAGTGTAATCAAAAATTTATCTCGCATTTTTGGCATCGCGTGTGGAGTATCAGACCATTCACTGGACCCAATCTTGGTCCCGGCATTGACACTGGCAGCCGGAGGTTTCATTATCGAAAAGCATATTTGTTTATCACGCAAGGAGGAAGGATTGGATGATCCTGTTGCTTTAGAACCAGATATGTTTAAGAAAATGTGCTCGGCCTTGAACTCTTTCAGTAAGAAAACCTACGATGAAATTATTGAAGGTCTGACCAAGTTGAATTATTCAATTGAGCTTATTAACGAGGTTATCGGCTCCGGCGAAAAAAAACTTTCCCCAGCAGAAACGAAAAACTATGGTCGCACCAATCGTAGCATTCACTATCTGAACGACCTCAAGAAGGGTGATGTAATTACCGATAAAGACATTGCGGTGTTACGCACCGAGAAGATTCTGTCACCAGGCGAAGCTCCAGAAATGTTTGACTCTTTTATTGGGGCGGTGTTACAACGCGCGGTAAAAAGCGGCGAAGGGTTGCTTATGGAGGACTTTCTTACTCGTAAATGAATTCTAGAGGATCC |
| *A. baumannii ACICU neuB,* codon optimised for *E. coli* (5’-3’) | GGATCCCAATTGAACTAGTACCAGCCACAGGCCCGTGCCGGGATCGAAGGCGAAGCGGCAGCCGATCAGGCGGAACTGGGCGCGGGCCATGGTCTCGAACAGGGCCGTCAGGTCGCGGGCGGCGTCCAGGTCGTCGTGGTCCAGCACCAGGCAGCGGATCTGCAGCCAGCCGTGGTCGGGCAGCAGGTAGAAGGCGCCCTCGTCCTGATCCTCGCCCGAAACCTCCAGCCCCCGGTCGATGGCTTCGACGACATAGCCGGCCGCGCGGCAGGTGTCGGTGAGCGCGGCCAGCAGGGCGGCTTCCTGGTCAGGGGTCAGGTCGGTCATGGGCAAGAGGTCCAGGTCGTGGTTTGTCGGCGGCTTCTAGCATGGACCGCCCGCGCCCGTGAGGCCGAGGATTTCGCGCTGGTCAGACAACCTACTTGCCGTCCCCACATGTTAGCGCTACCAAGTGCCGACGAACGCGCGCCGCCGACGGTGTCGGCGCTTCAGACGCTCGAGTTTTGGGGAGACGACGCCATATGTCCAAAAAAATTACTATCAATAACCGCGACATCGGTACAGATTTTCCGCCGTATGTGATTGCGGAACTTTCGGCAAATCACAATGGGCGTTTAGAAACTGCACTTAAAATCATCGAGGAAGCTAAGCGTTGTGGTGCTGACGCTATCAAACTTCAGACATACTCCGCAGACACCATCACACTGAATAGTCAGAACGAAGAGTTTATGATTCGCGGTGGTCTGTGGGATGGGCAGTCCTTGTACGAGTTATACCAGAAAGCGCAAATGCCTTGGGAGTGGCACAAGCCGCTGTTTGATCACGCGCGTGCCTTGGACATTACTATCTTCAGTAGTCCATTTGATCGCACCGCCGTAGATCTGTTAGAAGATCTGAACGCGCCTGCATATAAAATTGCAAGTTTTGAAGCTGTCGACCTTCCATTGATTAAATATGTAGCGTCTACTGGAAAGCCCATGATTATTAGTACCGGAATGGCCGACAAGGAAGAGATTGGTGAGGCGATTCAAGCCGCCTACGACGGTGGATGCAAGGAGTTAGTGGTTCTTCACTGCGTGTCCGGATATCCAGCCCCGGCAGAGGATTACAACTTACTTACAATGGTAGACATGGCCAAATCATTTAATGTCCCTGTTGGGCTTAGCGATCATACCTTAGATAACACGACTGCAATTACGTCCGTAGCATTGGGCGCATGTCTTATTGAAAAGCACTTCACCCTTGATCGTAATGGCGGGGGCCCAGATGATAGTTTTAGCTTGGAGCCTAAAGAATTACAGCAGTTATGTCAGGGAGCTAAAACAGCCTGGCAGGCCTTGGGCCATGTAGACTATGGTCGCAAATCGTCAGAGCAGGGGAATGCGCAGTTCCGCCGTAGTTTATACTTCGTCAAGGACCTGAAGGCCGGTGACGTCATTGACGAGTCAAGTATCCGCTCTGTACGTCCCGGGTATGGGCTTCCTCCTAAGTTTTACGACGAGCTGTTAGGAAAAAAGGTGACGCAAGACGTGTCTGTAAATACCCCAGTCAAGCTGGAGATCATCAGTTGAATTCTAGACTCGAGAGCTCGCGGCCGCA |
| *A. baumannii LAC-4 neuB,* codon optimised for *E. coli* (5’-3’) | GGATCCCAATTGAACTAGTACCAGCCACAGGCCCGTGCCGGGATCGAAGGCGAAGCGGCAGCCGATCAGGCGGAACTGGGCGCGGGCCATGGTCTCGAACAGGGCCGTCAGGTCGCGGGCGGCGTCCAGGTCGTCGTGGTCCAGCACCAGGCAGCGGATCTGCAGCCAGCCGTGGTCGGGCAGCAGGTAGAAGGCGCCCTCGTCCTGATCCTCGCCCGAAACCTCCAGCCCCCGGTCGATGGCTTCGACGACATAGCCGGCCGCGCGGCAGGTGTCGGTGAGCGCGGCCAGCAGGGCGGCTTCCTGGTCAGGGGTCAGGTCGGTCATGGGCAAGAGGTCCAGGTCGTGGTTTGTCGGCGGCTTCTAGCATGGACCGCCCGCGCCCGTGAGGCCGAGGATTTCGCGCTGGTCAGACAACCTACTTGCCGTCCCCACATGTTAGCGCTACCAAGTGCCGACGAACGCGCGCCGCCGACGGTGTCGGCGCTTCAGACGCTCGAGTTTTGGGGAGACGACGCCATATGATTTTTAACCAAGACGAAATCTACGTTATCGCGGAAATCGGTGTGAATCATAACGGTTCTGTGGAGTTAGCGAAGGAGTTAATTCTTAAAGCGAAAGAGTGCGGTGCCAACGCCGTTAAGTTTCAAACGTTCAAAGCCGACTCTCTTCTGAGTGACCAGACTGAGATGGCAGCGTACCAGAAGGAAAATACCAGTTCCAGTCAATCCCAATTAGAGTTGGTTAAAAGCTCAGAGCTTACTTACGAGCAAACTGAGGAAATTCAGAAATTCTGTGATGAACATCAGATCACATTTATTTCGACACCGTTCGACTCCGACAGCCTTAAATTCCTGGTGAACGAAATCGATGTACCGTATCTGAAGGTTTCATCCGCGGACATCAGTAACTTGCCGTTCCTGTACGAGATCGCTTGTTCCAAGAAACATGTAATCCTTTCTACCGGCACTGCAAGCCTTGGGGATATTGAACAAGCTCTGTCCGTGTTTGCATTCGTTATTGACAAGGGAACAGAAGTGCGCCCTTCCCAACAACTTTTTCGCGAGGCATATAGTAAGATTTCCGTACGCAAACAGCTTAAAGAGCAAGTATCTATTTTACACTGTGTCACGCAATATCCCGCCTTATTTGAGGAGTCTAATCTGAAGGCAATTTCCACGCTTAAAAACGTTTTTGACTTAGCGACGGGATTCTCCGACCATACTTTGGATGAATATGCCGCAGTAATCGGAGTTTCTTTAGGTGCTCGCATTTTCGAGAAGCATATTACTCTGGATAAAACAATGGCGGGGCCTGACCATGCCGCGTCAATGGAGCCCGACGAGTTTAAGCACTATGTCGAAATTCTGCATAAAACGTATGCTGCGTTAGGCGATGGGATTAAATTCATGCTTGAACAGGAGTCCGATAACTATTACTTGGTGCGCCGCAGTATTGTGGCAAAGACTGACATCGCCGAGGGGGAACTGCTTACGTCGGACAATGTGACAACGAAGCGCGCAGGGCGCGTATGTCTTGAGCCCAATAAGTACTGGGACGTGGTGGGTACAAAAGCTAAACGCAGTTTTAAGAAAAATGATTTTATCGAAATTTGAATTCTAGACTCGAGAGCTCGCGGCCGCA |
| *A. baumannii LUH5551 neuB,* codon optimised for *E. coli* (5’-3’) | GGATCCCAATTGAACTAGTACCAGCCACAGGCCCGTGCCGGGATCGAAGGCGAAGCGGCAGCCGATCAGGCGGAACTGGGCGCGGGCCATGGTCTCGAACAGGGCCGTCAGGTCGCGGGCGGCGTCCAGGTCGTCGTGGTCCAGCACCAGGCAGCGGATCTGCAGCCAGCCGTGGTCGGGCAGCAGGTAGAAGGCGCCCTCGTCCTGATCCTCGCCCGAAACCTCCAGCCCCCGGTCGATGGCTTCGACGACATAGCCGGCCGCGCGGCAGGTGTCGGTGAGCGCGGCCAGCAGGGCGGCTTCCTGGTCAGGGGTCAGGTCGGTCATGGGCAAGAGGTCCAGGTCGTGGTTTGTCGGCGGCTTCTAGCATGGACCGCCCGCGCCCGTGAGGCCGAGGATTTCGCGCTGGTCAGACAACCTACTTGCCGTCCCCACATGTTAGCGCTACCAAGTGCCGACGAACGCGCGCCGCCGACGGTGTCGGCGCTTCAGACGCTCGAGTTTTGGGGAGACGACGCCATATGATTTTTAATCAGGATGAAATCTATGTAATTGCTGAAATCGGAGTCAATCATAACGGGTCAGTAGACCTTGCAAAGGAGTTAATTCTTAAAGCCAAAGAGTCAGGCGCGAATGCAGTCAAATTCCAGACTTTCAAGGCCGAAGAAGTTCTGTCGGATCAGGCGGATATGGCGGCCTATCAAAAGACTAACACTGGGAAAGACAGCAGTCAACTGGAGATGGTCAAGAAACTTGAACTTTCATTCGAGGACACAAAAATTCTGCGTGATTTTTCTGAAAACAATGGAATTACCTTTATCAGTACACCCTTCGACAATGACTCATTGGATTTCCTTTCCAACGACCTGGACGTACCTTTCTTAAAAGTATCCTCGGCTGACATTAGTAACTTACCTTTCTTATATAAGGTAGCGCTGACAGGAAAGCCTGTGATTTTATCTACAGGTACAGCTTCGCTTGGTGATATTGAGAAAGCACTTTCGGTGTTCACGTATGTCACAGAGTATGGCAAAACTGAGTTACCCACGGTAGAGAAACTTCGCTTGAGCTACGCCAAACAGACGAATCGCCAAGCATTAAAAAATAAAATTGTAATTCTTCACTGCGTAACCCAATATCCTGCCACGATTGAAGGTAGTAATTTGTTGGCGATTAGCACAATTCAGAATGCGTTCGGACTGAACGTAGGATATAGCGATCACACCCTTAATGAGTACACCGCTATCGCTGCTGTCGCCCTGGGAGCAACAGTTTTCGAGAAACACATTACTTTGGACAAAAAAATGGACGGGCCAGATCACGCTGCATCTATGGAAATTGATGACTTTACTCGTTATGTTTCGACGCTTCGTAAAATCAAACTGGGGCTGGGTGATGGTATCAAGTTCATGCAGGAACAGGAAAAAGACAATTTTTCGTTAGTACGCCGTAGTTTGGTTGCCCGTAAGGAGATCCAACAGGGCGAGATTTTCACGGAGCAAAATATTGTGGCTAAGCGTGCGGGGAAGGTCCAACTCCTGCCTGAACAACTGTGGGAGCTTCTGGGAACCGCAGCCACGAAGAACTACAAAGTTAACGAATTAATTGAGAAATGAATTCTAGACTCGAGAGCTCGCGGCCGCA |
| *L. pneumophila neuB,* codon optimised for *E. coli* (5’-3’) | ATGGGGTCGAACCGCAAAATTAACGGTATTAAACCGCGCGGAAGTTCGATGACGTGCTTTATCATCGCAGAAGCGGGCGTGAATCATAACGGAGATTTGCAACTTGCGAAGGAGCTGGTTTATGCTGCAAAGGAGTCGGGTGCAGATGCCGTAAAGTTCCAGACATTTAAAGCTGACACACTTGTCAACAAAACGGTAGAGAAGGCTGAATACCAGAAAAATAACGCTCCGGAGAGTTCCACTCAATACGAGATGCTTAAAGCGCTTGAGCTTTCTGAGGAAGACCACTACTTATTATCCGAGCTTGCAAATTCTTTGGGAATTGAATTTATGTCAACGGGGTTTGATGAACAGTCTATTGATTTTCTTATTTCACTTGGAGTGAAGCGTCTGAAGATCCCATCGGGGGAAATCACCAATGTTCCGTATCTTCAACACTGCGCTTCCAAAAAATTGCCACTTATTATTAGCACGGGTATGTGCGATTTACAGGAAGTGCGTGTAGCCATTGATACCGTAAAGCCGTATTATGGCAACTCTTTATCTGATTACTTAGTTCTGCTTCATTGCACAAGTAACTATCCCGCTTCTTACCAAGACGTAAATCTGAAAGCGATGCAAACATTGGCGGACGAGTTTCAATTACCGGTTGGATACAGTGACCACACTTTGGGAATCTTGGTCCCGACGTTGGCAGTGGGTATGGGGGCATGTGTCATTGAGAAACATTTTACCATGGATAAGTCTTTGCCAGGCCCGGATCACCTTGCTTCAATGGACCCAGAAGAGATGAAGAACCTTGTACAAAGTATCCGCGATGCAGAGACTGTCTTGGGTAGTGGCGAGAAGAAACCATCAGATAACGAGCTTCCGATTCGTGCTTTGGTCCGTCGTAGTATCACCTTGCGCCGCGATCTGGTAAAAGGTGCCCAGATTTCCAAAGAAGACCTGATTCTTCTGCGTCCAGGCACTGGAATTGCACCGAGCGAAATCTCCAACATCGTCGGTTCACGTTTAAGTATGAACTTGTCGGCAGGAACCACTTTGCTTTGGGAACATATTGAGGCCTGACTAGCTTAAGGAATTCTAGAGGATCC |
| *P. aeruginosa Y82 neuB,* codon optimised for *E. coli* (5’-3’) | CAATTGAGATCTGCTAGCCCAGCCACAGGCCCGTGCCGGGATCGAAGGCGAAGCGGCAGCCGATCAGGCGGAACTGGGCGCGGGCCATGGTCTCGAACAGGGCCGTCAGGTCGCGGGCGGCGTCCAGGTCGTCGTGGTCCAGCACCAGGCAGCGGATCTGCAGCCAGCCGTGGTCGGGCAGCAGGTAGAAGGCGCCCTCGTCCTGATCCTCGCCCGAAACCTCCAGCCCCCGGTCGATGGCTTCGACGACATAGCCGGCCGCGCGGCAGGTGTCGGTGAGCGCGGCCAGCAGGGCGGCTTCCTGGTCAGGGGTCAGGTCGGTCATGGGCAAGAGGTCCAGGTCGTGGTTTGTCGGCGGCTTCTAGCATGGACCGCCCGCGCCCGTGAGGCCGAGGATTTCGCGCTGGTCAGACAACCTACTTGCCGTCCCCACATGTTAGCGCTACCAAGTGCCGACGAACGCGCGCCGCCGACGGTGTCGGCGCTTCAGACGCTCGAGTTTTGGGGAGACGACGCCATATGAAAGCCCCGGATGAGCGCGCTGTATTCGTGATTGCAGAAGCGGGCGTTAATCACAACGGTGATCGCGATCTTGCATTTAAGTTGATTGACGTTGCGGCCCAAGCTGGAGTAGACGCCGTTAAATTTCAGACATTTAATGCGAAGCGCCTGGCCTCTCGTTCAGCCCCGAAAGCGAATTATCAGAAACACACGACTGATGTTACTGAATCACAGCTTGCCATGCTGAAAAAGCTGGAGTTGCCTAAAGAGTGGCATTTCGAACTTCAGGCACATGCTCACCATAACGGGATTGAGTTTATCAGTACGGCCTTCGACAGTGATTCCCTGGCGTTTCTGGCGGAAATGCAACTCCCATTTTTCAAGGTTCCATCTGGCGAGCTGACGAATGGTCCCCTGCTTTGGGCGTTCGCTAAGACTGGAAAACCTTTGATTTTATCAACTGGGATGGCCACCCTTTCTGAAGTGGAACAAGGATTGGCTATCGTGGCCCACGCGCTGAGCTGCGATAACGAGCCCAAAGATATGGATGAGGTTTGGCGTCTGTGGTCGAACCCTTCGGTTCGTATGCAACTCCAGGGTCACGTGTCGCTTCTTCATTGCACGTCACAATACCCAACGCCACCTGACGAGGTCAACTTACTGGCTATGGACACGTTGCGTTCCTTTGGCTTAGCTGTCGGTTATAGCGATCATACTGAAGGTGGTTTGGTCCCGATCGCTGCCGTAGCGCGCGGCGCTTGCATTATCGAGAAACACTTTACCCTGGACCGCTCAATGCCGGGGCCTGATCATAAAGCCTCCCTTGAGCCGGGGGAGCTTGCACAGATGGTGGCGCAGATTCGCATGTTGGAAGTGGCATTAGGTTCGCCTTATAAAGCGCCTCAGCCGTCAGAGTGGGATACACGCCAAGCAGCGCGTCAACAAGTCGTAGCTGCGCGCGACATCGAGGCAGGCATGATCATCACTCGTGATGACTTGACTACGGCACGCTCTGGTCATGGGCTGCCTCCAACATCCTTATGGGAGTTGGTGGGATCGACTAGCAAGCGCGCATTTTTGGCTGGAGAAACGCTGGAGAAATGAATTCTAGACTCGAGAGCTCGCGGCCGCA |
| *P. aeruginosa PASP475 neuB,* codon optimised for *E. coli* (5’-3’) | CAATTGAGATCTGCTAGCCCAGCCACAGGCCCGTGCCGGGATCGAAGGCGAAGCGGCAGCCGATCAGGCGGAACTGGGCGCGGGCCATGGTCTCGAACAGGGCCGTCAGGTCGCGGGCGGCGTCCAGGTCGTCGTGGTCCAGCACCAGGCAGCGGATCTGCAGCCAGCCGTGGTCGGGCAGCAGGTAGAAGGCGCCCTCGTCCTGATCCTCGCCCGAAACCTCCAGCCCCCGGTCGATGGCTTCGACGACATAGCCGGCCGCGCGGCAGGTGTCGGTGAGCGCGGCCAGCAGGGCGGCTTCCTGGTCAGGGGTCAGGTCGGTCATGGGCAAGAGGTCCAGGTCGTGGTTTGTCGGCGGCTTCTAGCATGGACCGCCCGCGCCCGTGAGGCCGAGGATTTCGCGCTGGTCAGACAACCTACTTGCCGTCCCCACATGTTAGCGCTACCAAGTGCCGACGAACGCGCGCCGCCGACGGTGTCGGCGCTTCAGACGCTCGAGTTTTGGGGAGACGACGCCATATGAATAAGGTGATCTCGATCGCTGGTCGTGCCATCGGGCCAGATTACCCCCCATATATTATTGCCGAAATGTCTGCTAATCATAATGGCTCCCTTGAAACGGCGTTTCGCATTATCGAAGCCGCCAAGAACTGTGGGGCGGATGCGTTAAAGATCCAGACATACCGCCCAGATACGATCACACTTAACAGCGACTTACCAGATTTCCGCATCTCCGGGGGGCTGTGGGGTGGCAAAACATTGTATGAATTATATGAATGGGCCCACACTCCATGGGAGTGGCATACTCCGATGTTCGAATATGCTCGTAAAATCGGGGTAACCATCTTCTCCTCACCGTTTGATCCAACTGCTGTCGACTTGCTGGAAGACTTAAATGCGCCGGCATATAAGATCGCATCGTTCGAGGCGATTGATTTGCCGTTGATCAAATACGTGGCTGCCACGGGAAAGCCTATGATCATTTCCACTGGCATGGCGGACTTGGAGGAAATCCAGGAGGCAGTCGACGCAGCAAAAGCGGGTGGGTGTAAGGAATTAGCTATCCTTCACTGCGTGTCAGGATACCCCGCGCCGGCCCAGGACTATAACTTACGCACCATTACCGACATGCAGAAACGCCATGGTTTGGTTATCGGATTATCGGACCACACTCTTGACAACACGACAGCTATCGCATCGGTGGCCCTGGGGGCGAGTATTATTGAGAAACATTTCACCCTGAACCGCAACGGTGGAGGTCCCGATGACAGTTTTTCATTGGAACCAGCGGAGCTGAGTGCGCTTTGTCGCGACAGCAAGACTGCGTGGGCATCTCTTGGGATCGTAGATTACGGTCGTAAATCGAGCGAAGCAGGTAATGTGAAATTTCGTCGCTCCCTGTACGCCATCCGTGACATTGAGGCTGGAGAGCCGCTTACGGCAGAAAACATTCGCAGTGTGCGCCCCGGATATGGGCTGTCGCCCAAGCACTACAAAGAGGTCCTTGGCAAAACTGCGGTAGTTAAAATCCCTGCGCACACCCCTATTCACTGGGAAAAGCTCTCCGATGACTGAATTCTAGACTCGAGAGCTCGCGGCCGCA |
| *S. japonicum neuB,* codon optimised for *E. coli* (5’-3’) | GGATCCCAATTGAACTAGTACCAGCCACAGGCCCGTGCCGGGATCGAAGGCGAAGCGGCAGCCGATCAGGCGGAACTGGGCGCGGGCCATGGTCTCGAACAGGGCCGTCAGGTCGCGGGCGGCGTCCAGGTCGTCGTGGTCCAGCACCAGGCAGCGGATCTGCAGCCAGCCGTGGTCGGGCAGCAGGTAGAAGGCGCCCTCGTCCTGATCCTCGCCCGAAACCTCCAGCCCCCGGTCGATGGCTTCGACGACATAGCCGGCCGCGCGGCAGGTGTCGGTGAGCGCGGCCAGCAGGGCGGCTTCCTGGTCAGGGGTCAGGTCGGTCATGGGCAAGAGGTCCAGGTCGTGGTTTGTCGGCGGCTTCTAGCATGGACCGCCCGCGCCCGTGAGGCCGAGGATTTCGCGCTGGTCAGACAACCTACTTGCCGTCCCCACATGTTAGCGCTACCAAGTGCCGACGAACGCGCGCCGCCGACGGTGTCGGCGCTTCAGACGCTCGAGTTTTGGGGAGACGACGCCATATGACCGACTGGCTGCCTAAGAAGGCTATCGTAATCGCCGAACTGTCGGCTAATCACAATGGTTCTCTTCAACGCGCGTGTGATACAGTTCGCGCCATGGCGGACTCGGGTGCCGACATTGTTAAAGTGCAAACATACCGCCCCGAATCTTTGTGTCTGGACGTCGACAACGACTACTTCGGAAAACGTAAAGCAGGATTATGGAAAGGGCGTACTTTATGGAGCCTGTACCAAGAGGCAGCACTTCCGTATGAGTGGCATCAACCTCTGCAAGCATTAACTCACGAGTTAGGAATGGCGTTTTTTAGTTCGCCGTTTGACTTGGAGGGAGTTGATTTCCTGGAAACATTAAATGTACCTTATTACAAGATTGCGTCGTTTGAGATCAACCACATCCCCCTTATTGACAAAGCTGCGCGTACTGGAAAACCCTTGATCATCTCGACGGGAGTCGCTGACGACAAGGACATTGATCTGGCTCTTAAGACATGCTTTGAGGCAGGCAATACGAATGTTGCACTGTTGAAGTGCACCTCACAATACCCCGCGTCGTTAAATGACGCTAACTTACACATGATCACCGACATGCAAAAGAAGTTTAACGTCCCTATCGGCTTAAGCGACCATACAATGGGCTCCTTGGTTCCGATGACTGCGGTTAGTTTGGGGGCCCGTATCATCGAGAAGCACTTCACGCTGGATCGTCACGATGGTGGCCCGGACAGCGCATTTTCTATGGAACCGGACGAGTTTCGCGACATGGTCAACCAAATTCGCAACGTCGAGAGCTGCCTGGGTAAGGTCAACTATAGTGTTTCAGAACAAGACAAGTTGCGTCGCCGTAGTATCTTCTACTGTCGTGACCGTAGCGCCGGTGAATTGATTGGCGAGGAAGACATCAAGGTTCTTCGCAATGGATCTGGACTGCATCCTAAGCACATGAAGAGCTTGATTGGAGCCACTTTAACACGCAGCGTGAACGTAGGAGAACCAGTCTTGGCGGAGGATTTGGACAAAAAAGACTTTTTGCAACAATGAATTCTAGACTCGAGAGCTCGCGGCCGCA |
| *B. subtilis PY79 neuB,* codon optimised for *E. coli* (5’-3’) | AGATCTGCTAGCCCAGCCACAGGCCCGTGCCGGGATCGAAGGCGAAGCGGCAGCCGATCAGGCGGAACTGGGCGCGGGCCATGGTCTCGAACAGGGCCGTCAGGTCGCGGGCGGCGTCCAGGTCGTCGTGGTCCAGCACCAGGCAGCGGATCTGCAGCCAGCCGTGGTCGGGCAGCAGGTAGAAGGCGCCCTCGTCCTGATCCTCGCCCGAAACCTCCAGCCCCCGGTCGATGGCTTCGACGACATAGCCGGCCGCGCGGCAGGTGTCGGTGAGCGCGGCCAGCAGGGCGGCTTCCTGGTCAGGGGTCAGGTCGGTCATGGGCAAGAGGTCCAGGTCGTGGTTTGTCGGCGGCTTCTAGCATGGACCGCCCGCGCCCGTGAGGCCGAGGATTTCGCGCTGGTCAGACAACCTACTTGCCGTCCCCACATGTTAGCGCTACCAAGTGCCGACGAACGCGCGCCGCCGACGGTGTCGGCGCTTCAGACGCTCGAGTTTTGGGGAGACGACGCCATATGGCCGCTTTCCAAATTGCCAACAAGACCGTTGGGAAGGATGCGCCAGTGTTCATTATCGCCGAAGCGGGCATCAATCACGACGGGAAGTTAGACCAAGCATTTGCACTGATCGACGCGGCAGCAGAGGCAGGAGCCGACGCAGTGAAATTTCAAATGTTTCAAGCAGATCGTATGTACCAAAAAGATCCTGGACTTTATAAAACAGCAGCAGGCAAAGATGTCTCGATCTTCAGTCTGGTACAAAGTATGGAAATGCCTGCGGAATGGATCTTACCATTGCTGGACTACTGCCGTGAGAAACAAGTCATCTTCTTATCGACAGTCTGCGACGAAGGCAGCGCTGACCTTTTGCAGTCTACTAGCCCGTCAGCGTTTAAAATCGCGTCCTATGAGATCAACCATTTACCTCTGTTGAAGTACGTTGCACGTTTGAACCGCCCAATGATTTTCTCGACCGCGGGGGCGGAGATTAGTGACGTACACGAGGCTTGGCGCACTATTCGTGCAGAAGGGAATAACCAGATTGCCATTATGCACTGCGTAGCTAAGTACCCCGCCCCACCGGAATACTCAAATCTGAGCGTAATTCCCATGTTAGCCGCGGCATTTCCCGAAGCCGTGATCGGATTCTCGGATCACAGTGAGCACCCGACAGAAGCTCCCTGTGCAGCAGTGCGCTTGGGCGCCAAGTTGATCGAAAAGCACTTTACAATCGACAAGAACCTTCCTGGAGCAGATCATAGTTTTGCGTTAAATCCTGATGAGCTGAAAGAGATGGTGGACGGGGTCCGTAAGACTGAGGCTGAGCTCAAGCAGGGCATCACTAAACCTGTGTCTGAAAAACTGCTTGGCAGTAGTTATAAAACCACTACCGCCATTGAGGGGGAAATTCGTAATTTTGCCTATCGCGGTATTTTTACAACAGCTCCTATCCAAAAGGGAGAGGCATTTAGTGAAGATAACATCGCTGTACTTCGTCCCGGACAAAAACCCCAAGGATTGCACCCGCGCTTCTTCGAACTTTTAACGAGTGGGGTGCGTGCAGTACGCGATATCCCAGCGGACACTGGCATTGTCTGGGATGATATTTTGTTAAAAGATAGCCCCTTCCATGAGTGAAACTAGTACTTAAGAAGGAGAGAATTCATGTCTAGAAAAGAGCTCGCGGCCGCGGATCC |
| *K. sibirica neuB,* codon optimised for *E. coli* (5’-3’) | CAATTGAGATCTGCTAGCCCAGCCACAGGCCCGTGCCGGGATCGAAGGCGAAGCGGCAGCCGATCAGGCGGAACTGGGCGCGGGCCATGGTCTCGAACAGGGCCGTCAGGTCGCGGGCGGCGTCCAGGTCGTCGTGGTCCAGCACCAGGCAGCGGATCTGCAGCCAGCCGTGGTCGGGCAGCAGGTAGAAGGCGCCCTCGTCCTGATCCTCGCCCGAAACCTCCAGCCCCCGGTCGATGGCTTCGACGACATAGCCGGCCGCGCGGCAGGTGTCGGTGAGCGCGGCCAGCAGGGCGGCTTCCTGGTCAGGGGTCAGGTCGGTCATGGGCAAGAGGTCCAGGTCGTGGTTTGTCGGCGGCTTCTAGCATGGACCGCCCGCGCCCGTGAGGCCGAGGATTTCGCGCTGGTCAGACAACCTACTTGCCGTCCCCACATGTTAGCGCTACCAAGTGCCGACGAACGCGCGCCGCCGACGGTGTCGGCGCTTCAGACGCTCGAGTTTTGGGGAGACGACGCCATATGTTAGAGATCAACATTCGTGTTGAGAATCAGCTTATCAGTAGTAACTCCCGCCCTTTCATTATCGCTGAGATGTCCGGCAACCACAACCAATCGTTAGAGCGTGCTCTGCAGATTGTTGAAGCGGCAGCGAAGAGTGGCGCTCACGCACTGAAAATCCAGACTTATACTGCAGACACGATTACTTTGAACTCCGAGCAGGCGGATTTCATGATCGAGGATCAGGAGAGCTTGTGGAAGAATCGCAAGCTGTACGACCTTTACAACGAAGCGTACACTCCGTGGGAGTGGCACAAACCCATTTTCGACCGTGCCAAGGAATTAGGCATGATTGCGTTTTCCACGCCCTTCGACAATACCGCGGTTGATTTCTTAGAGACCTTGAATGTCCCACTGTATAAAATCGCTAGTTTCGAGAACACTGATCTTCCTTTAATTAAACGTGTAGCAGAAACAGGCAAACCAATGATCGTGAGTACTGGTATGGCAAGTGTAGCTGAGCTGGATGAACTGGTAAAGACGGCCAAGGAGGCAGGCTGCAAGGATCTTATTTTATTGAAGTGCACTTCTACCTATCCTGCGACACCCGAGAACACTAATATTATCACGATTCCTCACATGAAGGACCTTTTTGATGTGCAGGTGGGCCTTTCAGATCATACAATGGGTACTGGTGTATCAGTTGCCGCCGTGGCCCTGGGTGCTACTGTAATCGAGAAACATTTTACGCTTTCACGTGCCGAAGGCGGAGTGGATGCGGCGTTCTCCTTGGAGCCCGCAGAGATGACTGCCTTGGTGGAAGAGACGGAACGCGCCTGGCAGGCCATCGGCAAGGTGACCTATGGTCCTACTGAAAAGGAAAAGGCATCCCTGAAGTTCCGCCGCTCAATCTACGTGTCGAACACAATCGTAGAAGGAGACCTTTTTACAGAGGAAAATATTAAAATTGTCCGTCCGGGACTTGGACTTGAGCCTAAGTACTATCCCATCTTACTGGGTAAGAAAGCAAAGAAATCATACCAGTTTGGAGAGCCGATTAAATTCGATGACCTTATCTGAATTCTAGACTCGAGAGCTCGCGGCCGCA |
| *M. humiferrea neuB,* codon optimised for *E. coli* (5’-3’) | AGATCTGCTAGCCCAGCCACAGGCCCGTGCCGGGATCGAAGGCGAAGCGGCAGCCGATCAGGCGGAACTGGGCGCGGGCCATGGTCTCGAACAGGGCCGTCAGGTCGCGGGCGGCGTCCAGGTCGTCGTGGTCCAGCACCAGGCAGCGGATCTGCAGCCAGCCGTGGTCGGGCAGCAGGTAGAAGGCGCCCTCGTCCTGATCCTCGCCCGAAACCTCCAGCCCCCGGTCGATGGCTTCGACGACATAGCCGGCCGCGCGGCAGGTGTCGGTGAGCGCGGCCAGCAGGGCGGCTTCCTGGTCAGGGGTCAGGTCGGTCATGGGCAAGAGGTCCAGGTCGTGGTTTGTCGGCGGCTTCTAGCATGGACCGCCCGCGCCCGTGAGGCCGAGGATTTCGCGCTGGTCAGACAACCTACTTGCCGTCCCCACATGTTAGCGCTACCAAGTGCCGACGAACGCGCGCCGCCGACGGTGTCGGCGCTTCAGACGCTCGAGTTTTGGGGAGACGACGCCATATGTTCAATCGCGTCTTCATTATTGCAGAAGCGGGCGTAAACCATAATGGCGATTTAGAATTAGCGAAAAAACTTGTCGATGCTGCAGTCGAGGCAGGCGCCGACGCTGTCAAATTCCAGACCTTCAAGGCGGAGGAGGTAGTGACCCCAAATGCTGAACGTGCGCAATATCAAAAGGATAACATGCCAGAGAAAGACGAGTCACAGTTAGAAATGATTAAACGCTTGGAACTGAGTTATGCTCAGTTCCGTGAACTGTATGATTATTGCCGTCAGAAGGGGATCATCTTTCTTTCCTCCCCCTTTGACCAAGAGTCAATCGATTTCTTGGCCGAACTTGGTGTGCCTTACTTCAAGATTCCTAGTGGGGAAATCACCAATTATCCGTTTTTACGCCGCATCGCCGAAAAAAAACGCCCAGTCATTCTTTCCACAGGTATGGCGACATTGGGGGAGGTGGAGGGAGCGTTACAAGTATTACGTGAAGCTGGCGCGAAAGACATTACCTTGCTTCACTGCACGACTAGCTACCCGGCGTTACCCGAGGAGGTCAATCTTAAAGCAATGCTGACCATGAAACAAGCATTCGCCCTTCCAGTAGGATACAGCGATCACACCATGGGAATCGCAGTCCCTATTGCCGCAGCCGCCCTGGGTGCCGAGGTGATTGAGAAGCACTTAACGTTGGATCGTAATTTGCCGGGGCCGGACCACCGTGCATCATTGGAGCCTGGAGAGTTCAAGGAAATGGTGGCGGCCATTCGCCAGGTCGAAAAATCCTTGGGCGATGGAATGAAGCGTCCGGCACCTGGGGAGTTGGCAGTGATGCCGGCTGCGCGTCGCAGTCTTGTGGCGGCCCGCGATATCGCCGCAGGTGAAATCATTACGGAATCATGCCTTACCGCCCGCCGCCCAGGTACTGGGATTCCGCCGAATTTCTGGGACGTAGTGGTGGGTCGTCAGGCCCGCCGCGACATTGCAGCAGGATCTATCCTTAGTTGGGACATGATTTGAAACTAGTACTTAAGAAGGAGAGAATTCATGTCTAGAAAAGAGCTCGCGGCCGCGGATCC |
| *Mycobacterium sp. KBS0706 neuB,* codon optimised for *E. coli* (5’-3’) | CAATTGAGATCTGCTAGCCCAGCCACAGGCCCGTGCCGGGATCGAAGGCGAAGCGGCAGCCGATCAGGCGGAACTGGGCGCGGGCCATGGTCTCGAACAGGGCCGTCAGGTCGCGGGCGGCGTCCAGGTCGTCGTGGTCCAGCACCAGGCAGCGGATCTGCAGCCAGCCGTGGTCGGGCAGCAGGTAGAAGGCGCCCTCGTCCTGATCCTCGCCCGAAACCTCCAGCCCCCGGTCGATGGCTTCGACGACATAGCCGGCCGCGCGGCAGGTGTCGGTGAGCGCGGCCAGCAGGGCGGCTTCCTGGTCAGGGGTCAGGTCGGTCATGGGCAAGAGGTCCAGGTCGTGGTTTGTCGGCGGCTTCTAGCATGGACCGCCCGCGCCCGTGAGGCCGAGGATTTCGCGCTGGTCAGACAACCTACTTGCCGTCCCCACATGTTAGCGCTACCAAGTGCCGACGAACGCGCGCCGCCGACGGTGTCGGCGCTTCAGACGCTCGAGTTTTGGGGAGACGACGCCATATGCAAGAAATTAGCATCGCCGGTCGCCGTATTGGTCCCGACCATCCGCCGTACGTTATTTGTGAACTCTCGGGTAACCATAACGGCGAGCTTGCGCGCGCACTGCGGATGATTGAAGCCGCGAAAGCGACCGGTGCGGATGCCGTGAAGCTCCAGACGTATACGGCCGACACGATCACCCTCGACCATGATGGTGACCAATTTCGTATTAAAGGTGGTCTCTGGGATGGCCAACGGCTGTATGACCTTTATCAAGAAGCGTCCACCCCGTGGGATTGGCACGAAGCGCTTTTTGCGAAGGGTCGCGAGCTTGGCATTACGGTGTTTTCCAGCCCGTTCGATAAAACCGCCGTTGACCTCCTTGAAGGTCTTGGTGCCCCGGCCTACAAAGTTGCATCGTTCGAAGTCGTCGACCTGCCCCTGGTCGAGTACATCGCCTCCAAAAAAAAGCCGATGATCATCAGCACGGGCCTGGCAAACCTTGGCGAAATCCAAGAAGTTATTACCACCGCCCGTAAATCCGGCGCGGATGGCCTCGTGGTCCTGCATTGCATCTCCGCATATCCCGCACCCATGGAGGATGCGAATCTGCGTACGATTCCGAATCTCGCGGAGACGTTCGGCGTTATTTCCGGCCTTAGCGATCACACCATGGGTACGGCCGCAGCCGTGGCGGCGGTCGCGCTGGGTGGCTCCGTGATTGAAAAGCACTTTACCCTCGCGCGCGCCGATGGTGGTCCCGACTCGGCATTCTCCCTCGAACCCGCCGAATTTACGCGGCTCGTTGAGGACTGTAAAGGTGCCTGGGCGGCACTTGGTCGTATCCGTTACGACCTTAAGGGCTCGGAAGCCGGCAACATCGTGTATCGGCGGTCGCTTTACGTTACGCGTGATGTCAAAGCGGGTGAGACGCTGAGCGAGGCGAACGTTCGCAGCATTCGTCCGGGCTATGGCCTTGCCCCCAAGCACCTCCCGGAAGTTCTGGGCCGTCGTGCAGCCCGCGATCTCGCACGCGGCGAACCGTTCGCCTGGTCGATGGTCGAGTGAATTCTAGACTCGAGAGCTCGCGGCCGCA |
| *Halorubrum sp. PV6 neuB,* codon optimised for *E. coli* (5’-3’) | GGATCCCAATTGAACTAGTACCAGCCACAGGCCCGTGCCGGGATCGAAGGCGAAGCGGCAGCCGATCAGGCGGAACTGGGCGCGGGCCATGGTCTCGAACAGGGCCGTCAGGTCGCGGGCGGCGTCCAGGTCGTCGTGGTCCAGCACCAGGCAGCGGATCTGCAGCCAGCCGTGGTCGGGCAGCAGGTAGAAGGCGCCCTCGTCCTGATCCTCGCCCGAAACCTCCAGCCCCCGGTCGATGGCTTCGACGACATAGCCGGCCGCGCGGCAGGTGTCGGTGAGCGCGGCCAGCAGGGCGGCTTCCTGGTCAGGGGTCAGGTCGGTCATGGGCAAGAGGTCCAGGTCGTGGTTTGTCGGCGGCTTCTAGCATGGACCGCCCGCGCCCGTGAGGCCGAGGATTTCGCGCTGGTCAGACAACCTACTTGCCGTCCCCACATGTTAGCGCTACCAAGTGCCGACGAACGCGCGCCGCCGACGGTGTCGGCGCTTCAGACGCTCGAGTTTTGGGGAGACGACGCCATATGGAGATCGACGGCACGCGTATCGGCCCTGATAGCCCTCCTTTTTTTATTGCAGAGGCGGGGGTCAATCACAACGGAGAGCTGAAGAAAGCAAAAGAGCTGATTGATGTAGCCGCTGATGCCGGTGCCGACGCGGTGAAGTTTCAGACATTCACGGCAGATCGTCTTGTGACGCCTCATGCCGACAAAGCTGACTATCAGACGGAGACCACCGGCGAGGGCGGGCAATATGAGATGTTAAAGCAATATGAGTTAGATCGTGAAAGTCACCGCTTACTTTTAGACTACTGCTCAAAAAAAGACATTACGTTTTTGTCCACTCCTTTCGACCGTGAAAGCGCGGATATGTTGAAACAATTAGGTGTGGGGGCGATTAAGCTGGGTTCGGGTGAACTTACTAACATCCCATTGATCGAACATGTCGCCAAGTTCGATTTGCCTCTTATTGTTTCTACCGGGATGGGGACTTTGGAAGAAGTAGAGCAGGCGTATGAGGCAATCCAATCAGTAGACGTAGGGGCGGACATCGTTTTTCTGCATTGTACCTCGACATATCCGTGCTCTCCTGAAGACGTCAATTTGCGTGCGATGGAAACAATCAAGGAGAAACTGGATGTGTCGGTCGGGTATTCGGATCACACTGTTTTGCCGGAAACACCTGCCTTTGCCGTTGCTGCCGGGGCGAGTATCTTAGAAAAACACTTCACACTTGATAGTACGCTTCCAGGGCCTGACCACGAGGCGTCGCTTGAGCCTGAAGGACTGAATCATGCTGTTGACCTGGTGCACACCGCAGCGCAGATTCGTGGTAATCCGCGCAAACAACCCACCGAATCTGAACAAGAGAATATCCGTACCATCCGTAAGTCGCTTTACGCAGCATCGGACTTGGACAGTGGAGATTCATTAGAAGAGTCTGACATTGCTATTTTACGCCCTGAAGATGGTCTGAGCCCTCAATGGTATGACAGTGTTATTGGTATGAAAATCACGCAGGATATCCCCCAGGGTAAGCCCATCACCAAGTCTGATCTTGAGATCGAGGAAGGGGGATTATGACTAGCTTAAGGAATTCTAGAGGATCC |
| *M. smithii neuB,* codon optimised for *E. coli* (5’-3’) | AGATCTGCTAGCCCAGCCACAGGCCCGTGCCGGGATCGAAGGCGAAGCGGCAGCCGATCAGGCGGAACTGGGCGCGGGCCATGGTCTCGAACAGGGCCGTCAGGTCGCTGGCGGCGTCCAGGTCGTCGTGGTCCAGCACCAGGCAGCGGATCTGCAGCCAGCCGTGGTCGGGCAGCAGGTAGAAGGCGCCCTCGTCCTGATCCTCGCCCGAAACCTCCAGCCCCCGGTCGATGGCTTCGACGACATAGCCGGCCGCGCGGCAGGTGTCGGTGAGCGCGGCCAGCAGGGCGGCTTCCTGGTCAGGGGTCAGGTCGGTCATGGGCAAGAGGTCCAGGTCGTGGTTTGTCGGCGGCTTCTAGCATGGACCGCCCGCGCCCGTGAGGCCGAGGATTTCGCGCTGGTCAGACAACCTACTTGCCGTCCCCACATGTTAGCGCTACCAAGTGCCGACGAACGCGCGCCGCCGACGGTGTCGGCGCTTCAGACGCTCGAGTTTTGGGGAGACGACGCCATATGGAGTTCAAAATTGAAGATCGTCTGATCGGTGATGGCCACCCTGCATTTATCATCGCAGAATTAAGTGCCAATCACATGAATGATTATGATATCGCTGTAAAAACAATCGAGGCCATGGCAAAAAGTGGGGCCGATGCGGTGAAGTTTCAGACGTACACACCTGACACAATTACCTTGGACTGTGACAATGAATACTTCCAGATCAAGCAAGGCACGATCTGGGATGGGCAAGTGTTATATAACCTTTACGAAGATGCATTTATGCCCTGGGATTGGCAACCTAAACTTAAAAAGGTCGCTGAGGACTTAGGTTTAATCGTGTTCTCATCTCCGTTTGACGAAACTTCTGTAGACTTTCTGGAAGACATGGACATGGGCGCATACAAGATTGCATCGTTTGAAATTACGGATATTCCACTGATTGAATATGTAGCCAGCAAAAACAAGCCTGTAATCATTTCAACTGGCATCGCTTCCAAAGAAGACATTGATCTGGCTATCAAAACTTGCAAAGATGCCGGGAATGATAAGATCGCAGTTCTTAAATGTACGTCAAGCTACCCGGCTCCTTTGGAAGAGATTAATCTCAAGACAATCCCCGATCTGAAGGAAAATTTCAAGACTGTCGTGGGACTGTCGGATCATACCTTGGGATCTGACGTGGCAGTGGCCAGTGTGGCTATGGGCGTGAAAATCATCGAGAAACACTTTATTTTAGACCGCACGATGGAGGGACCAGATAGCGACTTCAGCATGGAACCCGACGAGTTCAAACAAATGGTCGATTCAATCCGTAATGTGGAAAAAGCTCTTGGCAAAGTATCATACGAATTATCTGATAAGATGAAGGCAAATCGCGAGTTTTCCCGTTCTCTGTTCGCAGTGAAAGACATTAAAAAGGGTGAGCTTATTACTAAGGACAACGTTAAATCAATCCGCCCCGGCTTCGGTCTTCACCCAAAATATTTATCTGAAATCATCGGATGTCGCGCCAGCGAAGACATCGACCGTGGAACACCGTTTAAGTTGGAGTTTGTGAATAAGTGAAAACTAGTACTTAAGAAGGAGAGAATTCATGTCTAGAAAAGAGCTCGCGGCCGCGGATCC |
| *flmG Cc-Bs* chimera*,* codon optimised for *E. coli* (5’-3’) | GGATCCCAATTGAACTAGTACCAGCCACAGGCCCGTGCCGGGATCGAAGGCGAAGCGGCAGCCGATCAGGCGGAACTGGGCGCGGGCCATGGTCTCGAACAGGGCCGTCAGGTCGCGGGCGGCGTCCAGGTCGTCGTGGTCCAGCACCAGGCAGCGGATCTGCAGCCAGCCGTGGTCGGGCAGCAGGTAGAAGGCGCCCTCGTCCTGATCCTCGCCCGAAACCTCCAGCCCCCGGTCGATGGCTTCGACGACATAGCCGGCCGCGCGGCAGGTGTCGGTGAGCGCGGCCAGCAGGGCGGCTTCCTGGTCAGGGGTCAGGTCGGTCATGGGCAAGAGGTCCAGGTCGTGGTTTGTCGGCGGCTTCTAGCATGGACCGCCCGCGCCCGTGAGGCCGAGGATTTCGCGCTGGTCAGACAACCTACTTGCCGTCCCCACATGTTAGCGCTACCAAGTGCCGACGAACGCGCGCCGCCGACGGTGTCGGCGCTTCAGACGCTCGAGTTTTGGGGAGACGACGCCATATGTCGCGCAAGAGCGCTTTAGAATCGAGCGCTTCTGTCTTGGCCCAGGCAGATGTTGGAGCCAGCGGGATTCATCCGAGCGTAATTGCGGATGCAATGGGGGATTCGGCGAGTGCCGAGGCATTAGAGCGTTTGAACCGCGCGGCGCAGGACACAAAAAACGTCGACAACGCAAAGCATTTGGCACGTGCGATCCAGGCTGTCCAGCTGCAAGACTACGCTAAAGCTGACAAGCTTGCGCTTAAGCTTCTTGAAAAGGATGAACGCTTAGGGTTGGCTTGGCACATCTTAGCCATTGCCCGCGAAAAGACAGGTGACTTCGCCTCATCTCTGCGCGCTTACGAAGCTGCTCTGGCACTTCTTCCTGATCATGGCCCTGTAGCAGGGGATTTAGGCCGTTTAGCATTCCGCATGAATATGCCCGAACTGGCAGCGAAATTCTTTGCGCATTATCGCTTGGCCCGCCCGGACGACGTCGAGGGAGCCAATAACTTAGCTTGTGCCCTTCGCGAGCTGAACCGTGAATCTGAAGCTATCGAGGTATTGAAGGCAGCGCTTGGAGCGAATCCAGAGGCAGCGGTCCTGTGGAACACACTGGGCACTGTGTTATGTAATATTGGCGACGCCGCAGGGAGCATCGTATTTTTTGACGAGAGCTTGCGCTTGGCACCTGACTTCTCCAAGGCTTACCATAACCGTGCTTTTGCACGCCTTGATCTGGGGGAGATCGAAGCAGCACTGGCGGATTGTGAGGCAGCTATGCGTTCCCCAGGCAGCCCAGAAGATCTTGCAATGATGCAGTTCGCACGCGCTACAATCCTTCTTGCATTGGGTCGTGTAGGGGAGGGTTGGGAGGCCTATGAAAGCCGCTTTAGCCCCGCACTTAGTGACGCCATGCATGTTGCCGTCGATGCTCCCCGTTGGGACCCGGCGACGCAGGATATCGCAGGGAAACGTTTGCTGGTAGTCGGTGAACAAGGCATTGCGGATGAGATGGTTTTCGGAGGATGCTTACCCGATGTAATTGAGGCGGTGGGACCAACAGGCAAAGTCTTCATCGCCGTGGAAGCTCGCTTGGTGGACTTGTACCAGCGCTCTTTCCCCACGGCGGTTGTCGGTGCCCACCGTGCGGTCCGCCTTGAAGGCCGCTTGACACGCTACTGCCCTTTTATGGAGCAAATCGGTGAAACTGAGGGAAAAGCAGACGCCTGGACACCCATGGCCAGTTTATGCGCCGTTTACCGTCGTGACCTGTCAGCGTTTCCCGATCGCGATGGGTACTTGATGCCAGATCCAGTAAAGGTGGCCCGTTGGAAGTCCGAGTTAGAGGCGCTTGGTCCTGGGCTTAAGGTTGGTCTGCATTGGAAGTCACTTGTACTGACAGGGGTGCGTGCCCGTTATTTCTCTAGTTTCGAGCGCTGGAAGCCAGTGCTGACTGCTCCTGGCTGTATTATGGTCAACTTACAGTGCGGGGACGTGACTGAAGACCTGGCAGCTGCGGAGGCTGCGGGTGTACGTATCTGGACACCACCAATGGATCTTAAGGACGATCTTGACGACCTGGCGGCCTTGTCTTGTGCACTTGATCTTGTTGTCGGTCCTGGGATCGCTGGTACGAACATCGCTGCCGCAGCCGGAGCACGCACATGGTTGATTCACGCCCCCGACGATTGGCATCTGCTGGCGACCGATCGCTATCCCTTCTATCCGCGCGTACGTACTTTCGCCACTGGAGGCTTTGATGGTTGGCCTCGCGCGATCGGCGCCGTGCGCGCGGCTCTGGAAAATGAGGTTGGAAACGCAGCATGAATTCTAGACTCGAGA |
| **Synthetic DNA** (from Genewiz, Leipzig, Germany) | |
| *B. subvibrioides Bresu_3266, Bresu_0765, Bresu_0506, Bresu_3264, Bresu_0507, Bresu_3265,* codon optimized for *E. coli* (5’-3’) | AGGATCCGGATGTGAGCGGATAACAATTACGAGCTTCATGCACAGTGAAATCATGAAAAATTTATTGGCTTTGTGAGCGGATAACAATTATAATATGTGGAAAGAAGGAGATACCATATGGAAGGGGACCGTACGACTGCGAGCTTGGCTGGGCAACGCGTGCTGGTCACCGGGGCTGGTGGGTTCATTGGCAGTCGTCTGTGCGAGCGTTTAGTAGCAGATGGGGCTGAAGTCCGTGCCTTGGTACGTTATACGTCGGATGGTGATGCTGGATGGTTGGATCGCTCCCCTATCCGCAAGGACATCGCCGTAGTTCGCGGCGATTTAGCCGACCGTGATAGTGTATTTGCAGCGGTACGCGACCGTGATGTAGTGTTCCATCTGGGCGCGTTAATCGCCATCCCATATTCCTATGAGGCGCCGGAATCCTACGTTCGTACGAATATCCTGGGCACCCTTAATGTGCTTCAGGCAGTACGTGAGCTTAGCGTGGGCCGCCTGATTCACACAAGTACTTCAGAGGTGTATGGGTCGGCCCAGACTGTTCCAATGACAGAAGCACACCCCCTTGTCGGGCAGTCGCCTTATAGCGCATCCAAAATCGGGGCGGACAAGTTGGCAGAATCTTACCATCGCAGCTTCGGGACGCCTGTGGTTACATTACGTCCTTTCAACACTTTTGGCCCGCGTCAGAGCGCCCGCGCCGTCATTCCCTCGATTACTATGCAACTGCTTGCCGGGCGCACCATTCGTATGGGTGATACACGCCCAACCCGCGACTTTGTTTTTGTAGACGATACAGTGGACGCCTTTGTCCGTGCGGCGACAGCATCGGGCATCGAGGGCCTTACCATTCATTTCGGCGGCGGGCGTGAAATTGCCATTGGCGATCTTCCAGCTTTAATCGGGGCTGCTGCGGGCCTGCCCGTAAGTGTTGAGATCGACCCCCAACGTTTGCGCCCTGCTGCTAGTGAAGTAGAACGCCTGATTGCAGACGCATCATTGGCCCGCCAGCGCCTTGGCTGGCAGCCGCGCGTCTCGGTAGAGGAAGGATTAGCCCGCGTCGTTGCATTTATTCGTGACCACCCGGGCTTATACCGTCCAGAAGAGTACGCCGTCTGAAGATCTCCCCCGGGAAGAAGGAGATATACCATGCGCCGCATTCCTCTTTCCGCCCCAGACCTTGGACCCGATGATCGCGCCCGTCTTATCCAAACTTTTGACGACGGATGGGTGTCATCGGCTGGTCCTGTTGTCGAGGCATTTGAACAAGCGTTTGCGGATCACGTTGGATTAGCCCACGCAACGGCCACAAGTTCGGGAACTGCGGCATTGCACCTGGCCTTGCACATGTTGGACTTGAAGCCCGGAGACGCTGTGATCGCGCCTACTCTGACATTCATCGGAGGAGTGGCCCCCATTGTGCACGCGGGAGCCCGCCCTCTTTTTGTGGATAGCTCTCCCGACGACTGGAATCTGGACCCCGCCCTGTTGGACGCAGCGTTTGCAAAAGCCCGTAC  AGAAGGTCTTACAGTGCGTGCCATCGTCCCCGCGGATCTTTATGGGCAAGGCTGTGATATTGGTGCAATCGGGGCAGTGGCTGCCTCCCACGGTGTACCTGTGATCTTGGACTCCGCAGAAGCAGTTGGCGCAATGGTGGGAGGTCGCCACGCCGGTCACGGAGCCTCAGCCGCGGGCTTCTCTTTTAACGGTAATAAAATTATCACTACCGGGGGCGGTGGTATGCTGGCATCGGACGACGGAGCCCTGATTGCTCGTGCTCGTTCTCTGGCGGCGGCAGCACGCATCCCTGCCGTGCACTATGAGCACGCTGAGGTTGGGTTTAATTATCGTATGTTATCTTTATCAGCTGCATTGGGATTAAGCCAGTTACCGACCTTGGAGGCCAAGGTAAATCGCCGTCGCGCAATCTTTGACCATTATCGTGGGCGCCTGGGGGGTCGTTATGGGGTAGACTTCGCACCAGAGGCTCCTGGGCGTCGTCATACCCGCTGGCTTAGCGTCATGGTCATGGACCGCGAAATTACTGGCGTCTCGCCTGAGCGTTTACGCCTTGCGCTGGCAGCCGCCGACATTGAAGCGCGTCCCGTTTGGAAGCCGATGCACTTGCAACCGGCGTTCCGTGATGCTCCAGTTCTGCAAAATGGCGTAGCAGAAACGCTTTTCGCCGGGGGATTATGCCTGCCTAGTGGCTCAAGTCTGACTGGCGCTGACATTGACCGTGTTTGTGATGTCATCGAAGCTACCTTGGACGCTTGAGCTAGCGGAGCTCAAGAAGGAGATATACCATGAGTCGTCACGGTCCCACCTTCGGGTTAATCGGCACTGGAGGATTTGCGCGCGAGGTCATGCCCGTAGCTCGCGCGTTTCTTCGTCATCACCCCGATTTAGGGATTGAGCCCGGTCGTATTGTCTTCGTAGATCGTGAAGCCGGAGCCCCCGTTGGCGGGGTTCCGGTACTTTCGGAAGCAGAATTTTTAGACCTTGACGGCGACCGCCATTTTTCCCTTGCCATTGGCGACGGAGCAGTGCGTCAGACGATCGCGGCTCGCCTTGAGGCAGCGGGATGCCGCCCATTATCACTGCGCGCAGACAATGTTCTTTTGCCTGATGACTTAGACTGTGGCCCAGGCGCTCTGTTCGCTCCATTTTCTATGGTGACTGCCGACGCACGTATCGGACGTCAATTCCAATGCAACCTTTACTCCTACGTCGCACATGACTGTGTGATCGGCGATTACGTTACTTTGGCTCCCCGTGTCTGTTTAAACGGAAACGTAGTAGTGGAAGATTTCGCTTATGTTGGTACAGGGGCAGTGATCCGCCAGGGTACTCCAGATAAACCGCTGGTGTTGGGCCGTGGATGCGTGATCGGGATGGGGGCAGTAGTCACTAAAGATGTAGCGCCGGGAGTAACGGTAGTCGGTAATCCGGCACGCCCCTTGGAGGCTCAATGAGATATCGCGGCCGCAAGAAGGAGATATACCATGGTCCGTTCAAAACGCTCGGTGGGAGTTTTCACGGCCACTCGCGCAGAGTATGGGTTATTACGCCCGCTGCTTGCCGCGCTGGATCGTTCCGCCACTTTGACTCCGCGCTTGATCGTTAGTGGGACTCATCTTAGCGATCGTCACGGAGGCACATTAACGGAGATCGAAGCAGACGGGCGCACCCCATCTGCCTGCGTCCCCGTGCTGTTAGCTGATGATTCCGGACGCTCCGTTGCTGCCGACATGGCGGCAGTGCTGGCCGGAGCAGCAGAAGCGTATCGCACTCTGGAACTTGAGGCCGTGATCATCCTGGGTGACCGTACTGAAGCATTGGCCGCAGCAGCAGCCGCTGTGCCTCTGGCCCTGCCTATTGTGCACCTGGAAGGTGGACATCGCACAGCCGGCGCTGTAGACGATGCCATTCGTCACGCGATTAGCAAATTGGCCGCGCTGCACTTTACTGGGGCGGAACCTTATCGTCAGCGTCTGTTACAGATGGGTGAAGCCCCAGAGCGCGTGTTTACAGTCGGATCGACCGGGGTGGACAATTTAGAAGCCTTCGGTCGCAAAACAGCCGCCGAAGCTAGTGCCATTACAGGATTGGATCTGCCCGAAGGGTTCGTGTTGGCCACGTTCCATCCGGAGACACTTGCTGCGACACCCGCGGGG  GATCAAGTGGCTGCATTTATTGCGGGATTAGAGGGTGCTGGAGACCGTACTTTGTTAATCACCCTTCCCAACGCCGACGTAGGATCAGGGACAGTACGCGCCGCTCTTGAACGCTTTGCTGCACGCGCACCTGACCGTGTCCGCCTTGTGGCATCGTTAGGAGCAAAGGGGTACGCCGCTGCCTTGACTGCATGCGCAGCTGTCGTCGGCAATTCAAGTAGTGGGATGATTGAAGCACCAGCCGCAGGAGTTCCTACTGTTAATGTCGGTGACCGCCAAGAAGGACGCTTACGCGCGCCCTCCGTGATCGACTGCGCGTTGTCCCCAGACGCAATCGCCTCCGCGTTGCGTCGCGCGTTGGACCCGGACTTTCGCCGTATGGCCAAGGCTCAACCCCCTATGTTCGGGGACGGACATGCCGGCCAACGCATTGTAGACATCTTGGAAACAACCGATTTCGCCGGACTTGCGCGCAAGCCCTTTATTGACTTGCCCATGCGTTGACAATTGCTGCAGAAGAAGGAGATATACCATGACGGAGCGTGTTTTCTTCATCGCGGAGGCAGGTGTCAACCATAATGGGGACTTAGACCTGGCACTTCGCTTAGTGGACGTAGCTCATCGCGCCGGTGCAGACGCTGTAAAGTTTCAGACCTTTACCACCGATCGTGTAATTGCAGTGGATGCACCGCTTGCTGAGTACCAAAAAGCCAACGCAGGAGCACCGTCAGCGGTGGAGATGCTGAAAGGATTCGAACTTCCACGTGCTGATTTTGCCCGCATCGCCGACCATTGTCGTGCTATTGGCATTGAGTTTATGTCATCACCCTTCGACTTGGAGTCAGCGGCTTTCCTGGCCGGGATTGGGATGACTAAATTCAAACTTGCCTCGGGCGAGATTACTCACGCTCCATTTGTCCGTGGAGTGGCTCGCTTGGCAGCTGACGCTGGTGGGAGCGTGATCTTGTCAACCGGAATGTCTACCTTGGATGAGGTCGCGGGCGCGGTCGGCTGGATCGAGGGTGAAGGTTGTCGTGATTTGACAATCCTTCATTGCGTGTCAAATTACCCTGCTCCCGCAAAGGACGCGAACTTACGCGCCATGGACACGCTTCGCGAAGCCTTCGGTCATCCTGTGGGATGGTCCGACCACACATTAGGTGACGCAGTGTCGTTAGCCGCCGTAGCCCGCGGCGCCCGTGTCGTTGAAAAACATTTTACCCTTGACAACGCACTTGCAGGTCCCGATCATGCAATGTCCATGGACCCCGACGGGTTAGCACGTTTGATTGCGGGCCTTCGTACAGTTGAGGCGTCACTTGGCGACGGGATTAAACGCCCAGTGGAAGCAGAACGTGAAATCATGACAGTCGCCCGCCGTTCGTTATTTGCAGCGCGTGACATCGCCGCAGGTGCGGTGGTGACTGAAGCAGACTTAATCGCATTGCGCCCCGGCGATGGCATTAGCGCGGCTCGCCATGCTGACGTAGTTGGTCGCACAGCCGCACGTGCTTTACCCGCAGGGACCAAGTTAGCGCCGACTGACTTGGTATGAAACTAGTACTTAAGAAGAAGGAGATATACCATGATTGATGGCCGCCGCGTATTAGGGATTGTTCCAGCACGTGCAGGTTCACAAGGCTTACCCGGTAAGAACCTGCGTTCGCTTGCTGGGCGCCCTATGACCGCTTGGACTTTAGACGCTGGCTTAGCAAGTGAAACATTGGATCACCTGGTAGTATCCACCGATGACCCAGGGGTAGCAGAACTTGCCCAGCGCGCGGGGGTTCCGGTAGTCGATCGCCCTCCTGGTCTTGCTGGCCCTCAGGCGTCGGTAATGGATGCCATCGCGCATGTGCTTCAGACGTTGGATCAGGATTGGGATTATGTGGTGCTTTTACAACCCACTTCCCCCCTTCGTACTCAAGTTGACATCGACGCGGCAGTAGCGTTATGTCACGACCGTGACGCACCGGCCGTTATTGGCGTGTCCCCCATGTTAAAACCAGCATCTTTTTATGGTCAGGTCGACCCTGATGGTACGTTCCATCGTGCAGCCGCCGCAACCGGCGAACGTGTCGTTGTTAATGGAGCATTATACGTGGGCCGCCCGGACTCAATGTTAGCAACACGTGGCTTCGTAGGCCCAGGGACCTTAGCATACTTGATGCGCCCAGAGTGCGGGTGGGACGTGGATACCGCTTTTGATTTCGCGGTTTGCGAGGCTCTGTGGCCTCACGTACGTGGTGAGTCTAGTCGCCCATTGGATAGCGTCCGCCGTTGAGAATTCCTCGGCCTCTAGAACTCGAGGCCTA |
| *C. botulinum neuB,* codon optimised for *E. coli* (5’-3’) | CCATATGAAAGACTTCTACATCGGAAAGAAAAAGGTGGGCGAGAAGTCGTCAATCTTTATTATCGCAGAAATTGGAGTAAATCATAATGGAAACATCGACCTGGCCTATAAGCTGATCGATTTGGCATCGAAGGCCGGTGCCGATGCTGTTAAATTTCAAATGTTCTATCCTGAGAAACTGTGTTCAGAGATTTACCGCAAAGATGAAATCGAAATGTTGAAAAAATATGTTTTATCTTTTGAAAATATGAAGAAATTACGCGACTATACATACTCCCTGGGGTTAGAGTTCATCGTCACGCCCTTTGACTTTAAGAGCTTAGATGACATCATCAATCTGGACTGTAGCGCCATTAAAGTCGCATCAGGGGAGCTTACGCATATCCCGTTTATCAAGAAGGCCGCCAGCTATAATAAGCCCCTTATCATCTCTACAGGCGCTTCGAACCTGTCCGACGTCGAACGTGCAGTCATTGCCATTAAGTCAGTGGCTGACGAAAAAATCTCAATTCTTCATTGCGTTTCTACTTACCCGAGTCCAGACGAATGTCTTAACCTTCGCGCATTGAAGACTCTGGAAGTTGCGTTTAATGACTGTATCATCGGTTTCTCTGATCATTCGCTGGGTTATACAGCCTCAAATCTTGCTGTCGCCTTAGGCGCGAGTATTATCGAGAAGCACATCACCTTAGACAAGAACCTGAAGGGTCCTGACCACAAAGCATCCGCCAGTCAAGGCGAGTTCGTCAACATGGTTTATGACATTCGCAAAGCGGAGAAAATGTTAGGCGATGGCTATAAAAAACCCCAAGCCTGTGAGGAAATCATTGGTCGTAGCATCGTGGCGGCCCGTGACTTGGAAGCAGGGGAGGTGTTACGCAAAGAGGATATCGATTACAAACGTCCTGGGTGGGGGATTCGTCCGTATGACGAGAATAAGATTATTGGTGCTCATTTGCAGAAGAGTATCAAAAAGGATGACATTCTTCAGTTGGATTATTTCATGCGTAAAATTGGGGACCAAAACGGTCGTTGAAATTCTAGACTCGAGAGCTCGCGGCCGCAGAATTC |
| *M. humiferrea MOHU_20790*, codon optimized for *E. coli* (5’-3’) | GAGCTCAGATCTGCTAGCCCAATTGCAGATCTAGGATCCGGATGTGAGCGGATAACAATTACGAGCTTCATGCACAGTGAAATCATGAAAAATTTATTGGCTTTGTGAGCGGATAACAATTATAATATGTGGAGGGGAATTGTGAGCGGATAACAATTCCCCTGTAGAAATAATTTTGTTTAACTTTAATAAGGAGATATACCATGGGCATGGGCTGGCATGATGACAAGTTAGGTGAAATTTGGGCGTTGCCGGGTGAGTCGCTTAAGCTGGCCCTGCCGCGTATGGATCGTGCAGGTTTGCAGGTCCTGCTGGTGGGCGACGCCGAGCGCCACCTGTTCGGGATCATCACCGACGGGGACATCCGTCGCGCCTTACTTCGCGGCGAATCGTTGGACGTACCTGTGGAACAGGTTATGCAGGCGCAACCGAAAGTCTTATCAGCGGGTGTTTCGTTAGACGTCGCTCGTCGCCTTATGTTAAATCATAACATCCGTCACATTCCGCTTGTTAATAATGAGCACCAAGTAGTCGATCTTCTGTTATGGATCGACCTTTTTGGCTCAAAGGTCGAGGCGCGTCCAGAACCCGTCGTCATTATGGCGGGTGGTAAGGGAACTCGTCTGGACCCCTTCACCAAGATCCTTCCCAAACCAATGATCCCCTTGGGAGATAAGCCTATTGTGGAAGTCCTGATGGACCGCTTCTATGACCATGGGTTTTCACAGTTTATCTTGTCTGTTGGGTACAAAGCCGAAGTAGTAAAGTTATACTTTAACGATAGTAATGGGCGCCCATACAAAGTCTCTTTCGTTCAGGAGGATGAACCCTTAGGAACTGCTGGCGCCTTAGGGCTGCTGCGCCAACAGTTGCAGCGCACATTCCTTGTCACGAACTGCGACGTTATCATCGAGATGAACTACGGAGAGCTTCTGCGCTACCATCACGAAAAAGGGAACGCATTGACGATCGTTGGTGCTTTGCGTGATTTCACTATTCCATACGGTGTTTTACGCACTGAAGCAGGTGAGTTCCATCAGATCGAAGAGAAGCCTTCGTTTCACTTCCTGGTAAATATCGGTTTGTACGTTCTTGAACCGGAAGTCCTGGAGGGGCTGGAAAATGGGTCGTTCATTCACATGACCGATCTTATCATGGCCGCTAAGGACAAAGGATTGCGTGTTGGTGTATATCCTCACCATGGGCGCTGGTTTGACATTGGGCAGTGGGACGAATACCGTCAGACCTTACGTGCCTTCGAGGGCTTGGTCTGAATTCTAGAGCGGCCGC |
| *G. kaustophilus neuB,* codon optimised for *C. crescentus* (5’-3’) | CATATGTCCAAGACGTTCATCATCGCGGAAGCGGGCGTGAACCACAACGGCTCCCTCGACCTCGCCTTTCAACTCGTCGATGCCGCCGTGGAGGCGGGCGCGGACGCCATCAAGTTTCAGACCTTCAAGACGGAACATCTGGTGACCAAGTCCGCCCAGCAAGCGGAGTACCAAAAGAAGAATATGGGGAAGTCGTCGAGCCAGTACGAGATGCTCAAGAAGCTGGAGCTGTCCTACGCGGATTTTAAGAAGCTCAAGCAGTACTGCGATGAGAAGGGCATCATGTTTCTCTCGACGCCGTTCGACCTCGAGTCCGTGGACTTCCTCATCCAGGAGCTGAAGCTCAATGTCATCAAGATCCCCTCGGGGGAGATCACCAATGCGCCCTACCTCCACAAGATCGCGCTCCACGGGGTGAATGTGATCCTCAGCACCGGCATGGCGACCCGCGATGAGATCCATCATGCGCTGGCCTTCCTCGCCTATGGGTTCGCGAACAAGCGGGATGTCAGCTTTGATAAGGCCAAGCGGTTCTACCAGACCAACGAGGCGAAGATGCTGCTGCAGGAAAAGGTCAGCATCCTCCACTGTACCACGGAATACCCGGCCCCGTATGAGGACGTCCATCTCAACGCGATGGATGACATGAAGGAAGAGTTCAATCTGTCGATCGGGCTGTCCGACCATACGGAGGGCATCGTCGTGCCCATCGCCGCGGTCGCCAAGGGGGCGAAGATCATCGAAAAGCATTTCACGCTGGACAAGATCCTGCCGGGCCCGGACCACAAGGCCAGCCTGGAGCCCAACGAACTCAAGGAGATGATCCAATCCATCCGCATCATCGAGAAGGCGCTCGGGGAAAAGCAGAAGCGGCCCACGCAGATCGAACTGAAGAACAAGGAAGTCGCGCGTAAGTCCCTGGTGGCCGCCAAGTCCATCAAGAAGGGCGAAGTCTTTACGTTTGACAATCTCACCGTGAAGCGTCCGGGCACCGGCATCGAGCCCTACTATTATTGGGATTACATCGGGCAAAAGGCGCAAAAGGACTATGAGGAAGATGAAGTCATCACCTGAATTCTAGACTCGAGAGCTCGCGGCCGCAGAA |
